# Predicting Personalised Therapeutic Combinations in Non-Small Cell Lung Cancer Using In Silico Modelling

**DOI:** 10.1101/2025.01.07.631497

**Authors:** Matthew A. Clarke, Charlie George Barker, Ashley Nicholls, Matt P. Handler, Lisa Pickard, Amna Shah, David Walter, Etienne De Braekeleer, Udai Banerji, Jyoti Choudhary, Saif Ahmed, Ultan McDermott, Gregory J. Hannon, Jasmin Fisher

## Abstract

The disease burden from non-small cell lung cancer (NSCLC) adenocarcinoma is substantial, with around a million new cases diagnosed globally each year, and a 5-year survival rate of less than 20%. A lack of therapeutic options personalized to individual patient genetics, and the targeted therapies that exist quickly succumbing to resistance, leads to high variation in survival. Patient stratification combined with greater personalisation of therapies have the potential to improve outcomes, however, the wide variation in mutations found in NSCLC adenocarcinoma patients mean that experimentally determining suitable treatment combinations is time-consuming and expensive. Here we present an *in silico* model encompassing tumour intrinsic key oncogenic signalling pathways, including EGFR, AKT, JAK/STAT and WNT for efficiently predicting rational drug-drug and drug-radiotherapy combination therapies in NSCLC. Using this model, we simulate diverse genetic profiles and test over 10,000 therapeutic combinations to identify optimal strategies to overcome resistance mechanisms specific to genetic profiles and p53 status. Our *in silico* model reproduces drug additivity experiments, predicts radio-sensitising genes validated in a CRISPR screen and identifies 53BP1 as a potential drug target that improves the therapeutic window during radiotherapy, as well as potential to use ATM inhibitors to overcome p53 loss-of-function driven radiotherapy resistance. We further use the *in silico* model to identify a 19-gene signature to stratify patients most likely to benefit from radiotherapy and validated this using TCGA data. These results further demonstrate the utility of *in silico* mechanistic modelling and present a bespoke computational resource for large-scale screening of personalised therapies applied to NSCLC.

## Introduction

Lung cancer is the leading cause of cancer-related deaths worldwide, resulting in an estimated 1.8 million deaths annually. Non-small cell lung cancer (NSCLC) adenocarcinoma accounts for ∼85% of lung cancer cases and has a 5-year survival rate of 19%^1^. Treatment options have advanced substantially over the last 10 years, with new surgical approaches, chemotherapeutic drugs and better targeted radiotherapy, with patients now often treated with several of these in combination^1,2^. However, overall survival remains low^3^, in part because current approaches do not account for the diversity of oncogenic mutations driving NSCLC^4^. Furthermore, in cases where targeted therapy is applicable, resistance is inevitable, with the majority of patients treated with first-generation EGFR inhibitors developing resistance after 6-9 months of treatment^1,5–7^. A lack of understanding of how tumour genetics affect response to therapy, and the mechanisms underlying resistance, limit our ability to rationally design treatments that pre-empt and overcome resistance, and personalise therapeutics to maximise individual patient outcomes^8^. Radiotherapy delivers a high dose of radiation to cancer cells to cause the rapid accumulation of double-stranded breaks (DSBs)^9,10^. If unrepaired, these DSBs trigger cell death^11^, while healthy cells are better able to repair these breaks and survive. Radiotherapy has a significantly lower treatment-associated mortality than surgery^12^, although the efficacy of surgery is higher than that of radiotherapy in terms of overall survival^13^, owing in part to adaptation of the tumour and evolution of radio resistance-associated mutations^14^. Extending the benefits of radiotherapy to more patients and improving efficacy to reduce the chance of resistance emerging requires better understanding of how variation in treatment response is driven by differences in driver mutations. This will allow tailoring of radiotherapy to patient biology by combining it with radio-sensitising treatments.

Recent advances in modelling patient biology as executable computer programs are now beginning to uncover the principles underlying how the genetics of a cancer contributes to its behaviour and the potential mechanisms that give rise to phenotypes such as therapeutic resistance^15–21^. Here, we present a bespoke executable model of NSCLC adenocarcinoma oncogenic signalling that covers key pathways including the cell cycle, apoptosis cascade and DNA damage response (DDR). This model can mechanistically link oncogenic genetics and signalling, by representing biological processes as nodes within a qualitative network, with functions determining the activation of biological components^22^. The model is ‘executed’ by allowing nodes to interact with each other dynamically, such that the stable state of the network represents the overall cellular phenotype^23^. The model can be used to test combinations of mutations and inhibitors involving proteins that are not yet druggable, allowing identification of potential drug targets for future research and development. It can also be interrogated in greater depth than real systems as every variable and interaction in the model can be controlled individually, allowing for more complete exploration of the mechanisms underpinning observations. Moreover, this mechanistic understanding of how therapy interacts with the oncogenic mutations allows us to identify the genetic determinants of responses to therapy.

Here we use this *in silico* model to explore 136 possible radio-sensitising treatments across a panel of 7 cancer cell lines and explore the effect of specific mutations on treatment sensitivity. Once the model successfully recapitulates previously published sensitivities of different NSCLC adenocarcinoma cell lines to combination drug treatment and radiotherapy, we use the model to screen ∼66,000 combinations to find the optimal therapeutic strategy for a given genotype. We exploit the interpretable nature of our i*n silico* model to understand why these sensitivities occur and how specific mutations determine the NSCLC adenocarcinoma responses to therapy. We then perform *in silico* screens for perturbations that can sensitise cancer cells to radiotherapy while protecting healthy cells, as well as overcome p53-driven radiotherapy resistance. Overall, our platform is a powerful tool for understanding therapeutic responses to treatment in NSCLC. Harnessing the power of executable models will provide a more complete understanding of how genetics contributes to oncogenic signalling and enable therapy to be tailored to the genetics underlying the cancer, improving treatment outcomes for patients.

## Results

### An executable network map of KRAS-, EGFR- and MET- driven NSCLC adenocarcinoma

To understand the contributions of oncogenic mutations in NSCLC adenocarcinoma to drug and radiotherapy response, we built an executable qualitative network model using the BioModelAnalyzer (BMA) tool (https://biomodelanalyzer.org, see Methods) following the workflow outlined in Figure 1A. The model describes the core pathways covering the most common driver mutations in NSCLC patients^24^, including EGFR, AKT, JAK/STAT and WNT pathways, as well as commonly overexpressed genes such as MYC and NRF2. We modelled how these mutations drive changes in cell cycle and apoptosis. To model radiotherapy responses, we include the DNA damage repair (DDR) pathways, homologous recombination (HR) and Non-Homologous End Joining (NHEJ).

**Figure 1.**
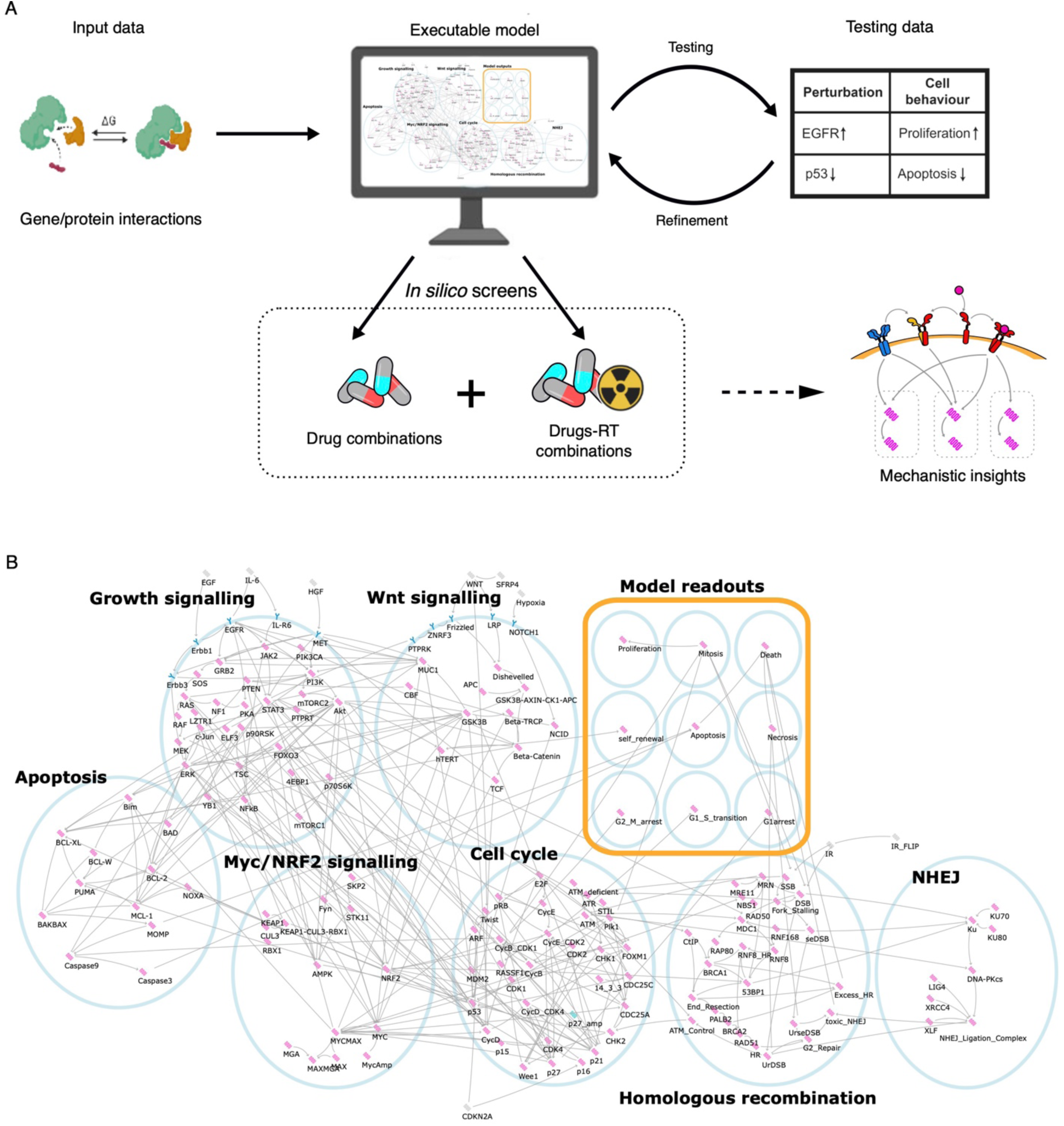
Outline of the workflow and executable NSCLC adenocarcinoma model. **A)** Schematic showing the workflow used to construct, test, and analyse the model. **B)** A network visualising the architecture of the executable model as seen in the BMA tool. Different subnetworks are enclosed by a blue circle, model inputs are coloured grey, receptors are coloured blue and cytosolic proteins are coloured pink. Edges represent regulatory interactions between proteins and are pointed for activating edges (→) and blunt for inhibiting (⊣). The phenotypic readouts are shown enclosed in a yellow box.

An extensive literature survey of 171 research papers identified interactions between genes and proteins in NSCLC adenocarcinoma resulting in a model with 160 nodes and 360 edges describing activating regulatory relationships or inhibiting relationships between biological components (Supplementary Table S1). To provide context to these interactions, each node has a corresponding target function defining its activity (Shown in Figure 1B, Supplementary Table S1, S2 and S3). The value describing the activity of nodes is discrete and ranges between 4 levels (0 to 3). A level of 0 indicates loss-of-function and 3 indicates gain-of-function.

### The NSCLC adenocarcinoma network model reproduces experimental data

To train the model, we collated a list of 26 publications reporting the behaviour of NSCLC adenocarcinoma cell lines under different experimental conditions, separate from and not used to build the model. These publications describe the behaviours (e.g., growth rates, levels of apoptosis, protein levels) of different cell lines under perturbation by drugs and/or radiation. Cell lines were modelled by setting the nodes that represent genes with known driver mutations to the appropriate constant value (0 for loss-of-function and 3 for gain-of-function). Cell line mutation data were drawn from Cell Model Passports^25^ and our own experiments (see Methods, Supplementary Table S4 and Supplementary Figure S1). We then used the model to find the stable states of the system, which we compared to the phenotypes measured in the experiments. Comparison of the simulated behaviours with the published observations was used to iteratively improve the target functions of the nodes within the model until the model recapitulates the behaviours observed in the published experiments (see Methods, Supplementary Table S5). We also modelled the sensitivity of six cell lines to single drug treatment with and without radiotherapy (HCC44, NCIH1373, NCIH23, NCIH358, NCIH1792, SW1573) and compared to the published literature, to test the DNA-damage repair section of the model (Supplementary Figure S2, S3). We examined common targeted therapies including EGFR^26,27^, MEK, MET,^28^ and DNA-PK inhibitors^29,30^ and further verified that the model reproduced the effects of knock-out experiments on key proteins in oncogenic processes, including KRAS^31–34^, MUC1^35^, NRF2^36,37^, JAK2^38^, RAD51^39^, ATM^40^, RNF8^41^, NOTCH1^42^ and TP53^32^. When comparing the model behaviour to these published experiments, we see an accuracy of 91% (Supplementary Table S5, See Methods).

To identify treatments that maximise the difference, for the same dose, between the effect on cancer and healthy cells (maximise the therapeutic window) we also simulated healthy lung cells. We assume that the signalling processes in these cells are the same as in lung cancer except without driving oncogenic mutations. We tested this assumption by comparing to published experiments conducted using the immortalised epithelial lung cell lines HBEC3 and BEAS-2B^32,43^ (Supplementary Table S4). Having reproduced existing experimental data, we next used the model to systematically predict the effect of unseen combinations of knock-outs in our panel of NSCLC adenocarcinoma cell lines.

### Targeted monotherapies inhibiting Erbb1, RAS and STAT3 have an effective but narrow therapeutic window

To simulate personalised therapies with single targeted drugs, we systematically set the value of 95 druggable (see Methods, Supplementary Table S6 and Supplementary Figure S3) nodes in the model to 0 in six NSCLC adenocarcinoma cell lines (SW1573, NCIH1792, HCC827, PC9, A549 and NCIH1993) chosen for a range of driver mutations to reflect the genetic heterogeneity of NSCLC (See Supplementary Figure S1). By setting the value of nodes to a constant value of 0, we mimic the effect of inhibitors that could be used as targeted therapy. Overall, we observed that cancer cells are more resistant to targeted therapy than healthy cells when considering both cell proliferation and death levels (Supplementary Figure S2). Perturbations that are effective at killing cancer cell lines are those that directly target proteins downstream of activated oncogenes specific to those cell lines (e.g., EGFR in HCC827 and PC9 or KRAS in NCIH1993, NCIH1792 and A549), however, these treatments are also lethal to healthy cells. Inhibition of STAT3 kills cells in the A549 and NCIH1993 cell lines (Supplementary Figure S2C), a pattern observed with all other potential treatments demonstrating how small differences in mutational profile can lead to total resistance to therapy.

When identifying potential cytostatic treatments, we identify more potential drug targets. This includes inhibiting MYC, which limits oncogenic growth to the basal growth rate of healthy cells while also reducing apoptotic signalling in all cell lines profiled apart from A549. In general, we see that healthy cell lines are more susceptible to treatment via the inhibition of single proteins than cancer cells (Supplementary Figure S2C). This underlines the importance of using combination therapies to maximise the perturbation space to find cancer-specific treatment options with a good therapeutic window.

### NRF2 inhibition is predicted to protect healthy cells while 53BP1 inhibition sensitise cancer cells to radiotherapy

We next sought to assess combining targeted therapy with radiotherapy to see whether this would improve predicted treatment efficacy. We therefore used the model to assess the radio-sensitising effect of gene knockouts and compared the predictions to an *in vitro* CRISPR screen of radiation sensitive and resistant lung adenocarcinoma cell lines (HCC44, NCIH1373, NCIH23, NCIH358, NCIH1792, SW1573). We set one node at a time to a constant value of 0 to mimic a knock-out, in addition to cell line-specific mutations (Supplementary Table S4), and compared this effect with or without radiation, modelled through the value of the ‘IR’ node, (Supplementary Figure S3). The effect of IR is modelled through the induction of single-strand breaks (SSB), double-strand breaks, and replication-induced or single-ended double strand breaks (seDSB)^44^. We modelled the impact of SSB on fork stalling and collapse, and the subsequent generation of seDSB^45^ as well as the downstream repair of DSB and seDSB by homologous recombination and non-homologous end-joining^46^, with unrepaired breaks and toxic chromosome fusions leading to increased cell death. We define an effective radio-sensitising knockout as one that leads to more cell death as part of a combination compared to either the gene knockout or radiotherapy alone. We term the additive difference between cell death in the combination treatment and the most effective of these two monotherapies as ‘radio-sensitisation’ (see Figure 2A).

**Figure 2.**
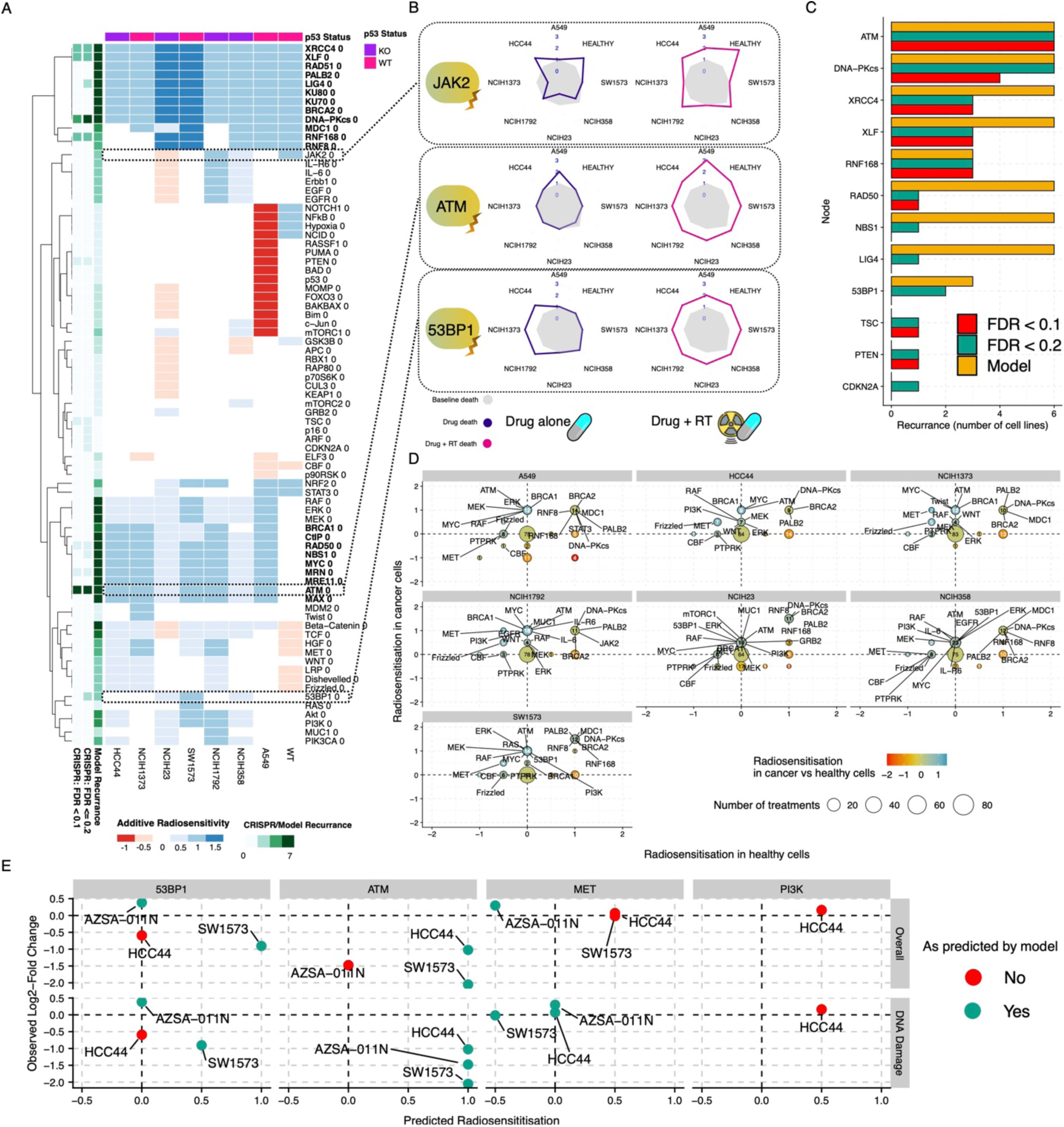
Systematic screen of perturbations on the *in silico* model reveals inhibition of ATM and 53BP1 as radio-sensitising agents. A) A heatmap showing predicted radio-sensitising effect *in silico* knockouts (rows, where the node value is set to 0), on NSCLC adenocarcinoma cell lines (columns). Radio-sensitisation is the increase (blue) or decrease (red) in cell death in the combination of IR and knock-out compared to the most effective of IR or knock-out applied alone. Side panels (green) show the number of cell line backgrounds in which the treatment was radio-sensitising in the model or the CRISPR screen (at FDR ≤ 0.2 and < 0.1) where we count a recurrence if radiosensitivity is observed at this threshold for any timepoint in the screen. Knockouts that confer broad radiosensitivity are shown in bold. **B)** Radar plots showing the levels of death after drug treatments (for ATM, 53BP1 and JAK2) (dark blue) and after drug treatment and irradiation (magenta) compared to the baseline (grey). Specific drug combinations that are broadly radio-sensitising across backgrounds are outlined in black in the heatmap in panel A. **C)** Number of cell lines in which knockouts were predicted to be effective in: the network model (yellow), CRISPR screen with FDR ≤ 0.2 (green), or CRISPR screen with FDR < 0.1 (red). Note: p16 and p14ARF knockouts were both observed to be effective in the CRISPR screen. These are both controlled by the CDKN2A node in the model and so are grouped under that node. **D)** X-axis change in healthy cell radio-sensitivity. Y-axis relative change in cancer cell radio-sensitivity. Blue is more radio-sensitisation of cancer than healthy; red is more radio-sensitisation of healthy than cancer. Point size is the number of knockouts that have this effect. **E)** Observed Log2-Fold-Change (y) vs predicted radio-sensitisation for genes in focussed CRISPR screen (x). Values shown are for maximum timepoint. Colour is blue for validation (FDR<0.1 for cases where model predicted sensitisation), red otherwise. AZSA-011N cells are healthy cell organoids. Any genes for which knockout was found to be essential at all timepoints were removed. Top row is predicted radio-sensitisation measured by change on overall cell death (measured by node ‘Death’), bottom row is predicted radio-sensitisation measured by DNA-Damage specific cell death (measured by node ‘Necrosis’).

The model predicts broad classes of radio-sensitising knockouts. Some knockouts are radio-sensitising in both healthy and cancer cells; primarily inhibition of the NHEJ pathway, as expected^47^. Other known treatments such as ATM inhibition^48^ are radio-sensitising across all cancer cell lines, but we also see cell line-specific radio-sensitising knockouts. For example, 53BP1 inhibition is particularly additive in SW1573 (Figure 2A-B). Such cell line specific behaviour demonstrates the difficulty in maintaining an effective therapeutic window across diverse patient mutations, and underlines the requirement for tailored therapeutic approaches to diverse mutational profiles.

We compared the model results to an *in vitro* CRISPR screen in the six NSCLC adenocarcinoma lines treated with radiotherapy (HCC44, NCIH1373, NCIH23, NCIH358, NCIH1792, SW1573) (Figure 2C). The model correctly predicts those knockouts that led to radio-sensitisation in the *in vitro* CRISPR screen in 9 out of 12 cases (Figure 2C).

The model suggests that the radio-sensitising effects of knockouts outside of the NHEJ pathway are highly specific to mutational context, as we saw for many monotherapies (Figure 2A). In addition, when radiotherapy is applied, the model predicts similar levels of death in NSCLC adenocarcinoma and healthy cells (Supplementary Figure S3B-C). However, the degree to which a drug radio-sensitises a cell (the difference between the combination of IR and knock-out compared to the most effective of either IR or knock-out applied alone) is often higher in the cancer cells than the healthy cells. In particular, there is a subset of cases where all the death in healthy cells is driven by IR, but the majority of death in the cancer cells is driven by the combined effects therapy and IR. This suggests that by reducing the IR dose we could reduce the death in the healthy cells more than in the cancer cells and widen the therapeutic window. Combined with the personalised nature of radio-sensitisation, this prompted us to query the model for knockouts that lead to the largest change in radio-sensitivity.

Figure 2D shows the predicted additive radio-sensitisation of cancer cells versus healthy cells. Many treatments are predicted to radio-sensitise both cancer and healthy cells, for example those involved in DDR pathways. In the case of radio-sensitisation of cancer cells specifically, we predict that inhibition of MYC is particularly effective and specific. This is partly due to the essential role of MYC in proliferation^49^, but also because of the regulation of NBS1 by MYC, which, as part of the MRN complex, plays a key role in coordinating the response to DNA damage. Conversely, we predict that loss of members of the PTP family (e.g., PTPRT), leads to reduction in radio-sensitivity in healthy cells, protecting them from toxicity, while having no effect on many of the cancer cell lines. This is due to an increase in JAK/STAT signalling, with a concomitant increase in MCL-W that suppresses apoptosis, and NRF2, which aids in repair of radiation induced DNA damage. This has less effect in cancer cells as they have already up-regulated NRF2. We examined death from DNA damage (through the node ‘Necrosis’) specifically as this may play a greater role in response to different radiotherapy doses than apoptotic death (Supplementary Figure S3D). We predict that inhibition of NRF2 is particularly sensitising to DNA-damage associated death in cancer cells, due to reduction in DNA damage repair^50^. Equally, suppression of the KEAP1/CUL3/RBX1 complex leads to an increase in NRF2 in healthy cells, protecting them, without aiding cancer cells, as these over-express NRF2 via mutant KRAS.

To validate this, we test a selection of these genes, MYC, 53BP1, MET, PI3K and ATM in a focussed CRISPR screen in cancer cell lines (SW1573, A549, HCC44) and healthy cell organoids (AZSA- 011N). Of these, MYC was unable to be tested as it proved essential in all cell lines. 53BP1 sensitised cancer cells and not healthy cells, in line with the model’s predictions, although it sensitised both SW1573 and A549 relative to healthy cells, where we predicted it would be effective only in A549. ATM sensitised all cancer cell lines, as predicted, as well as healthy cells, which the model did not predict. MET did not show the expected sensitisation of the cancer cells (Figure 2E (top)), nor the protection of healthy cells (AZSA-011N). However, in our model we treat Apoptosis and DNA-damage associated death (Necrosis) separately. When examining the latter, we find that the model’s predictions align well with observation in all genes and cell lines, other than for HCC44 in the case of 53BP1 and PI3K (Figure 2E (bottom)), although 53BP1 may be essential for HCC44 (essentiality was detected at early time points but not later timepoints).

### ATM inhibition overcomes radio-resistance conferred by TP53 deficiency

As a key regulator of DNA damage response which is often mutated in cancer, we wanted to explore the association between TP53 mutational status and radiotherapy response in cancer cell lines. Accordingly, we simulated the cell lines from our radio-sensitisation screen with TP53 activated in cell lines where it has a loss-of-function mutation and inactivated in cell lines where it is competent (Supplementary Table S4). The model predicts that TP53 deficiency reduces death from radiation alone in most cell lines (Figure 3A, Supplementary Figure S4). However, when combined with a radio-sensitising knock-out, there are many potential combinations that are predicted to induce death in cancer cell lines regardless of TP53 activation, overcoming the radio-resistance due to TP53 loss-of-function (orange points on dotted line, Figure 3A).

**Figure 3.**
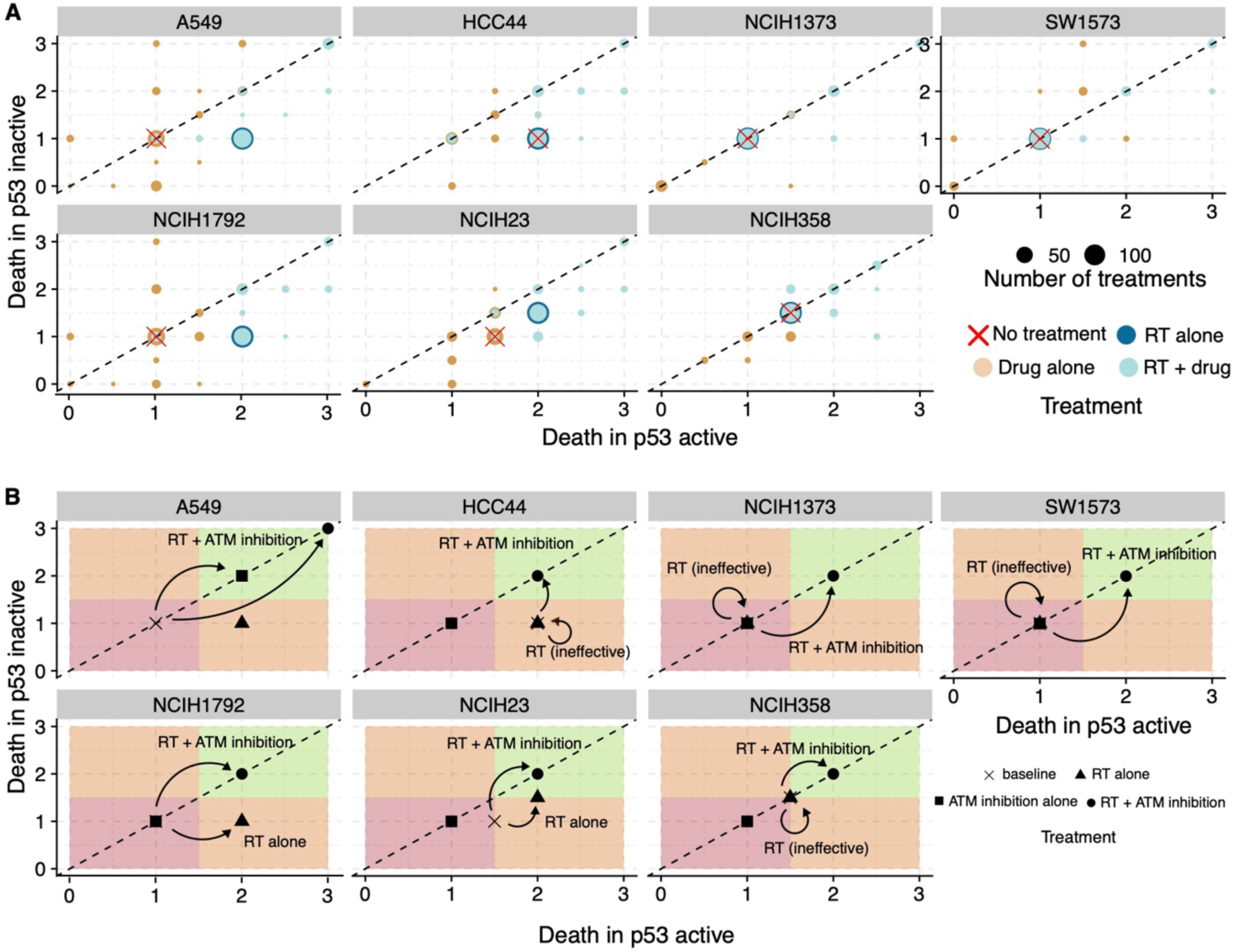
Radio-sensitising knock-outs increase death in a manner that is agnostic to TP53 status. **A)** Levels of death predicted by the models in cell lines that have been modified to have TP53 active (x-axis) and TP53 inactive (y-axis). Colours denote the type of treatment (dark blue for drug treatment alone, light blue for radiotherapy alone and orange for radiotherapy and drug treatment combined) and the size denotes the frequency of treatments in that position. Knock-outs below the dotted line are less effective in TP53 inactive cases then in TP53 active, knock-outs above the dotted line are more effective in TP53-inactive than active. **B)** Death in TP53 deficient (y-axis) and TP53 (x-axis) proficient cell lines showing the differences of radiotherapy alone and radiotherapy with ATM knock-out. Quadrants of the plots are coloured to denote desirable locations for therapeutic outcomes. Green indicates high cancer death regardless of TP53 mutational status, red indicates low death regardless of TP53 mutational status and orange indicates that knock-out efficacy is dependent on TP53 mutational status. Arrows show how in the case of ATM knock-out, the level of cell death with IR moves from below the dotted line i.e., less effective in the TP53 inactive case, to the dotted line, indicating that IR + ATM inhibition overcomes and neutralises the effect of TP53 inactivation.

ATM knock-out is an example of a radio-sensitising perturbation that is effective across all cancer cell lines, irrespective of TP53 mutation status (Figure 3B). Therefore, the model predicts that there is benefit in using radio-sensitising agents targeting those proteins identified in our i*n silico* knock-outs to improve the effect of radiotherapy for patients with a TP53 mutation. We experimentally validated these results by modifying SW1573 and NCIH1373 cell lines *in vitro* to flip the mutational status of TP53 (Figure 3C and Supplementary Figure S4A) Indeed, ATM sensitises cancer cells regardless of p53 status in all cancer cell lines for which this was tested in the focussed CRISPR screen (A549 and SW1573), as predicted.

### Predicted radio-sensitising targets stratify patient response to radiotherapy

Following our systematic *in silico* screen of radio-sensitising knockouts, we observed that there was a subset of genes that affect radiation sensitivity agnostic to mutational background. These 19 genes are marked in bold on Figure 2 and include members of the DNA damage repair pathways (ATM, MDC1, XRCC6, XRCC4 etc) as well as transcriptional regulators (MYC and MAX). We hypothesised that these candidate genes could be used to stratify patients as either radioresistant or radiosensitive. To explore this, we used the TCGA database (https://www.cancer.gov/tcga) of clinical data to see whether patients with low levels of gene expression (as measured by RNA-seq) or copy number variation (CNV) for our predicted radiosensitive genes survive longer after treatment with radiotherapy. We considered either of these properties to be analogous to the activity of the node within our network model. Patients with both low copy numbers (bottom 5%, n= 76) and of low RNA expression (bottom 10%, n=15) for at least one radio-sensitising gene, compared to all patients, were determined to be radiosensitive (Figure S5A, n=175).

When instead considering RNA expression and CNV separately, we found that with either definition of activity of a gene we see a lower hazard ratio (longer survival) for patients predicted to be radiosensitive after receiving radiotherapy (Supplementary Figure S5B-C, Cox proportional hazards regression; HR= 0.4299, P=0.06 for expression and HR= 0.6421, P=0.08 for CNV). When using both CNV and RNA to determine a consensus subset (see Methods) of radiosensitive patients we also see a significant difference in survival (Figure 4A, HR= 0.5842, P=0.03), suggesting that patients with a naturally lower activity of these genes have increased survival. When looking at patients that have not received radiotherapy (n=705, Supplementary Figure S5D) we see no significance (Supplementary Figure S5E-G), indicating that these genes are stratifying patients only in the presence of radiation (Figure 4B), consistent with our model’s prediction of these targets as potential radio-sensitising agents.

**Figure 4.**
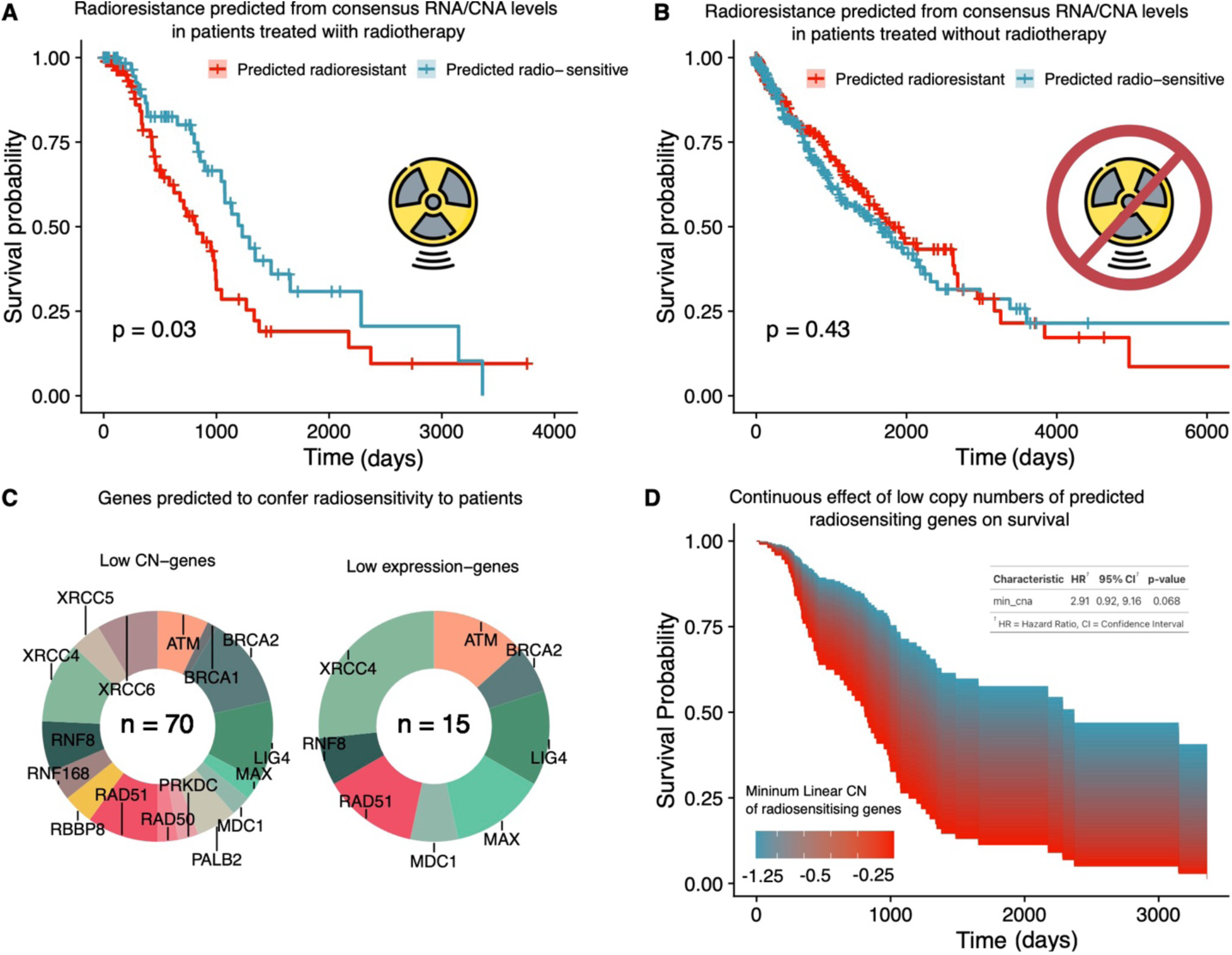
*In silico* predictions stratify radiation-treated patients into radiosensitive and radioresistant strata. **A, B)** Kaplan-Meier survival curves showing the differences in survival between predicted radioresistant patients (red) and radiosensitive patients (blue) for patients treated with radiation (B) and without radiation (C). The x-axis indicates the time (days) since diagnosis and y-axis indicates the survival probability. **C)** Pie-charts showing the genes with the lowest copy number (left) or expression (right) for each patient predicted to be radiosensitive. **D)** Continuously coloured survival area plots showing the effect of the minimum copy number of radio-sensitising genes on survival. The x-axis indicates the time (days) since diagnosis and y-axis indicates the survival probability. Colour indicates the minimum copy number of genes within our predicted set, with blue indicating lower copy number.

Lastly, we investigated whether there was a continuous relationship between the lowest level of a radio-sensitising gene in a patient and their survival following treatment with radiotherapy. This assumes that the level of radio-sensitisation in a patient is rate-limited by each of our model-predicted radio-sensitising agents. For each patient, we calculated the lowest linear copy number of each of the 19 genes (Figure 4C) and then used this continuous variable to predict survival in patients exposed to radiotherapy. The survival area plot for this analysis is shown in Figure 4D, showing significant association between the minimum CNV level of the radio-sensitising genes and survival (Cox proportional hazards regression; HR=2.91, P=0.07). This association does not exist for survival among patients not treated with radiotherapy (HR=1.19, P=0.6), indicating this relationship only exists in the presence of radiation. Collectively, these findings demonstrate that low CNA and low RNA expression of our predicted radio-sensitising genes are significant predictors of improved survival in patients receiving radiotherapy. However, our modelling and statistical analysis of clinical outcomes also shows that many patients are radioresistant and so this necessitates broader choices of therapeutic options.

### *In silico* screens identify MYC and ATR inhibitor as a cancer-specific treatment combination

To find further treatment combinations that are effective specifically on cancer cells we looked at different combinations of targeted therapies that could increase the difference in death between healthy and cancer cells. Combination treatments are also likely to reduce resistance compared to monotherapy^51,52^. We performed *in silico* screens of pairwise combinations of 95 druggable targets in the model for SW1573, NCIH1792, HCC827, PC9, A549 and NCIH1993 cell lines (see Methods). We isolated the treatment combinations that were predicted to have an additive effect on overall survival (proliferation - death) when compared to the most effective single treatment (see Methods and Supplementary Figure S6). This was measured for survival rather than death because combination therapy seeks to limit oncogenic growth as well as induce apoptosis, as opposed to radiotherapy which has the primary aim of causing apoptosis or DNA-damage associated death^53^. We then clustered these treatments into those that have a similar effect on the cell lines to highlight patterns of response and profiles of sensitivity. This produced 24 different subclusters, labelled *I - XXIV*, in order of their efficacy vs the most effective single treatment (see Methods). We see that clusters *I, II* and *IV* are the only broad-spectrum treatments that have a wide, additive effect across all cell lines tested (Figure 5).

**Figure 5.**
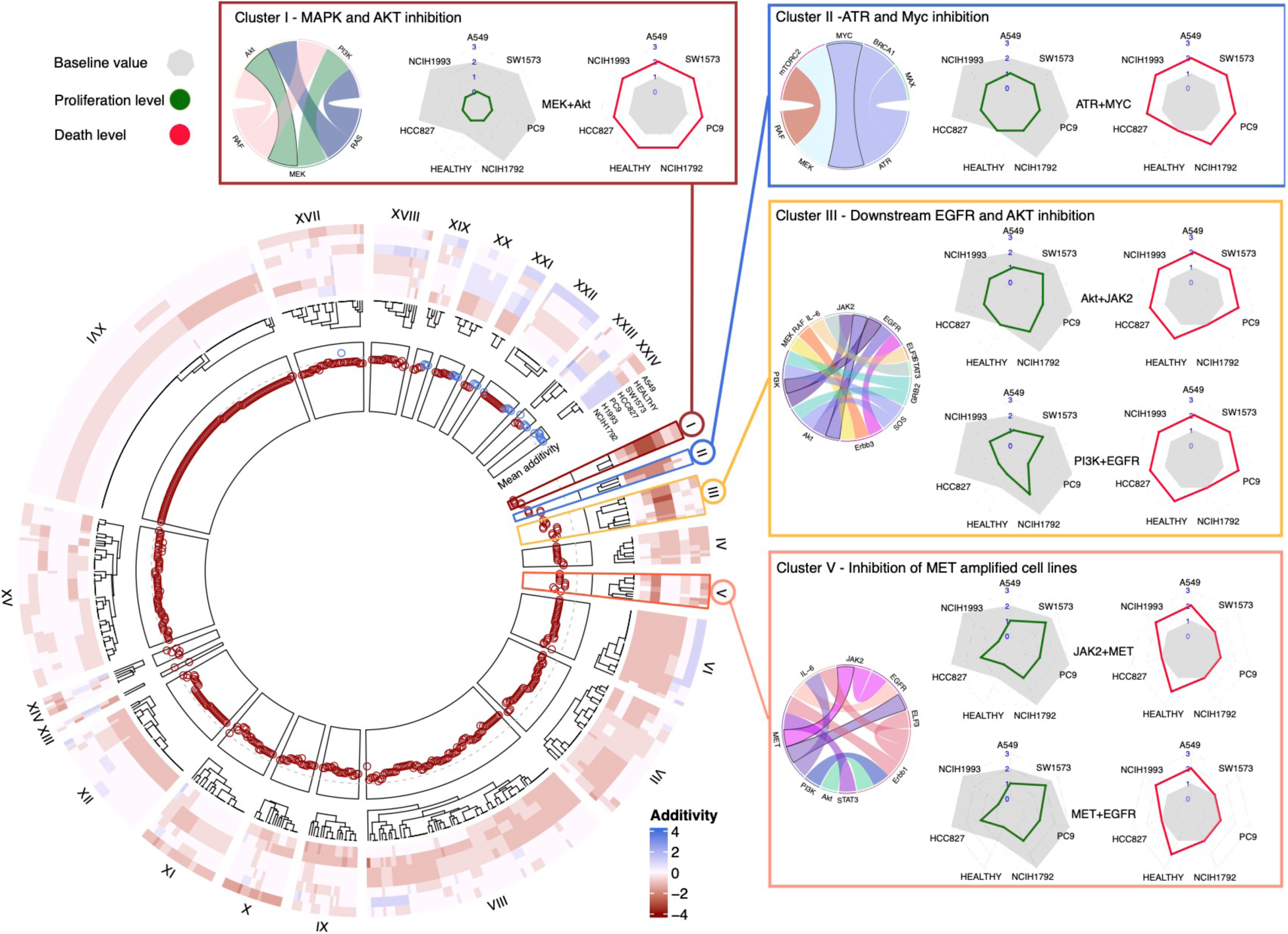
Systematic *in silico* combination screen on NSCLC adenocarcinoma network model. Circular heat map showing the additive effect of drug combinations on selected NSCLC adenocarcinoma cell lines. Columns (spokes of the wheel) are the drug combinations and have been clustered into groups (I-XXIV) of drugs that have a similar additive effect. Red indicates those combinations that have a greater additive effect, while blue represents those drugs with antagonistic effects (the drugs cancel one another out). The plot in the inner wheel of the circle shows the mean additive effect of a drug across all cell lines and is red for those drug combinations with overall additive effect and blue for those antagonising one another. Clusters I, II, III and V, and the most effective combinations within, them are highlighted. Circular chord diagrams illustrate the various drug combinations within a cluster and radar plots show the absolute levels of proliferation (green) and death (red) after drug treatment when compared to the baseline (grey). These are shown for specific drug combinations that are outlined in black in the chord diagrams. ⍰

We next compared our simulated results to experimental data generated by Nair and colleagues measuring the additive score of various drug combinations in SW1573, NCIH1792, HCC827, PC9, A549 and NCIH1993 (Supplementary Figure S7)^54^. We see that the data shows significant concordance with our predictions across all cell lines tested. Comparing Higher than Single Agent effect (HSA) to our predictions of additivity, we see moderate, but significant correlation (Kendall Tau = 0.16; p-value = 8.176e-11) across all drug combination predictions. Given the large proportion of intermediate or no additivity predicted by our model and measured by Nair and colleagues, we also looked at correlation of purely additive combinations as predicted by our model (those shown in Figure 2). In these groups we see stronger correlation (Kendall Tau = 0.27; p-value = 1.7e-07,). Correlations of the most additive treatments in Clusters *I - V* show stronger correlation still (Kendall Tau = 0.37; p-value = 2.3e-07). For reference, the average Pearson’s correlation across experimental replicates in GDSC^55^, CTD^56^ and PRISM^57^ drug screening platforms is ∼0.30^58^. We also find that the model’s predictions can recapitulate the sensitivities of different cell lines to specific drugs (See Supplementary Figure S8 and Supplementary Notes). These results indicate that our predictions correlate with published data and correlation improves when discriminating between more additive therapies.

According to the simulations, the most effective group of drug combinations by average additivity across cell lines are those targeting AKT and MAPK pathways (Cluster I). These have a uniform effect across cancer cell lines, increasing death from the untreated (baseline) level while also causing growth arrest (Figure 5). This combination does not discriminate between oncogenic or healthy cells, so although it is a reliable means of treating a wide range of mutational profiles in cancer, it is also likely cytotoxic to healthy tissue. Cluster II discriminates between healthy cells and tumorigenic cells and is composed of ATR and MYC inhibition. It limits growth to be on par (proliferation = 1) with healthy cells, while increasing death only in cancer cell lines. The remaining highly additive clusters are those dealing with RTKs mutated in specific cancer cell lines. For example, Cluster *III* is additive in cell lines with *EGFR* gain-of-function mutations, while Cluster *V* is additive in NCIH1993, a cell line with a *MET* mutation. These clusters illustrate the importance of cell line-specific key genetic determinants of response that govern the most effective and additive combinations needed for efficacious treatment. However, given drugs within Cluster II are broadly effective regardless of mutational profile, while sparing healthy cells, we decided to explore further the mechanism of action of this line of therapy.

### Combined MYC and ATR inhibition is predicted to limit cell growth and kill cancer cells through double stranded breaks

The model predicts that inhibiting MYC limits proliferation and causes cell death when combined with ATR inhibition in cancer cell lines without affecting healthy cells. This is in line with our earlier prediction of the effect of MYC inhibition in combination with radiotherapy. ATR inhibition alone is predicted to have little impact on all cell lines tested, whereas MYC and ATR inhibitions are predicted to increases cell death in cancer cell lines more than in healthy cells (Figure 6A). To assess the effect of these perturbations on the entire model state, we used Principal Component Analysis (PCA) to visualise the entire state space of the model following treatment with ATR and MYC inhibitors (Figure 6B), seeing that the treatment predicts to affect healthy cells (bottom left) less, while leading to a large difference in cancer cell lines (top). The loadings for different model variables are also plotted as vectors to highlight those that contribute to the separation in coordinate space of the PCA and represent variables that are changing mostly after perturbation. We predict that following treatment there is an increase in p15, G2/M arrest and toxic NHEJ, which is responsible for the increase in cell death. The variables and the regulatory interactions that exist between them are plotted in Figure 6C. This suggests how ATR and MYC inhibitions combine to limit oncogenic signalling, as well as the induction of DNA damage-associated death in cell line PC9 (the remaining cell lines in Supplementary Figure S9). MYC inhibition is predicted to disrupt the oncogenic operation of the cell cycle, while also inhibiting apoptosis and homologous recombination. Inhibiting ATR could then increase single-ended double strand breaks (seDSB) by preventing the cell from responding properly to replication stress, which is higher in cancer cells due to their higher rate of proliferation. MYC inhibition prevents proper repair of seDSB owing to the loss of MRN and thus ATM activation, leading to toxic NHEJ^46^. Healthy cells are also predicted to be less affected by the inhibition of MYC, which is more highly activated in every cancer cell line (Figure 5C). As a result, inhibition of MYC and ATR is predicted to have the greatest therapeutic window of all the treatments we simulate and could be a promising therapeutic candidate applicable to a wider variety of mutational backgrounds.

**Figure 6.**
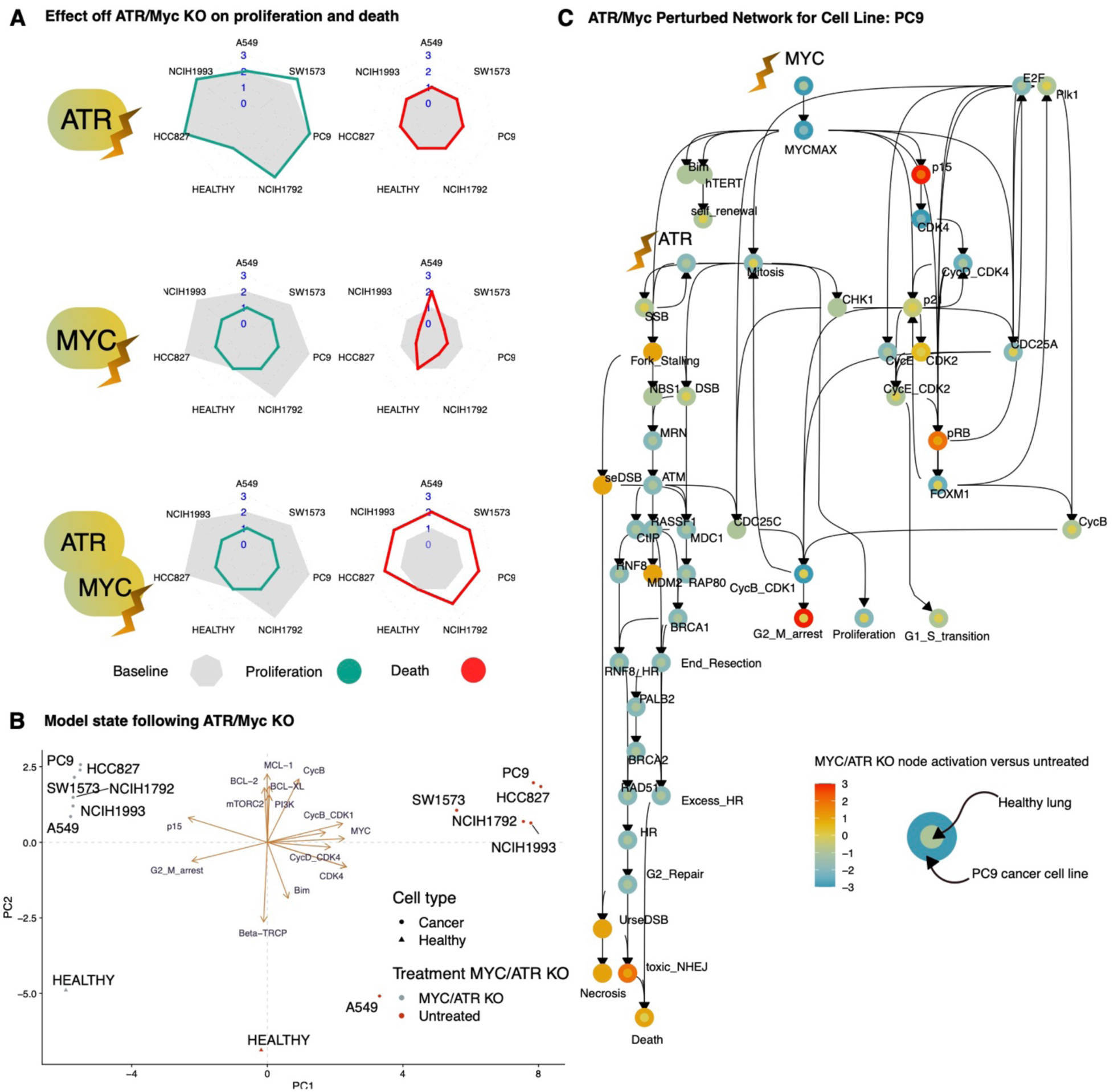
Mechanistic analysis of combined ATR and MYC inhibition illustrates their induction of DNA damage associated death. **A)** Radar plots showing the absolute levels of proliferation (green) and apoptosis (red) after MYC inhibition, ATR inhibition and combined MYC and ATR inhibition. The baseline growth and proliferation are shown in grey. **B)** PCA showing the model states of different cell lines following treatment with MYC/ATR inhibition. Cancer cell lines are plotted as circles and healthy cells (lacking oncogenic mutations) are triangles. Untreated cells are in red and treated cells are in red. Vectors are also included in the PCA to visualise the loadings of different variables and how they contribute to the separation of *in silico* samples in 2D space. **C)** Subnetwork of the model, covering the proteins and variables that changed after treatment of MYC/ATR inhibitors in PC9 (a cancer cell line). Colour of the node illustrates the difference between ATR/MYC inhibition vs untreated (blue for a relative decrease in activation and red for an increase) in PC9 (outer circle) and the difference in healthy lung cells (inner circle).

As of writing, MYC inhibitors are in early stage trials^59–61^, however, other specific drug combinations that we predict to be additive are composed of clinically approved drugs. Unlike MYC and ATR inhibition, these treatments are only effective in selective cell lines. To explore this effect more systematically, we investigated the key explanatory mutations in cell lines that drive responses to various groups of targeted therapy combinations that we predicted as being additive.

### EGFR and TP53 mutations explain additive response to combined MAPK and AKT inhibition

Given that treatment effectiveness varied with small changes in mutational profile, we aimed to find which mutations have the greatest explanatory power in drug response, as these could be used as biomarkers. Accordingly, we visualised drug additivity in the most effective combination clusters and aligned this with the mutations used to model the cell lines (Figure 7A). We predict that EGFR is strongly associated with response to treatments in clusters *I, III* and *V*. For example, treatments in cluster *I* (MAPK/Akt inhibition) are most additive in PC9 and HCC827 (EGFR mutant cell lines). Similarly, treatments in Cluster *III* are most additive in cell lines that contain EGFR gain-of-function, while the opposite is true for Cluster *V.* Additionally, TP53 mutations are associated with response to clusters I, and III with the most additive responders to those clusters of combinations being NCIH1993, HCC827 and PC9 (Figure 7A).

**Figure 7.**
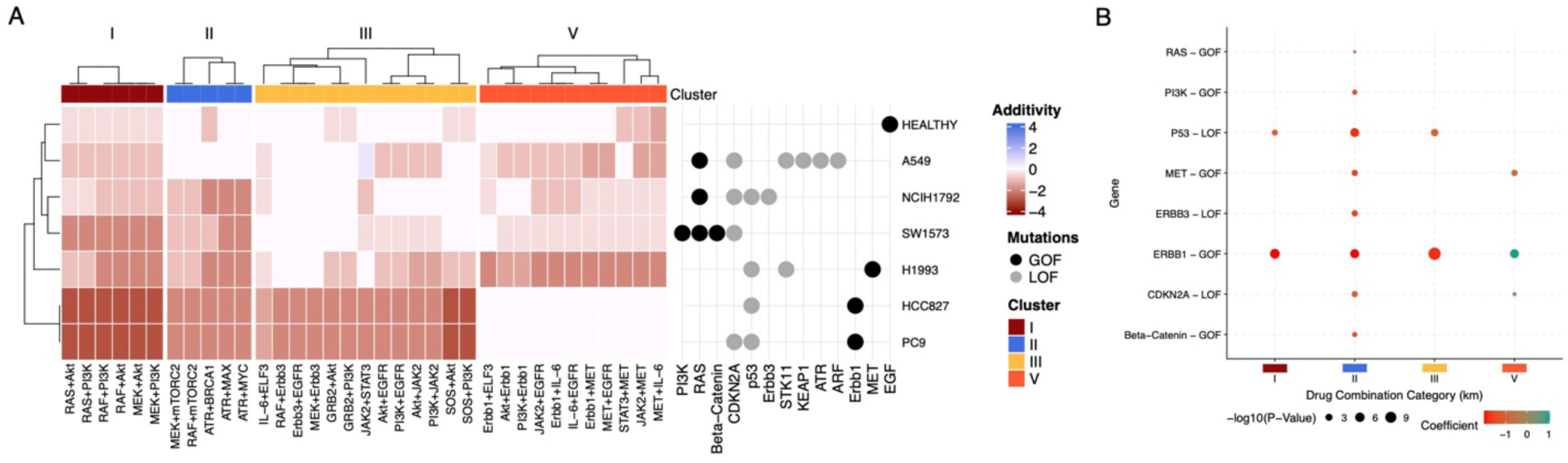
EGFR and TP53 mutations explain the response of cell lines to targeted combination therapy. **A)** Heatmap illustrating aligning the mutations of cell lines (right) with their response to the most effective drug combination clusters from Figure 5 (I in red, II in blue, III in yellow and V in orange). Columns of the heatmap indicate specific drug treatments and rows indicate cell lines. The colour in the heatmap refers to additivity with red indicating strongly additive behaviour and white indicating no additivity. The schematic (right) shows mutations as columns and cell lines as rows, with gain-of-function mutations for a cell line and loss-of-function (grey). **B)** Plot showing the coefficients of the linear model relating mutations present in a cell line (y-axis) to additive response against drug clusters (x-axis). Colour represents the value of the coefficient, such that blue indicates a mutation is associated with additive response to a cluster and red indicates that a mutation’s absence is associated with additive response to a cluster. Size of the dot represents the significance of the association.

We used a linear model to quantify the relationship between cell line mutations and their responses to combination treatments and find the most explanatory mutations for additive drug combinations responses in the different cell lines (See Methods). We found that TP53 and EGFR mutations are most broadly associated with an additive response to the tested combinations (Figure 3B). However, for Cluster V we see a negative coefficient for EGFR mutations, indicating that the absence of this mutation is required for additivity to be observed. Instead, a MET amplification (as is the case for NCIH1993) is associated with additivity in this cluster (Figure 7B). These results suggest which mutations determine response in different mutational profiles, and highlight EGFR, TP53 and MET as key for patient stratification. Such insight is key to bridging therapeutic responses and oncogenic mutations, allowing us to stratify patients to find the most suitable course of treatment for a given situation.

## Discussion

Matching the variety of mutational profiles exhibited by NSCLC adenocarcinoma with the wide range of existing and potential therapeutic options presents a huge combinatorial challenge. Here, we showcase an *in silico* platform for executable modelling to explore this immense therapeutic landscape. By using the broad and extensive prior knowledge from published experimental research, we constructed an executable model that can explore treatment options in an efficient and mechanistically interpretable fashion. Using this platform, we assessed ∼66,000 predicted combinations of targeted therapy and radiotherapy, finding the most suitable and effective treatment options, specific to given genotypes. We also investigated how mutations combine and determine response to therapy, and the mechanisms that underpin this, allowing for testable hypotheses to expand our understanding of oncogenic signalling, and demonstrate a novel method for stratification of patient sensitivity to radiotherapy using clinical data based on targets identified by the *in silico* model.

Finding that healthy cells are more sensitive to single-agent treatment than cancer cells in our *in silico* modelling and motivated by the need for combination treatments to prevent resistance^51^ we explored combinations of drugs with one another and with radiotherapy. We find that the most effective cluster of drug combinations (Cluster *I*) consists of combined MAPK/PI3K inhibition. AKT and MAPK are downstream processors of all growth pathways in the *in silico* model and most oncogene mutations found in lung cancer target members of these pathways or upstream regulators (e.g., KRAS, PIK3CA, EGFR, MET, ALK). MAPK/AKT inhibitors are often used as positive controls for synergy experiments and have previously been flagged as highly additive in lung cancer^54,62^. However, healthy cells also rely on these pathways and so their inhibition can cause side-effects^63^. We were able to identify other combinations that specifically targeted cancer cells owing to mutations in the NSCLC adenocarcinoma cell lines. Specifically, we found that EGFR is an important determinant for the additive response to several potential combinations. We also find that absence of EGFR mutation is as an important determinant of response to MET inhibitor combinations than the presence of a MET mutation itself. In general, we show that the mutational status of receptors is important in determining additivity to targeted therapy, and so a patient’s mutational status of EGFR, MET among others are particularly informative in determining personalised therapeutic options.

Most combinations have a similar effect on healthy and cancer cells. However, for both radiotherapy and combinations of targeted therapy, we predict that MYC has strong potential in targeting cancer cells without affecting healthy cells. MYC is frequently overexpressed in cancer, owing to its central position in the hourglass topology of pathways governing regulation and effect^64^. This importance makes resistance to MYC-targeting drugs extremely difficult^65^. This structure is reflected in our model and means that for the heterogeneity of cancer mutational profiles, they broadly produce a higher activation of MYC and its downstream targets^65^. This puts MYC in a unique place as a promising therapeutic target with inhibition well tolerated in healthy tissue^66^. However, despite advances in developing inhibitors targeting MYC^67,68^, none are currently clinically approved for treatment. Our work underlines the exciting prospect of MYC as a therapeutic target and goes further in suggesting ATR as a parallel target, offering the option to further exploit weaknesses in cancer cells DNA damage repair. We show that by targeting cancer cell reliance on MYC overexpression and the dependence of DSB repair on MYC via the MRN complex, one can produce therapeutics that are robust and broadly effective against varied mutational profiles. However, this depends on how sensitive DDR pathways are to MYC inhibition as MYC inhibitors such as Peptomyc^61^ and OMO-103^69^ are likely to be less effective compared to the full knockout simulated by the *in silico* model. Our model suggests that the effect of MYC inhibition on cancer cells is due, in part, to its regulation of DSB repair via the MRN complex. Prior work in cell lines found that DDR was robust to MYC downregulation^70^. Our analysis shows that a clinical signature (derived from *in silico* simulations) including MYC and its binding partner MAX act as a predictor of radiosensitivity (Figure 4A). Together these results suggest that the link between MYC and radio-resistance should be investigated further.

Radiotherapy remains a main component of the standard of care for NSCLC adenocarcinoma treatment^1^. A key factor in response to radiotherapy is TP53, with mutations disabling its activation, either in the TP53 gene or upstream, frequently seen across all tissues and often associated with therapy resistance^71,72^. The exact effect on radio-sensitisation is contentious, with TP53 loss-of-function often leading to radio-resistance^73,74^, but contributing both to radio-sensitivity and radio-resistance in different contexts and tissues^75–77^. We predict that TP53 loss-of-function mutations in NSCLC adenocarcinoma are broadly radio-protective, yet in the presence of specific mutational profiles can be radio-sensitising. A subset of potential treatments is radio-sensitising despite TP53 mutation, and some are completely invariant to it (Figure 3B). This suggests that radio-resistance can be overcome, or pre-empted, by identifying the correct treatment and offers hope that this may hold with a broader range of resistance mechanisms. Due to the increased toxicity of radio-sensitising treatments, we predict it will be important to stratify patients by TP53 mutation^78^ for drug-radiotherapy treatments. TP53 mutation occurs early in NSCLC adenocarcinoma^79^ and is strongly associated with metastatic seeding sub-clones. As metastases are the key driver of patient death^80^, treatments that target clones that are mutated in drivers of metastases are especially important. Our identification of radio-sensitising treatments that overcome TP53-driven radio-resistance satisfies a key part of this. We further use our model to identify a 19-gene signature that, by combining CNV and RNA-expression data, stratifies patients into radiosensitive and radioresistant groups. We suggest this can be used to better target radiotherapy treatment to those patients that are most likely to benefit from it in the clinic.

Radio-sensitising combination treatments are already being explored on a small scale, for example as part of the CONCORDE clinical trial^81^, and we demonstrate the potential of this approach with our large scale *in silico* and *in vitro* screens. We validate our *in silico* model predictions in a focussed CRISPR screen and find that the model prediction of DNA-damage related cell death is a good indicator of sensitisation. We were unable to test our predictions in cases where gene knock-out was lethal to cells without radiation, e.g., in the case of MYC. Such knock-outs are less well tolerated than intervention with drugs, as demonstrated by MYC inhibitors such as Peptomyc^61^ and OMO-103^69^ being well tolerated in clinical trials, and so further studies with inhibitors may prove fruitful. Calibrating contributions for targeted treatments and IR on different mechanisms of cell death (i.e., apoptosis vs DNA-damage associated cell death) is challenging. Indeed, we find that our model is more accurate when we output DNA-damage associated cell death and apoptosis separately. It is of especial note that inhibition of 53BP1 leads to radio-sensitisation in cancer cell lines but not in healthy cells. 53BP1 loss is a common resistance mutation to PARP inhibitors^16^, and thus this may suggest a collateral sensitivity that can be exploited for second line treatments. 53BP1 loss is a common resistance mutation to PARP inhibitors^16^, and thus this may suggest a collateral sensitivity that can be exploited for second line treatments.

Current experimental approaches to identify combinations of therapies involve performing global screens of perturbations on various cell models of NSCLC adenocarcinoma with differing mutational profiles. This can be done with drugs or CRISPR knockouts alongside ‘omics measurements, observing the survival of the treated cells to understand hidden mixtures of lethal targets^54,62,82^. For example, Jaaks *et al.* tested 2025 pairwise drug combinations *in vitro* using the Genomics of Drug Sensitivity in Cancer (GDSC) cell line screening platform to identify biomarkers of response to combination therapies^55,83^. While such screening strategies can yield extremely high throughput, identifying target hits and biomarkers of their response across a broad range of unrelated backgrounds, there are practical drawbacks of such approaches. For example, developing a mechanistic understanding is hampered by off-target effects in drugs and CRISPR guide RNAs^84^, which can lead to false positive^85^ and false negative results^86^. Additionally, these screens are time-consuming and expensive and cannot be performed on rarer genetic profiles easily, leading to results biased towards the most common cellular models.

Computational methods can circumvent some of these technical issues and provide complementary screening approaches. For example, Eduati *et al.* use perturbation screening data to train a network to uncover patient-specific mechanisms underpinning pancreatic cancer^87^. Their model was trained using perturbation screens of patient biopsies and tested pairwise combinations of 12 targetable nodes in the network. In our work, we build on this by using more coarse-grained, but large-scale modelling of biological processes, allowing for more targetable nodes (i.e., 95 druggable targets), and integrated radiotherapy. Our modelling approach also only require mutational data for individualisation of our generic model, rather than training on combinatorial perturbation screening on cancer biopsies, which are more expensive and invasive and have limited clinical applicability. Fröhlich *et al.* used a larger, pan-cancer mathematical model to predict effective drug combinations^88^. However, the complexity of the parameter space of these models makes them computationally intensive to run on novel mutational backgrounds and mathematical models with many parameters more prone to overfitting than coarse-grained executable biological models. Others have used phospho-proteomic changes^62^ and machine learning^89^ to predict drug sensitivity. However, while the accuracy of the method was high^62^, it is not clinically feasible currently to take phosphoprotein measurements of patients to determine drug response. Moreover, machine learning based approaches used to predict synthetic lethal targets lack the mechanistic interpretability of this method and often neglect radio-therapeutic combinations due to the relative lack of genome-wide data on radiotherapy sensitising agents. Our *in silico* model incorporates radio-resistance which is important to consider as it is already a common clinically used therapy.

In this study we demonstrate how executable modelling enables system-level experiments to be rapidly conducted. In addition, our *in silico* approach allows mechanistic transparency, providing insight into why treatments do and do not work and the biological underpinnings of mutational profile-specific sensitivities. Using mutational data, our executable models can be individualised, paving the way for ‘digital twins’ of tumours for testing hundreds of thousands of therapeutic options. We envisage that executable models have the potential to improve personalised therapy in NSCLC adenocarcinoma patients and help predict and combat the development of resistance to treatment.

## Methods

We performed all visualisation and data analysis using the programming language R, and data visualisation packages ggplot2, ComplexHeatmaps and Circularize. Figure 1A was created with BioRender.com.

### Qualitative networks

We represent the NSCLC adenocarcinoma signalling network as a discrete qualitative network^22^. This represents each gene, protein, or process as a node with multiple finite values of the range (0-3), where 0 represents very low activity e.g., due to a loss-of-function mutation, 1 represents normal behaviour, 2 elevated activity and 3 represents very high activity e.g due to a gain-of-function mutation. This was built, tested, and analysed using the open source webtool, BioModelAnalyzer (BMA, https://biomodelanalyzer.org) and associated command line tools (https://github.com/hallba/BioModelAnalyzer). The model and code for analysis is available at . Nodes in the network represent biological factors such as proteins or subunits of proteins. Input nodes represent growth factors, or in some cases amplification as with ATM. Output nodes are representing phenotypes such as death and proliferation. Regulatory interactions (both activating (→) and inhibitory (⊣)) are represented by edges between nodes. The level of activity of a node responds to the upstream levels of regulators of that node. This is determined by a mathematical function, unique to each node that integrates the levels of the upstream regulators to determine the target level of the target node at the next time-step, with the activity of each node changing by at most 1 unit per time step. This function is termed the target function. The default target function is the average of the activating upstream regulators, negated by the average of the inhibiting regulators (*avg(pos)-avg(neg)*). More complex functions are used as needed to recapitulate specific experimental behaviours (see below). These target functions along with corresponding references are described in Supplementary Table S2. A summary of the nodes included, and their HUGO gene symbol are summarised in Supplementary Table S3.

For a specified starting state of all nodes, the model will update node values synchronously, such that there will be one deterministic stable attractor of the network. This attractor can be a fixed-point attractor with one end state, or it can be a loop of states that the model cycles through, called a cyclic attractor. We test the model from all possible initial states, which allows for multiple attractors to be reachable, termed a bifurcation. In the case of a loop or bifurcation, the upper and lower bounds across all attractors are returned. This is done using the algorithm described in Cook *et al.*^90^. The algorithm for network simulation is found in Schaub *et al.*^22^. If there is no single attractor, we use the midpoint of the upper and lower bound as an approximation of the overall level of the node.

### Model construction and testing

Experimental evidence for each model edge is described in Supplementary Table S1. We manually curated a separate set of 26 papers describing the behaviour of different NSCLC adenocarcinoma cell lines under various perturbations and conditions. From these papers 69 separate experimental conditions were modelled and used to optimise the target functions of the model. This is described in Supplementary Table S2. Accuracy was calculated by taking the number of experiments recapitulated, versus the total number of experiments. An experiment was considered recapitulated if the predicted direction of change of the model is the same as that of the experiment.

### Mutational profiles for all cell lines were drawn from Cell Model Passports

To model cell lines we used mutational data from Van der Meer *et al.* as of 04/01/2023^25^. Mutations were modelled if they were listed as a driver or a deletion for a node in the model. If annotated as a gain of function the respective node was set to the maximum value, if a loss of function or a deletion the node was set to zero, except in the case of ATM. After the results of the first CRISPR screen, we observed that ATM knock-out was still effective in NCIH1373 and NCIH23 cells despite an ATM mutation observed in both, therefore we model this as a deficiency but not complete homozygous knockout of ATM. We focussed on annotated driver genes and complete deletions so as not to over-specify the model, and to focus on those changes that were likely to be consistent within the population of cell lines^91^.

### Mutational profiling of cell lines used in radiotherapy CRISPR screens

Mutational data was derived using publicly available datasets, principally DepMap and corroborated using western blotting (*e.g.*, TP53 status). Radiation sensitivity was assessed using colony forming assay, and longer-term cell viability assays (e.g., cell-titre glo >7day timepoint) and assessed against published literature.

### Modelling of combination and monotherapy

To find effective treatments, we inactivate (set target function to a minimum) or activate (set target function to maximum) all nodes representing druggable nodes in the network either singly or in a pair-wise combination, using the BMA Command Line tool BioCheckConsole. For each node in the network, we curated a list of drugs that could target nodes within the network, finding 95 candidate drugs using the druggable proteome in The Human Protein Atlas (https://www.proteinatlas.org/humanproteome/tissue/druggable; Accessed 04/08/2023). We evaluate these perturbations by identifying the stable state of the network for death and proliferation as outlined in the ‘Qualitative Networks’ section above. To have a single value to predict cell growth we take the difference between Proliferation and Death.

For combination therapy we simulated these drugs together and studied their interaction using a value we describe as additivity. This is analogous to the experimentally determined Higher than Single Agent effect (HSA) and describes the growth of cells after combined drug treatment relative to the most effective (minimum viability) of the individual drugs in the combination. Combination treatments were clustered using the k-means algorithm to find groups of therapies with similar levels of additivity across cell lines.

### Modelling of radiotherapy

To model the effects of radiotherapy, we increase the ‘Radiotherapy’ node to maximum. This leads to increases in the single and double strand break nodes. To model the ability of the cell to repair these insults, we allow these nodes to activate repair mechanisms, while also activating ‘UrDSB’ (Unrepaired DSB). If the repair mechanisms are active, as in healthy cells, this means that UrDSB is inhibited by the activity of those mechanisms to represent the initial insult due to radiotherapy being resolved without causing death, while any damage to the repair pathways will let DSB activate UrDSB and in turn leads to death. Unrepaired SSB are modelled as leading to single-ended or blunt ended DSB (seDSB and DSB) respectively but not leading to death by themselves.

We model radio-sensitisation as the increase in death, decrease in proliferation, or decrease in overall survival (proliferation - death) in the combination vs the most effective of the two possible monotherapies (knock-out alone or IR alone).

### CRISPR knockout of radioprotective and radio-sensitising genes

A CRISPR screen was performed using a pooled library across the 6 NSCLC, Cas9-expressing, lines using 1Gy given twice over 24 hours (i.e., 2Gy in 2 doses over 2 days). Cells were irradiated using an IBL 637 Irradiator (CS 137 gamma rays below sample). Cells were harvested from day 7 to day 31 (up until cells reached 10-12 doublings) and DNA was prepared for next generation sequencing. Hits were identified using MAGeCK RRA (version 0.5.8). Cut-offs of FDR<0.2 and FDR<0.1 were used to define significant depleted genes.

Due to the *in vitro* CRISPR screen considering all 20,000 human genes, when a knock-out was observed to have an effect in only a subset of the cell lines it was not possible to conclude whether this was due to a vulnerability contingent on a specific mutational profile, or the power of the screen. For this reason, we measure and compare the number of cell lines in which a knock-out was radio-sensitising in the CRISPR experiments vs the *in silico* model (i.e., recurrence).

A second screen was conducted with a focussed panel, across the SW1573, SW1573 *TP53*^-/-^, A549 and A549 *TP53*^-/-^ cell lines and AZSA-011N organoids. After 72 hours, we irradiated cell cultures with a single fraction of 4Gy. Post-irradiation, precisely after 1 hour, 24 hours, and 48 hours, irradiated and non-irradiated plates were fixed for immunofluorescence.

### Generation of organoids normal tissue organoids

Patient derived lung organoids (PDO) were isolated from adjacent normal tissues from consented NSCLC patients at Royal Papworth Hospital via the Royal Papworth Hospital Tissue Bank application ID T02612 / East of England - Cambridge East Research Ethics Committee reference 18/EE/0269.

### Cell culture

HCC-44-Cas9, SW1573-Cas9, H1792-Cas9, H1373-Cas9, H358-Cas9 and H23-Cas9 cell lines were maintained in RPMI 1640 (Gibco) supplemented with 10% FBS (Biowest), 1% GlutaMax (Gibco) and 1% pen/strep (Gibco). Cell lines were passaged every 3/4 days by dissociating to single cell using Accutase (Biowest).

### Genomic DNA isolation and Next Generation Sequencing (NGS)

Genomic DNA (gDNA) was isolated using the QIAamp DNA Blood Midi or Maxi Kit (Qiagen) according to manufacturer’s instruction. NGS libraries were prepared in a two-step process: the integrated lentiviral cassette containing the gRNA was amplified from gDNA. Forward primer 5’ ACACTCTTTCCCTACACGACGCTCTTCCGATCTCTTGTGGAAAGGACGAAACA 3’ with reverse primer 5’TGACTGGAGTTCAGACGTGTGCTCTTCCGATCTACCCAGACTGCTCATCGTC 3’ was used for this. PCR reactions with 5µg gDNA per well were set up using the Q5 Hot Start High-Fidelity 2× Master Mix (NEB #M0494) in a total volume of 50µl. PCR reactions were scaled accordingly to amplify the gRNAs at a coverage of at least 200-fold. The PCR products were then purified using QIAquick PCR Purification Kit (Qiagen #28106). Final NGS libraries were generated using 2.5ng of the purified 1st PCR product using the dual-indexing Illumina-compatible DNA HT Dual Index kit (Takara #R400661). 2nd PCR products were purified with AMPure XP beads (Beckman Coulter #A63881) at an 0.7 ratio. Purified 2nd step libraries were quantified using the Qubit dsDNA Quantification Assay Kit (ThermoFisher) and sequenced on NovaSeq6000 by PE50bp with a 30% PhiX spike-in.

### Genetic modification of p53 status of NSCLC cell lines

The SW1573 line underwent TP53 knockout using a standard CRISPR-based knockout approach to generate a polyclonal TP53 knockout population. Loss of TP53 was confirmed using PCR and western blotting.

### Stratification of patient data using results from model

Clinical and genomic data for lung adenocarcinoma (LUAD) and lung squamous cell carcinoma (LUSC) were obtained from The Cancer Genome Atlas (TCGA) Research Network (https://www.cancer.gov/tcga) and the cBioPortal platform^92,93^ via the RTCGA.clinical R package. Clinical data was categorized based on whether patients received radiotherapy or not, using the patients not receiving radiotherapy as a positive control. Molecular data included RNA expression and Capped relative linear copy-number (CN) values for each gene (from Affymetrix SNP6) and was extracted via the TCGAretriever R package. To identify patients with low expression or alterations in the genes of interest, the CNA and RNA datasets were combined for both LUAD and LUSC. We identified patients as radiosensitive if they had gene expression or CN values below specific quantile thresholds (5th percentile for CNA, 10th percentile for RNA) for at least one gene in our model-defined set of radio-sensitising genes. To associate patient strata with clinical outcomes we used the Cox proportional hazards regression model in R. To visualise the survival probability over time as a function of the lowest CN we used the R package contsurvplot^94^.

### Statistical analysis of combination drug sensitivity

For drug combination we use the dataset available from Nair *et al.* (2022) and take all the drug inhibitors they used and compare them to our predictions. Nair *et al.* use HSA and so we compared our value for additivity (see above). For statistical analysis, we visualise t test between HSAs of different treatments for different values of additivity that the model predicts (−3 through 1) and compare this to treatments predicted to be not additive. Also, to factor in the ordinal nature of the model’s prediction, we calculate overall correlation using Kendall’s tau test and bin experimentally measured HSA into 4 quartiles.

### Linear regression to identify genetic determinants of response

We separate the drug treatments into the clusters outlined above and train a linear model to relate drug response (measured by additivity) with the mutational profile of the cell lines. P values were adjusted for multiple hypothesis correction between each drug cluster using the method devised by Benjamini & Hochberg (1995)^95^.

## Acknowledgments

This work was supported by Cancer Research UK grant C17918/A28870 (G.J.H. and J.F.), CRUK Convergence Science Centre, Experimental Cancer Medicine Centre and NIHR Biomedical Research Centre (L.P., U.B. and J.C.), and National Institute for Health Research University College London Hospitals Biomedical Research Centre (J.F.).

