## Supplemental Materials for "Predicting Personalised Therapeutic Combinations in Non-Small Cell Lung Cancer Using In Silico Modelling"

### Supplementary Notes

To assess the predictions of the model, we compared our predictions of Additivity to those observed by Nair et al, which were not used to train the model. As of writing, MYC inhibitors are in early stage trials[1,2,3], however, other specific drug combinations that we predict to be additive are also tested in Nair et al.[4]. Here, we illustrate how the model can predict differential cell line sensitivities for the most effective drugs. Additive score, as predicted with the model, had a positive correlation with the HSA results of Nair et al. in 5 cases (Supplementary Figure S7A). We observe positive correlation between predicted additive score and HSA in all but one of the drug combinations (inhibition of AKT+MEK, EGFR+PI3K, JAK2+MET and EGFR/Erbb1+MET). This illustrates how the model can predict differential cell line sensitivities for the most effective drugs. JAK2+PI3K is the drug combination where the model prediction does not align with the experimental predictions. However, of these 6 combinations, JAK2+PI3K inhibition has the highest variance in Nair et al.'s results (Supplementary figure S7B) Nair et al.'s experimental results for additivity for JAK2 inhibition and PI3K inhibition are highly dependent on the specific drugs used to target either JAK2 or PI3K, for example drug Fedratinib and AT9283 have very different effects, despite both being inhibitors of JAK2, and as such the results are likely to be because of off-target effects of the drugs rather than through the direct inhibition of JAK2 or PI3K[4,5].

The model predicts AKT+MEK inhibition to be broadly effective across cancer cell lines. Indeed, it is a combination that is often used as a positive control for synergy experiments. The experimental data affirms this, aside from the cell line HCC827, which is predicted to be maximally additive (-3), but displays no additive inhibition in experimental data. This discrepancy conflicts with the similarity of mutational information for PC9 and HCC827, with both having oncogenic mutations in Erbb1 and p53 (Supplementary Figure S7G). Another study, Coker et al., measured synergy between these two drugs in HCC827 found strong synergy (Bliss score of 0.6, Supplementary Figure S7B)[6], implying high additivity also and thus agreeing with our predictions. From this we hypothesise that likely the strain of HCC827 that Nair et al. is using likely contains some uncharacterised differences that disrupt this synergy, otherwise it would behave similarly to PC9 as the simulations would suggest and as seen in Coker et al. In EGFR+PI3K combined drug inhibition we see the same discrepancy. Although we see correlation, this combination displays less additivity in HCC827 than our predictions would suggest. The consistent effects with both HCC827 and NCIH1792 suggest similarly that there are unknown mutations or amplifications within these cell lines causing additivity in ways that our model cannot predict[7].

The model correctly predicts additivity of MET inhibition with either JAK2 inhibition or EGFR/Erbb1 inhibition in NCIH1993. Predictions indicate this is because the MET receptor is amplified in NCIH1993, which drives oncogenic signalling primarily along down MAPK and AKT pathways, but JAK2 is still active. EGFR/Erbb1 is also able to activate JAK/STAT signalling and so combined inhibition of either of these proteins can produce an additive effect (Supplementary Figure S7G). Despite broad concordance between model and data, SW1573 appears to display some marginal additivity in the experimental data despite the model predicting it not to be additive (Supplementary Figure S7D). This cell line is characterised as having its oncogenic mutations in both PI3K and RAS, both downstream of MET (Figure 6G) and so the model does not predict an additive behaviour by targeting MET or JAK2. The experimental data from Nair et al. would suggest that MET plays a role in this cell line either in the response to inhibition of JAK2 or in that it is amplified in this cell line in a way that is uncharacterised.

### Supplementary Figures

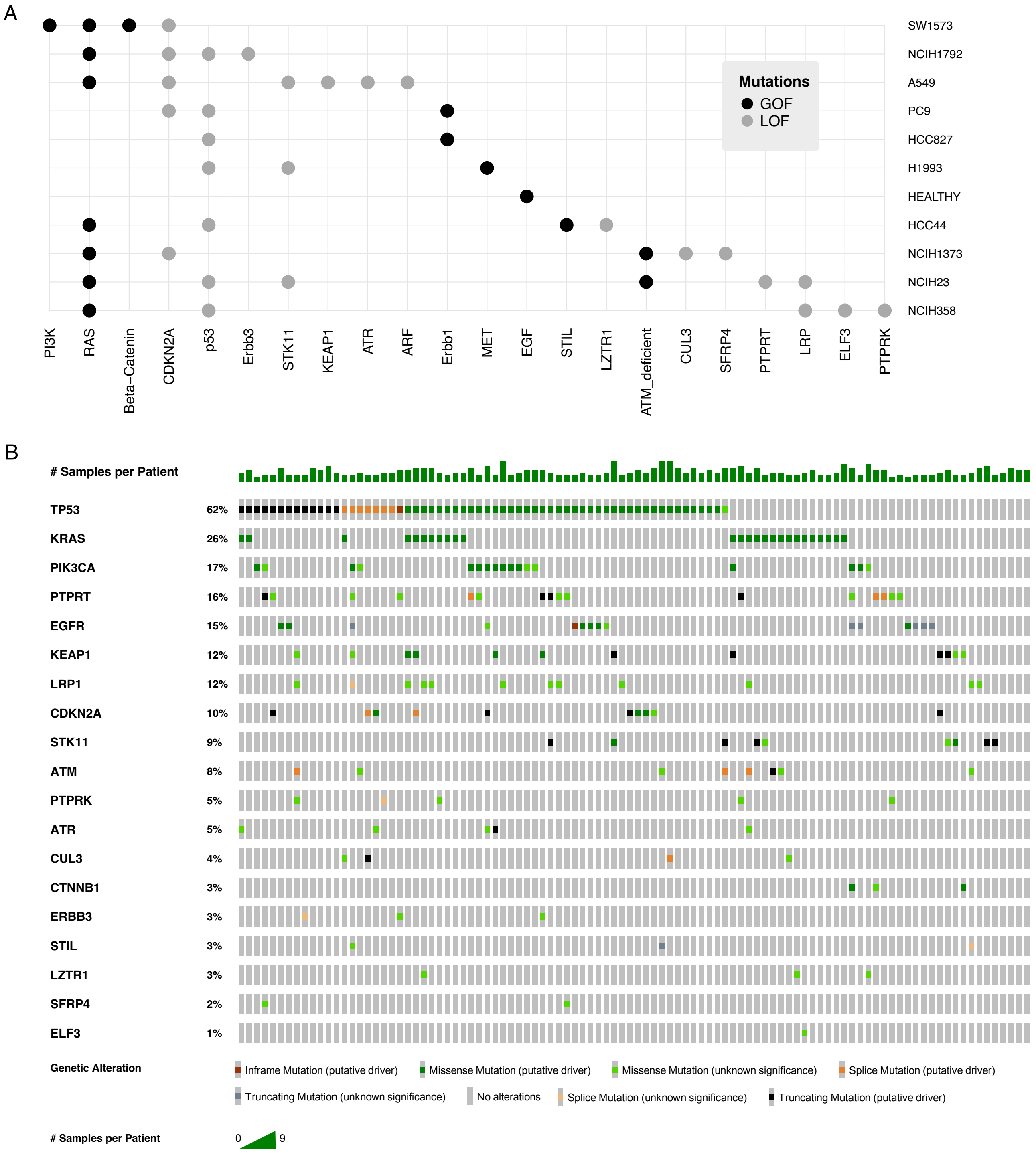


**Figure S1A-B. Summary of mutations in the modelled cell lines. A)** The schematic (right) shows mutations as columns and cell lines as rows, with gain-of-function mutations for a cell line and loss-of-function (grey). **B)** Schematic showing the presence of these mutations across clinical data (Plotted using cBio-Portal, using studies from TRACERx[96,97] ).

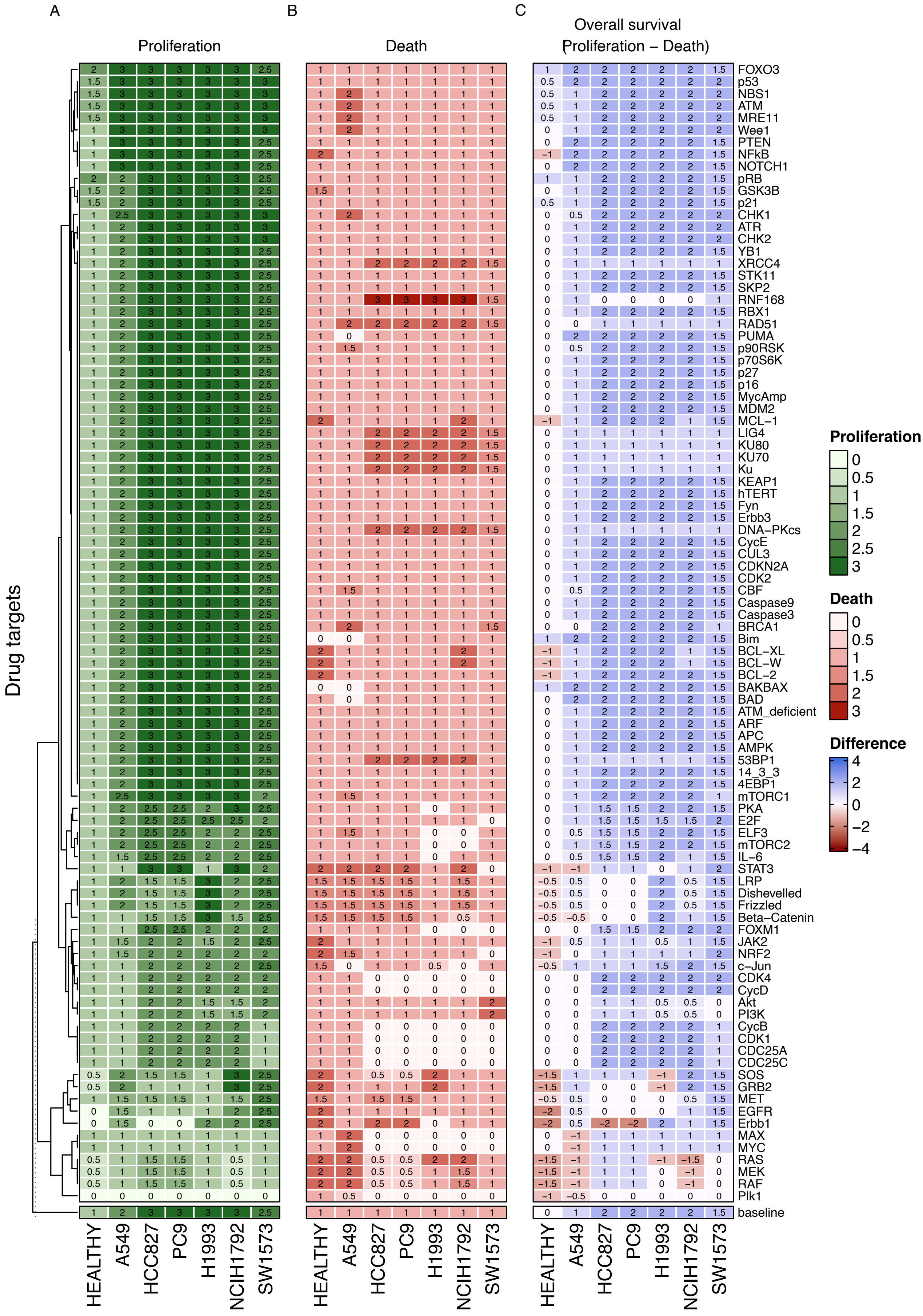


**Figure S2A-C. Heatmap showing the simulation of the model after perturbation with single drug treatments.** The columns are various cancer cell lines with healthy cells at the far left. Rows indicate druggable proteins that are included in the model. Values in the model correspond to proliferation (**A**, in green), death (**B**, in red) and the difference between proliferation and death (**C**, red to blue). For the difference between them, positive values (blue) indicates growth and negative values (red) indicate death. The baseline values for cell lines (readings without any perturbation) is shown on the bottom of the heatmaps.

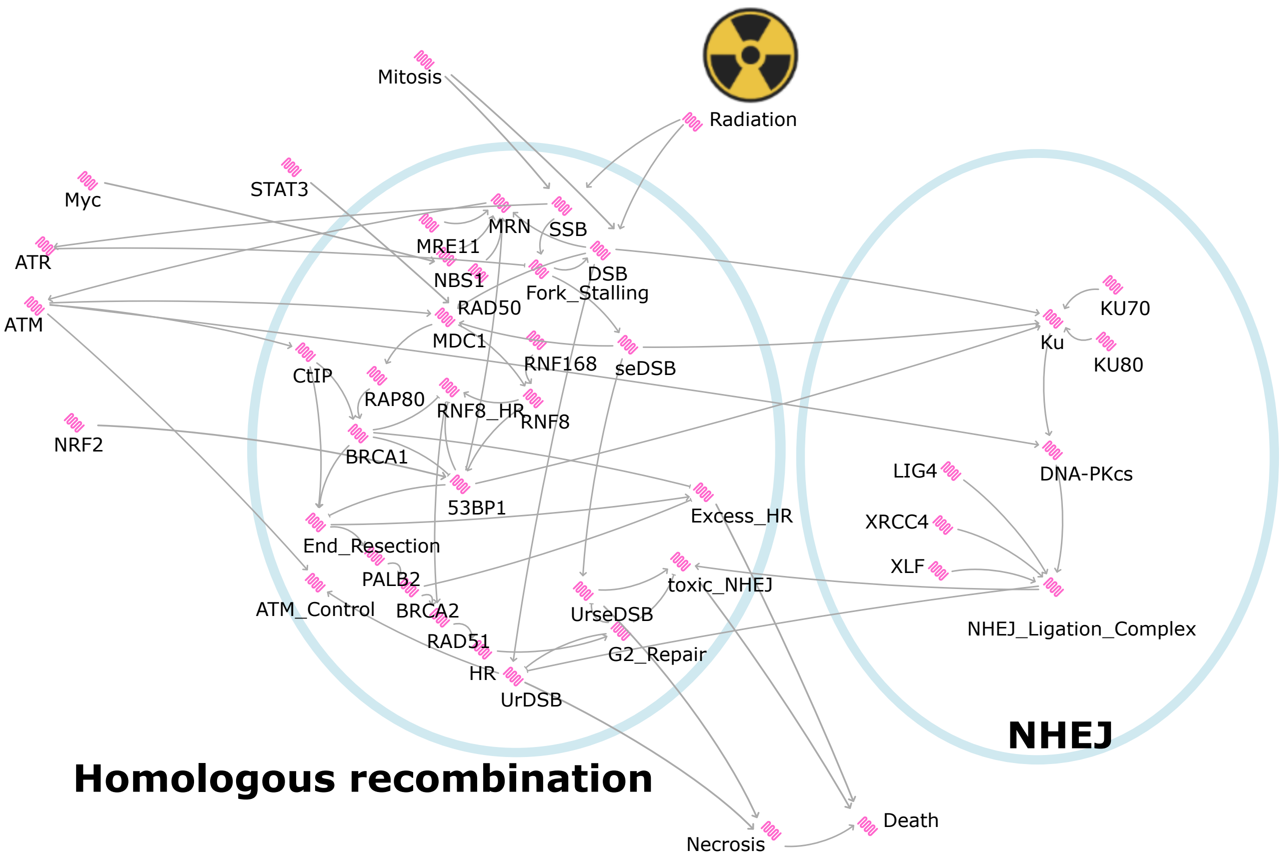


**Figure S3A. Supplementary for predictions pertaining to sensitivity to radiotherapy.** Schematic showing the part of the model that models irradiation and DNA-damage repair.

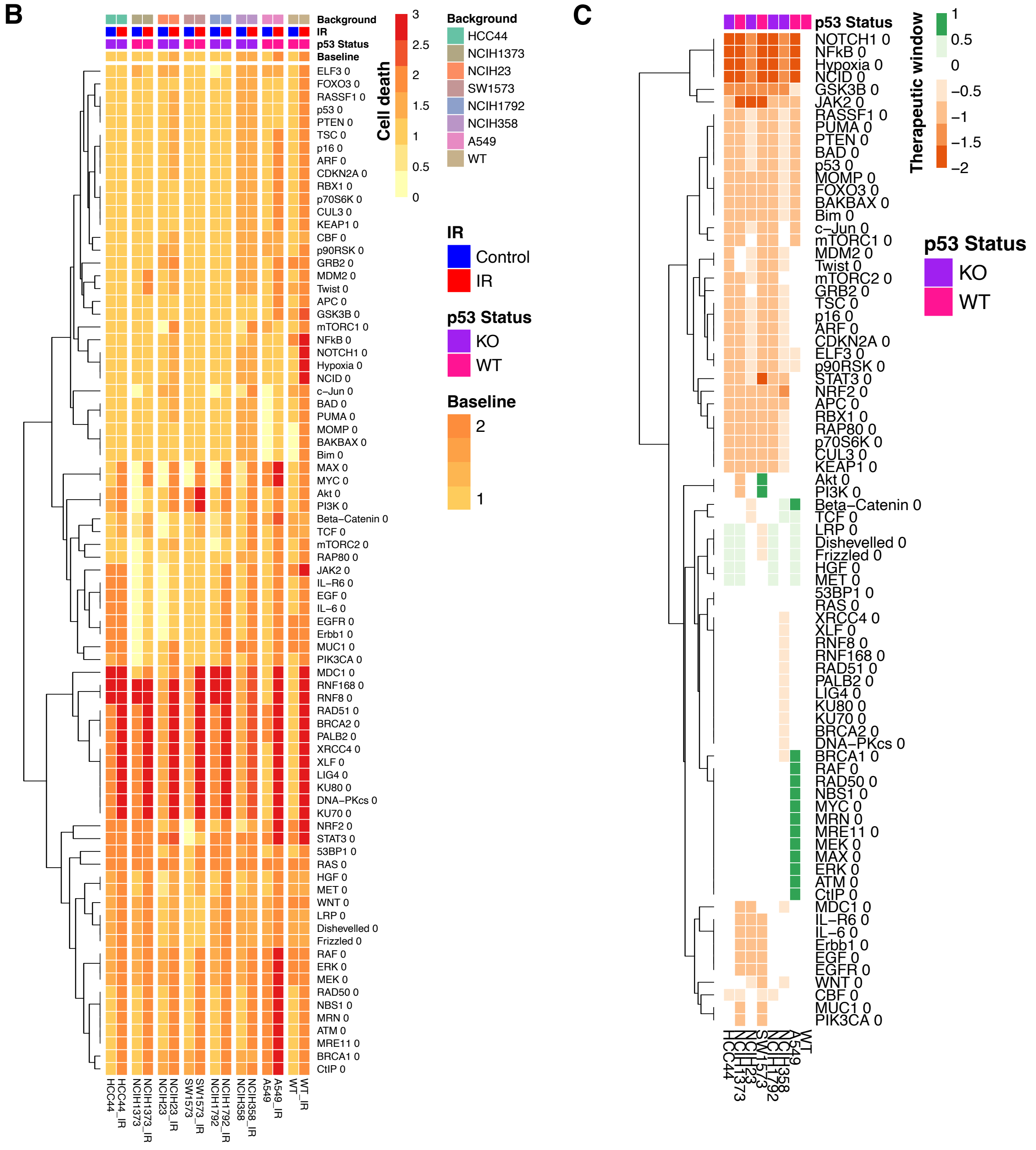

**Figure S3B-C. Predicted cell death and therapeutic window.** Effect of gene knock-outs (rows) on different cell line backgrounds (columns). Colour represents the activity of the node (cell death **(A)** and therapeutic window **(B)** difference between radiotherapy and knockout in cell line compared to WT cells).

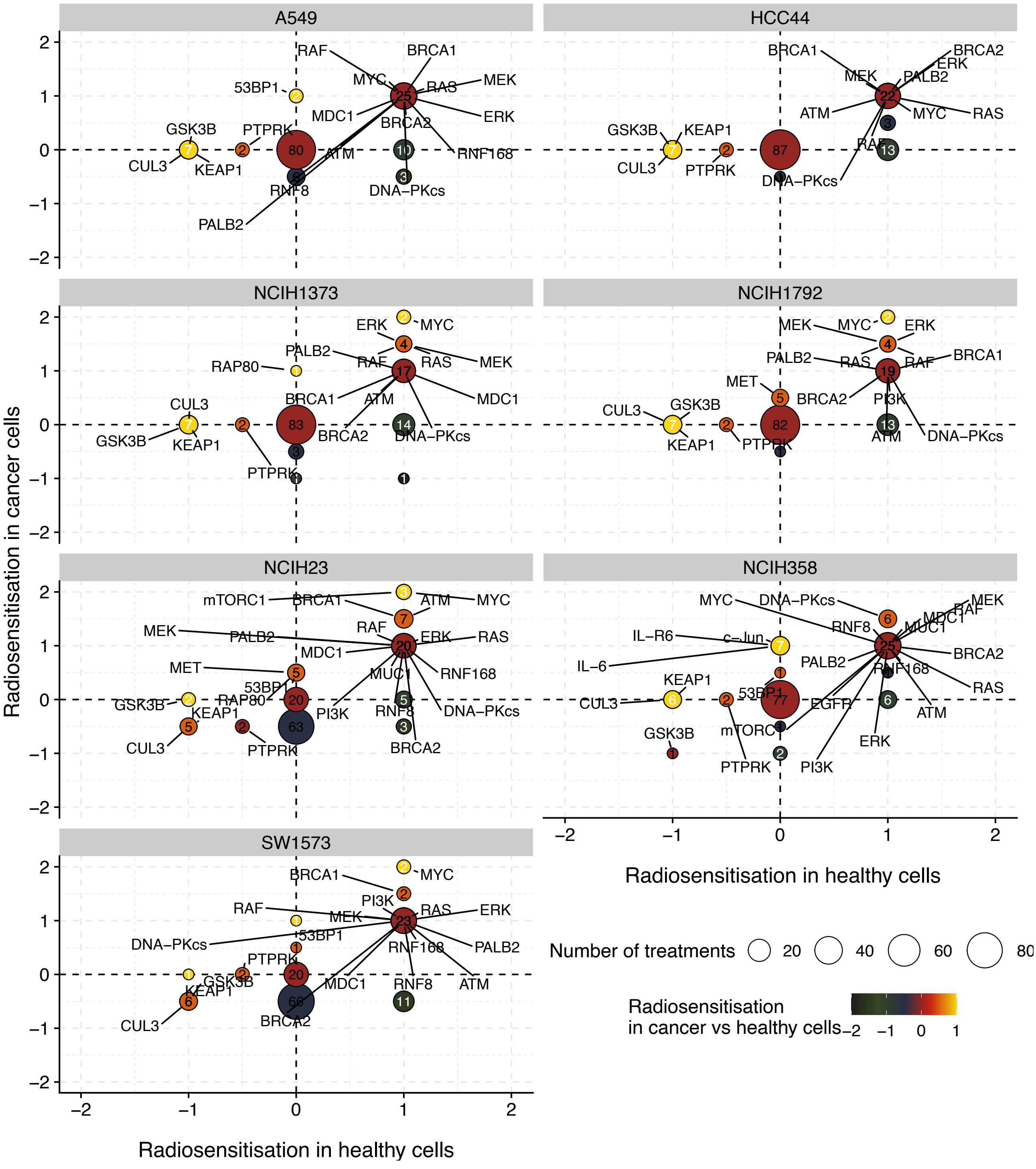


**Figure S3D. Scatter plots showing the difference in DNA-damage associated radiosensitivity in cancer (y axis) versus healthy cells (x axis).** Black is more radio-sensitisation of cancer than healthy, yellow is more radio-sensitisation of healthy than cancer. Point size is the number of knockouts that have this effect.

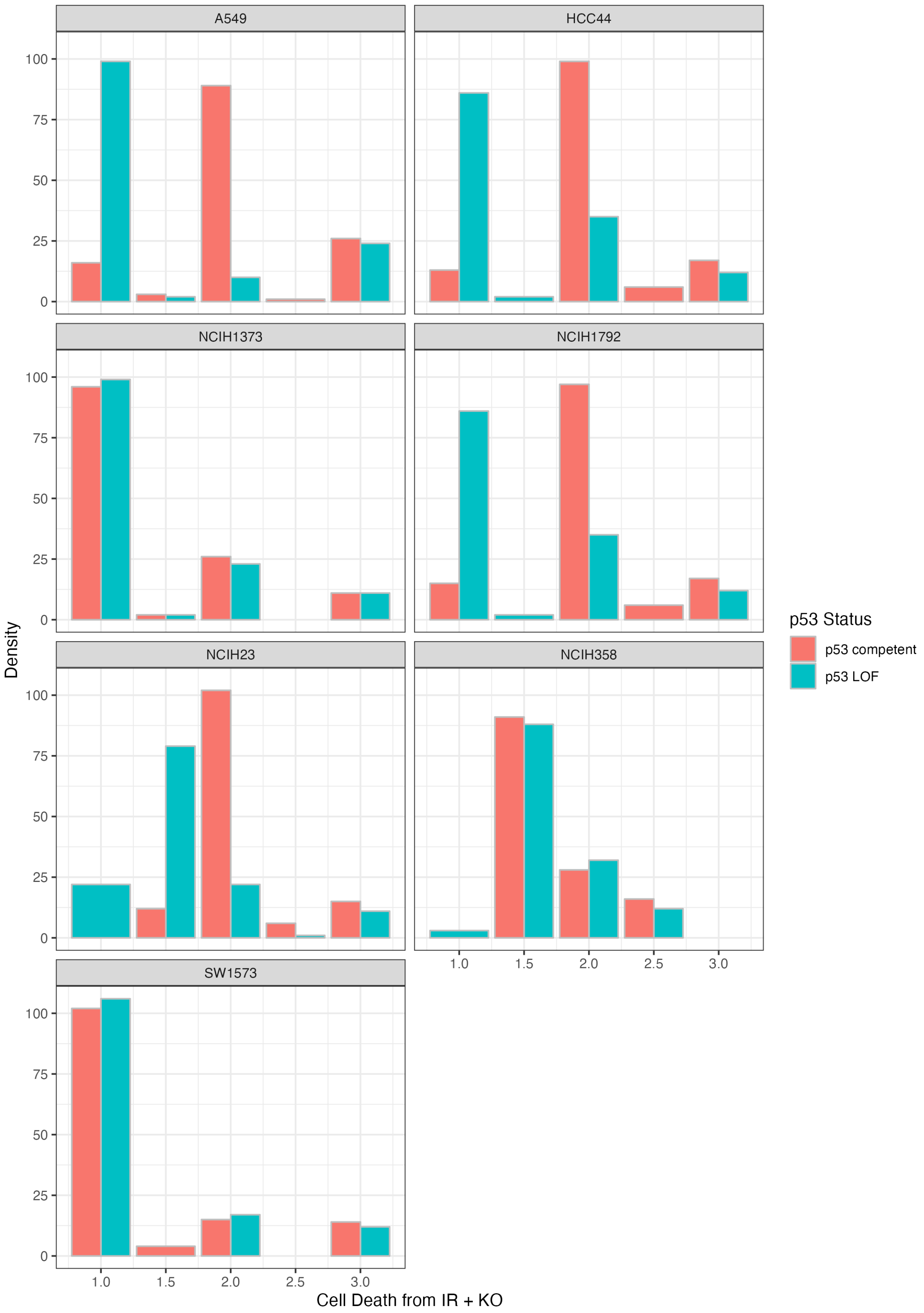


**Figure S4A. The effect of switching p53 status on response to radiotherapy continued.** Level of cell death (x-axis) in each cell line for each simulated knock-out in combination with IR (y-axis) comparing the effect in a p53-competent (blue) or a p53-deficient (red) version of that cell line.

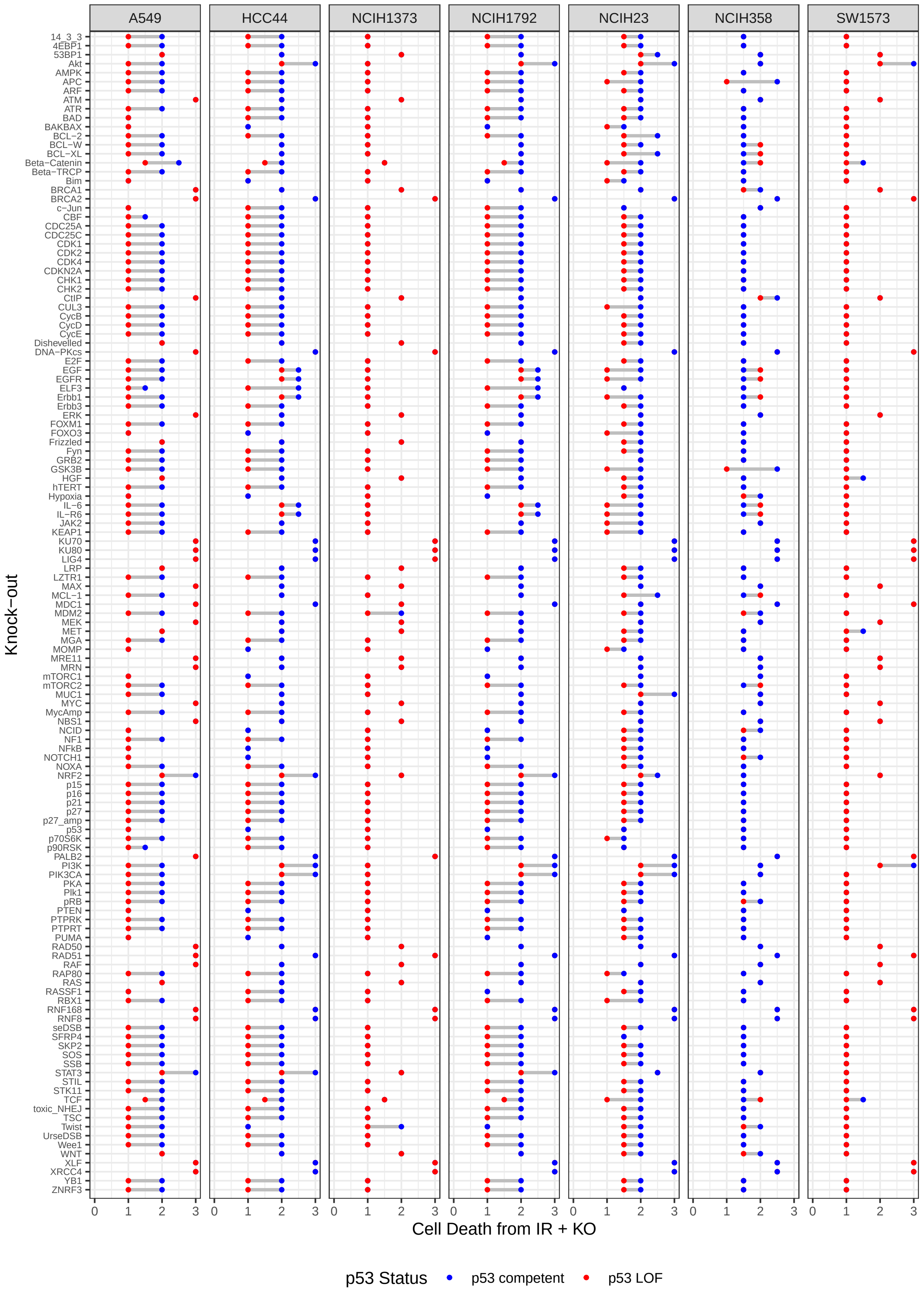


**Figure S4B. The effect of switching p53 status on response to radiotherapy continued.** Number of simulated knockouts leading to each possible level of cell death for p53-competent (blue) and p53-deficient (red) cell lines.

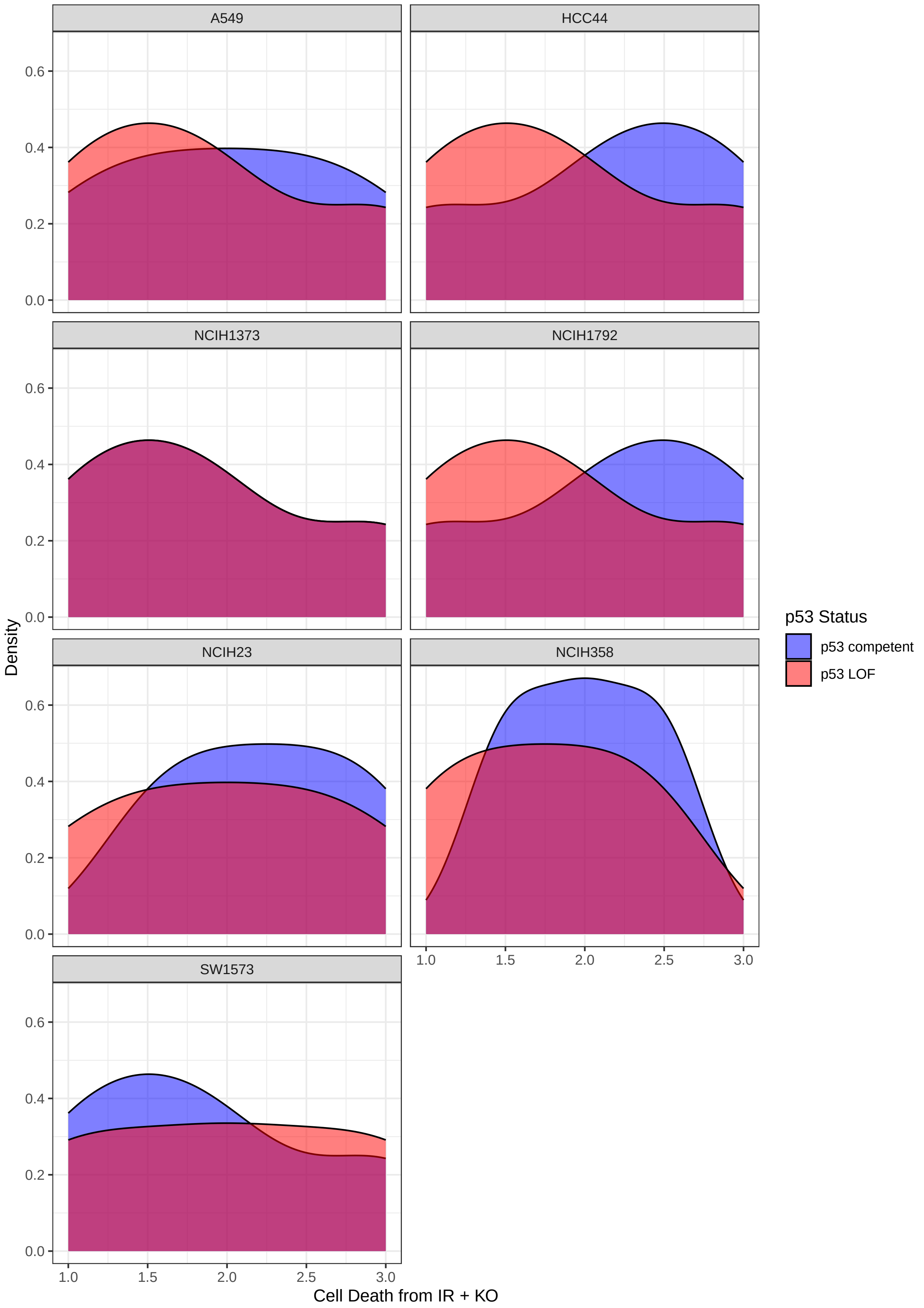


**Figure S4C. The effect of switching p53 status on response to radiotherapy continued.** Density plot of number of simulated knock-outs leading to each possible level of cell death for p53-competent (blue) and p53-deficient (red) cell lines.

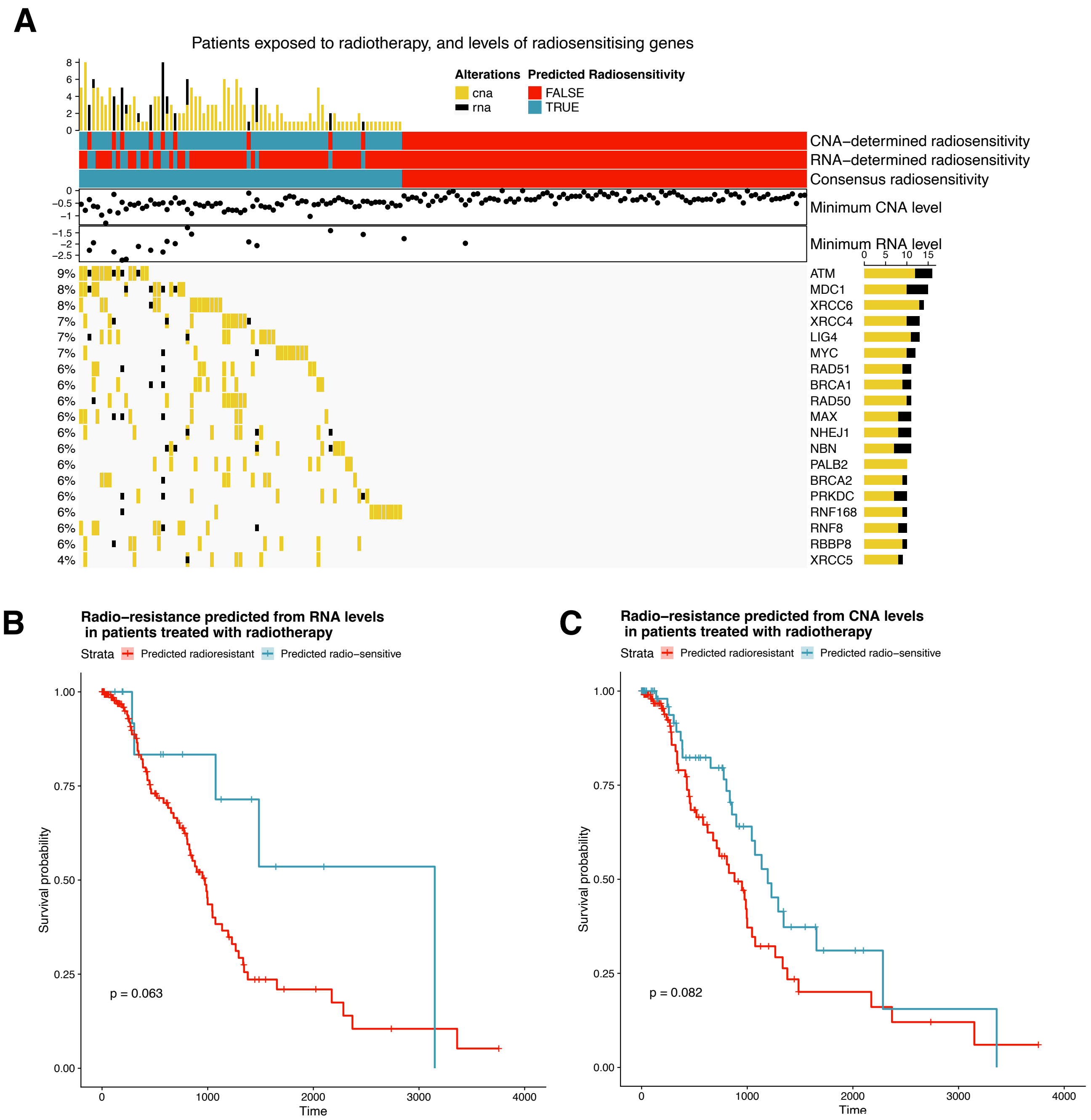


**Figure S5A-C. In silico predictions stratify radiation-treated patients into radiosensitive and radioresistant strata. C)** Plot illustrating TCGA data of radiation treated patients (columns) according to the basal activity of predicted radio-sensitising genes (rows). Cells indicate the presence of low levels of CNV (yellow) or expression (black) for a particular gene. The predicted radiosensitivity of the patients is marked blue (sensitive) or red (resistant) by the bars on the top. Also shown is the minimum RNA expression and CNV levels for each patient and histograms showing the frequency of alterations for each patient (top) and gene (left). **B, C)** In silico predictions stratify radiation-treated patients into radiosensitive and radioresistant strata. Kaplan-Meier survival curves showing the differences in survival (days) between radioresistant patients (red) and radiosensitive patients (blue) for patients treated with radiation. These patients were stratified based on their expression levels **(B)** or their copy-number levels **(C)**.

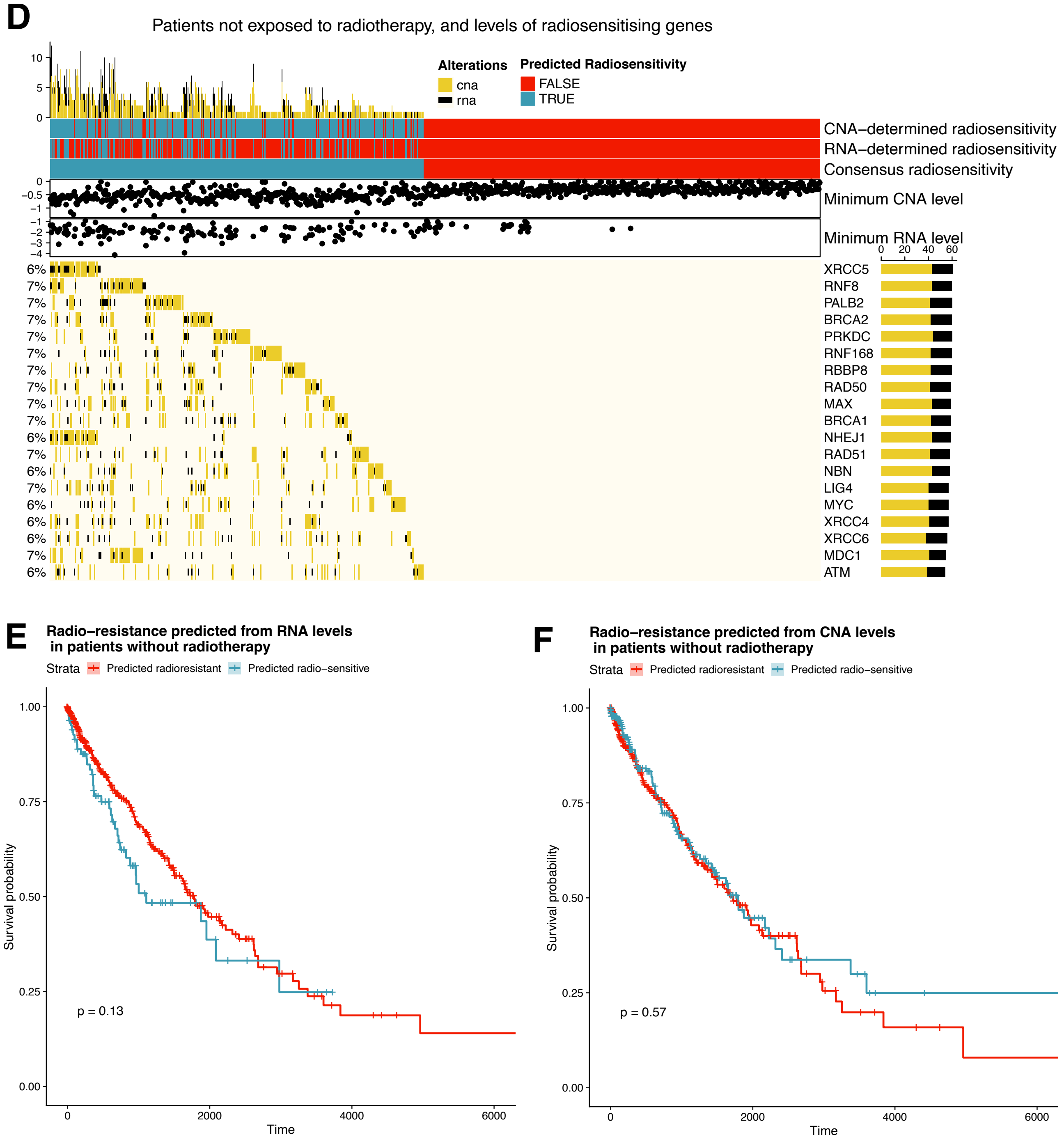


**Figure S5D-F. In silico predictions stratify radiation-treated patients into radiosensitive and radioresistant strata. D)** Plot illustrating TCGA data of patients not treated with radiotherapy (columns) according to the basal activity of predicted radiosensitising genes (rows). Cells indicate the presence of low levels of CNV (yellow) or expression (black) for a particular gene. The predicted radiosensitivity of the patients is marked blue (sensitive) or red (resistant) by the bars on the top. Also shown is the minimum RNA expression and CNV levels for each patient and histograms showing the frequency of alterations for each patient (top) and gene (left). **E, F)** Kaplan-Meier survival curves showing the differences in survival (days) between predicted radioresistant patients (red) and radiosensitive patients (blue) for patients treated without radiotherapy. These patients were predicted radioresistant according to expression levels **(E)** or copy number levels **(F)**. The x-axis indicates the time since diagnosis (days) and y-axis indicates the survival probability.

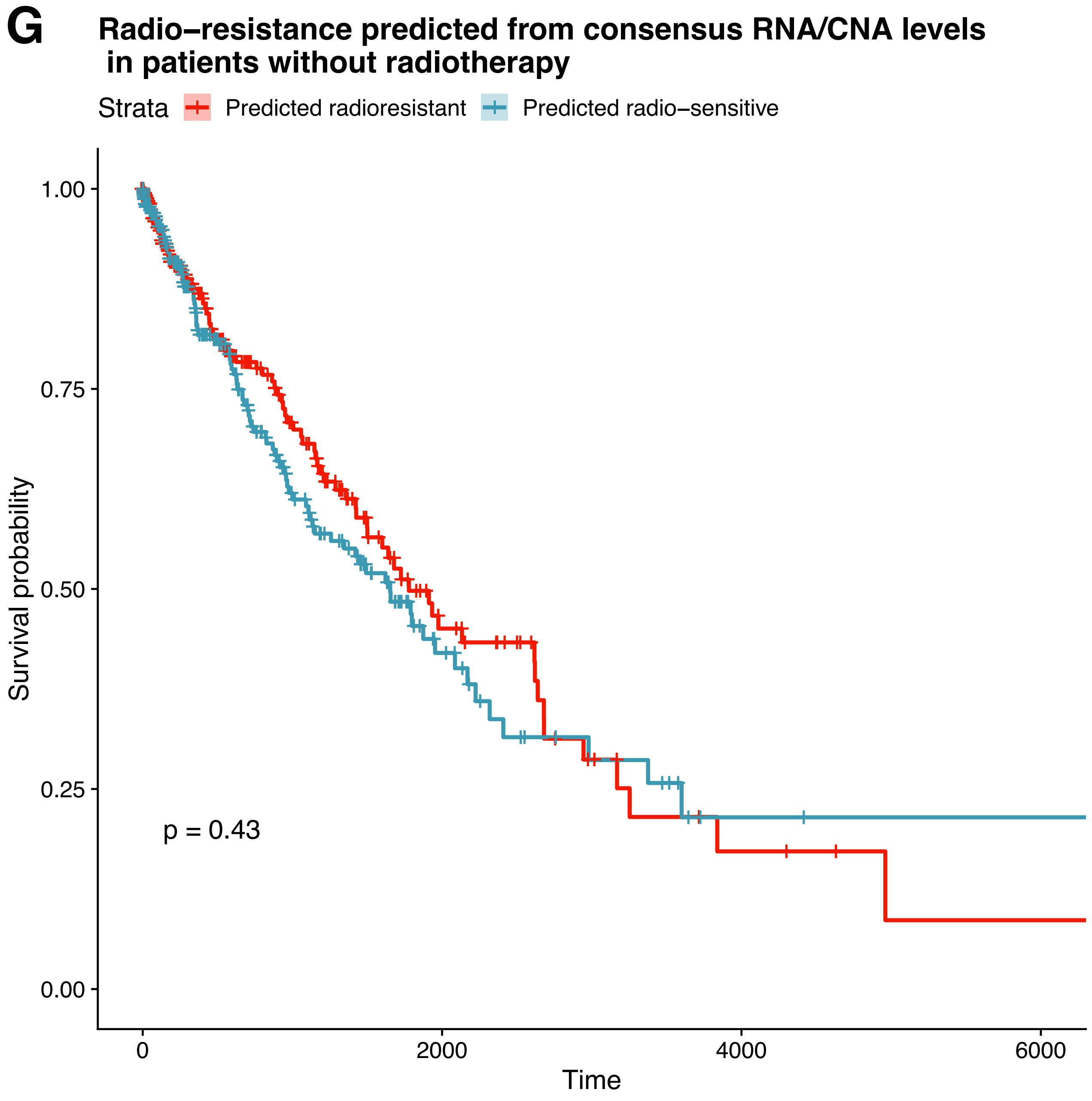


**Figure S5G. In silico predictions stratify radiation-treated patients into radiosensitive and radioresistant strata.** Kaplan-Meier survival curves showing the differences in survival (days) between predicted radioresistant (according to consensus expression and copy number levels) patients (red) and radiosensitive patients (blue) for patients treated without radiation.

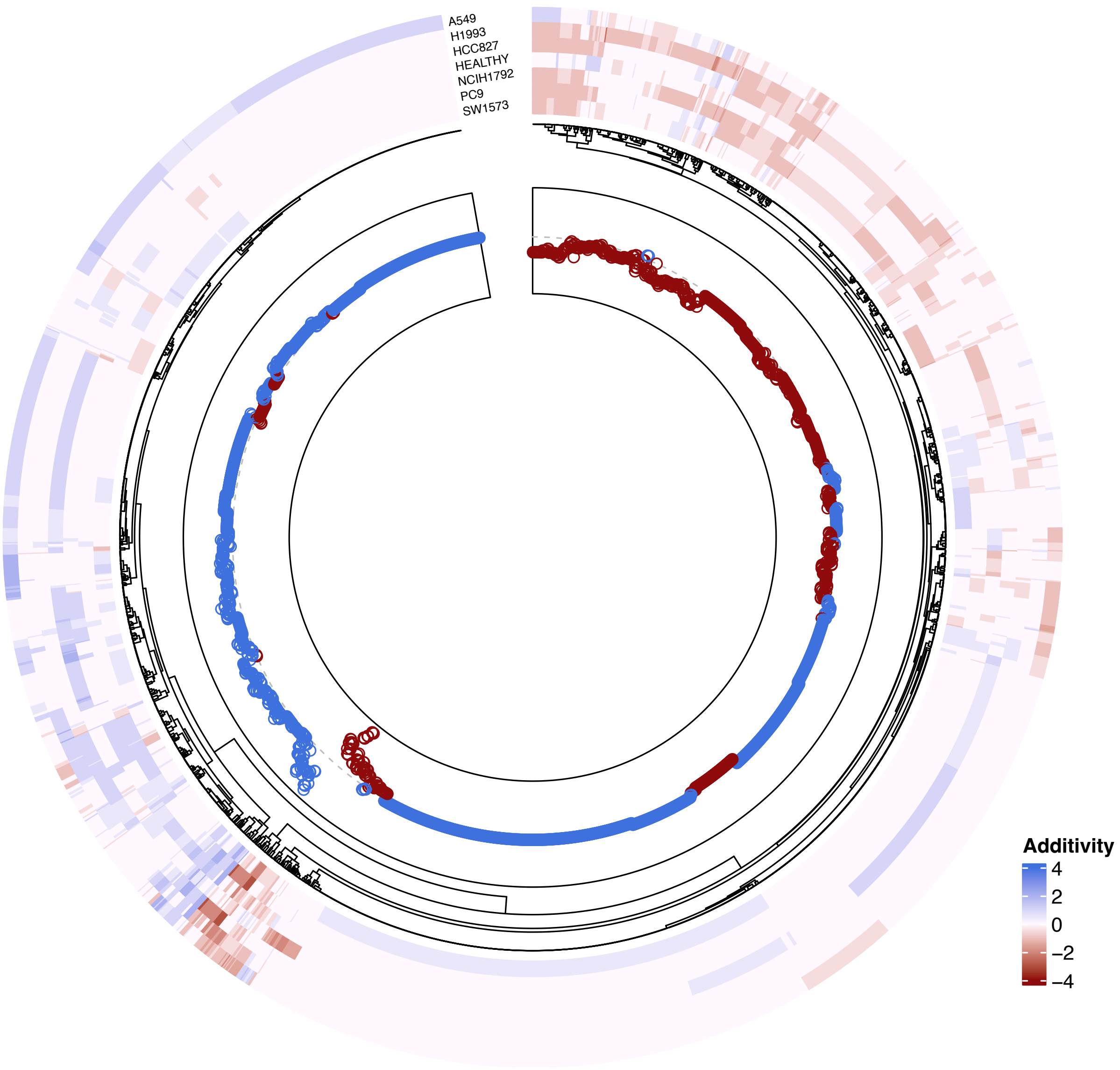


**Figure S6A. Heatmap showing additivity of drug combinations on selected NSCLC cell lines.** Columns (spokes of the wheel) are the drug combinations and rows are cell line. A darker red colour indicates those combinations that have a greater additive effect, while blue represents those drugs with antagonistic effects (the drugs cancel one another out). The plot in the inner wheel of the circle shows the mean additive effect of a drug across all cell lines, and is red for those drug combinations with overall additive effect and blue for those antagonise each other.

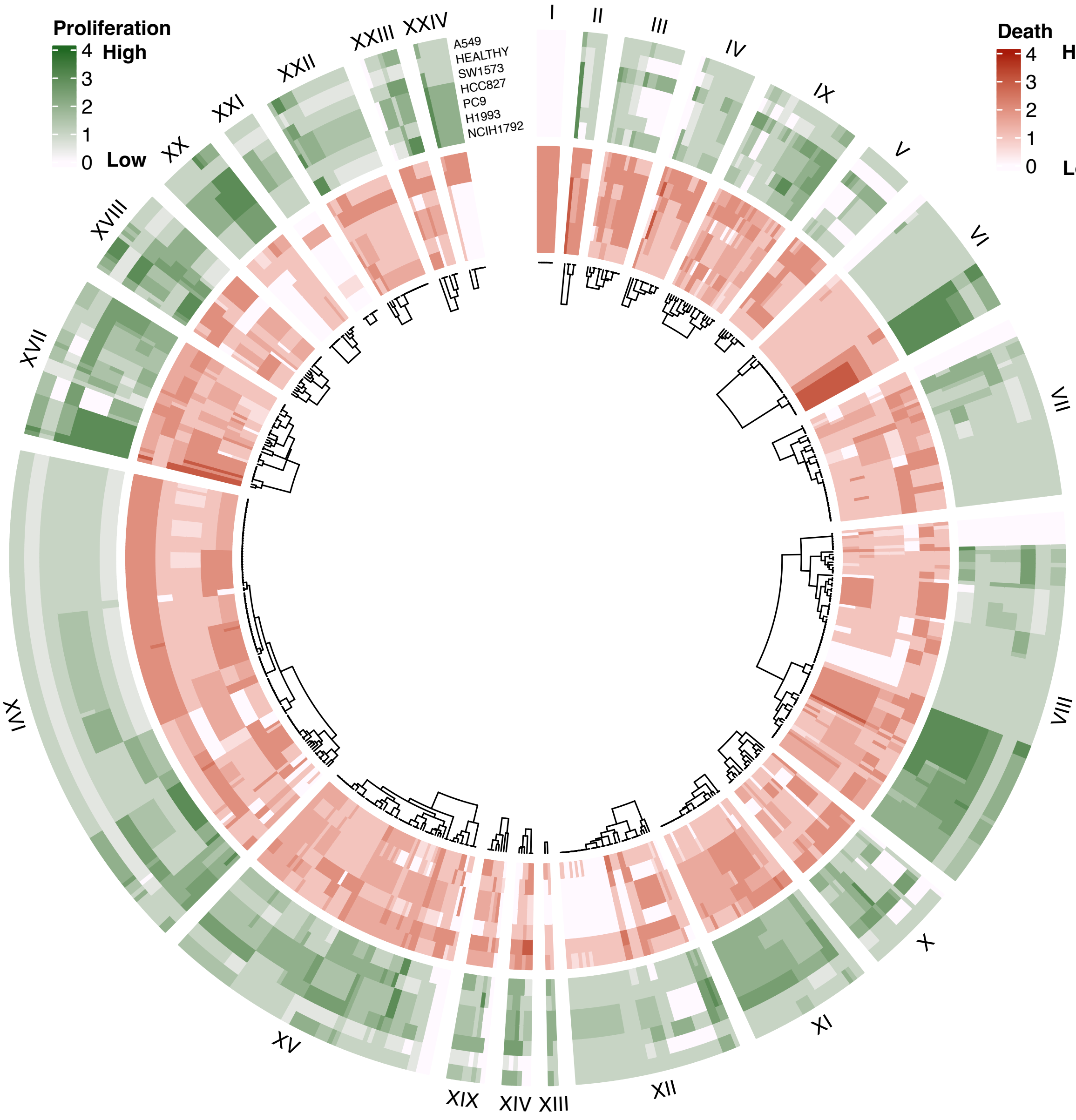


**Figure S6B. Heatmap showing effect of drug combinations on selected NSCLC cell lines.** Heatmap showing death (in red, inner circle) and proliferation (in green, outcircle). Columns (spokes of the wheel) are the drug combinations and rows are cell lines. Darker colours indicate higher values for the phenotypes.

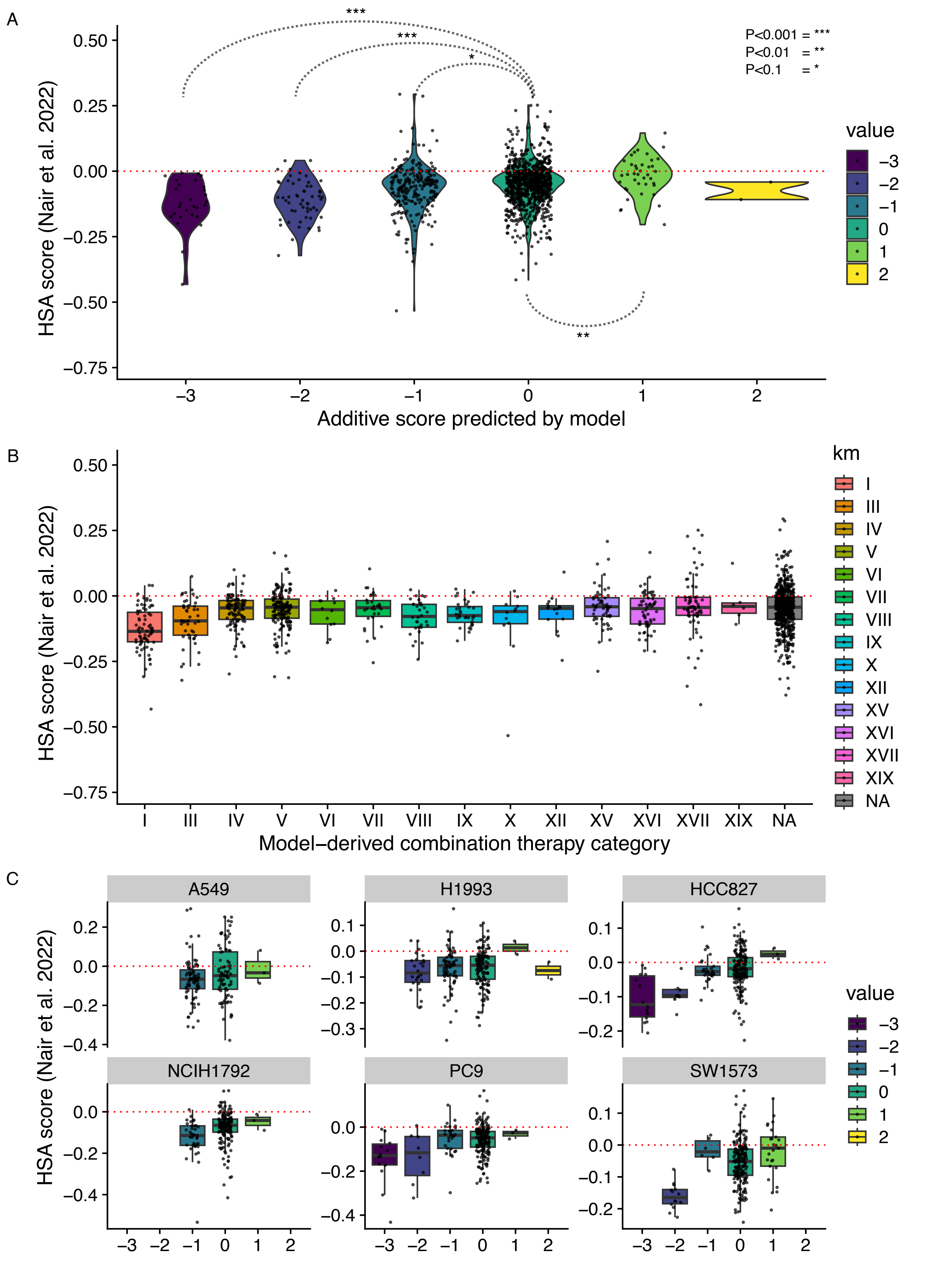


**Figure S7A-C. Concordance of simulated results for drug combinations with Nair *et al.* (2022). A)** Violin plot showing the additive score as predicted by the model (x axis, see Methods) versus the HSA measured experimentally by Nair *et al.* (2022) (y axis). Significance is illustrated through dotted lines from those drug combinations that are not predicted to be additive. **(B)** Visualisation of predicted additivity (x axis) of drug combinations versus experimentally measured HSA (y axis) for each drug cluster. **C)** Same visualisation as panel A, but broken down by cell lines tested by Nair *et al.* (2022).

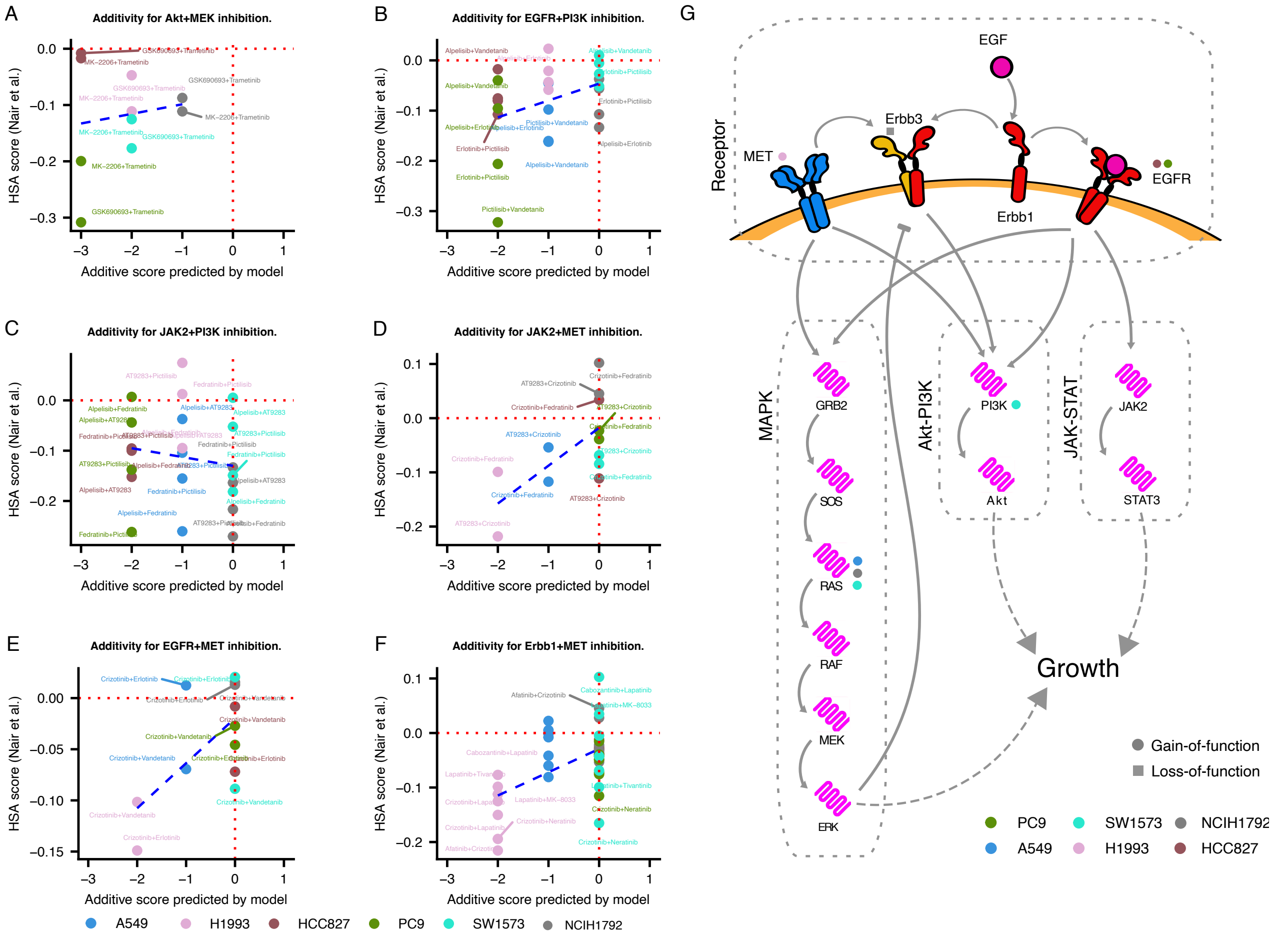


**Figure S8A-G. Concordance of simulated results with published results for individual drugs. (A-F)** Experimentally measured additivity (on the y-axis, HSA from Nair *et al.* (2022)) compared with the computational predicted value (x-axis) for Akt + MEK inhibition **(A)**, EGFR + PI3K inhibition **(B)**, JAK2 + PI3K inhibition **(C)**, JAK2 + MET inhibition **(D)**, EGFR + MET inhibition **(E)** and Erbb1 + MET inhibition **(F)**. The names of the drugs used are labelled on the points and the colours refers to the cell line with which the reading was taken. **G)** Subnetwork of the model illustrating the mutations in the cell lines affecting growth pathways and the regulatory interactions between them. Edges represent regulatory interactions with a pointed arrowhead referring to an activatory interaction and a blunt arrowhead indicating inhibitory interactions. Coloured circles indicate where a cell line has a gain-of-function mutation and square circles indicate where they have a low-of-function mutation.

H

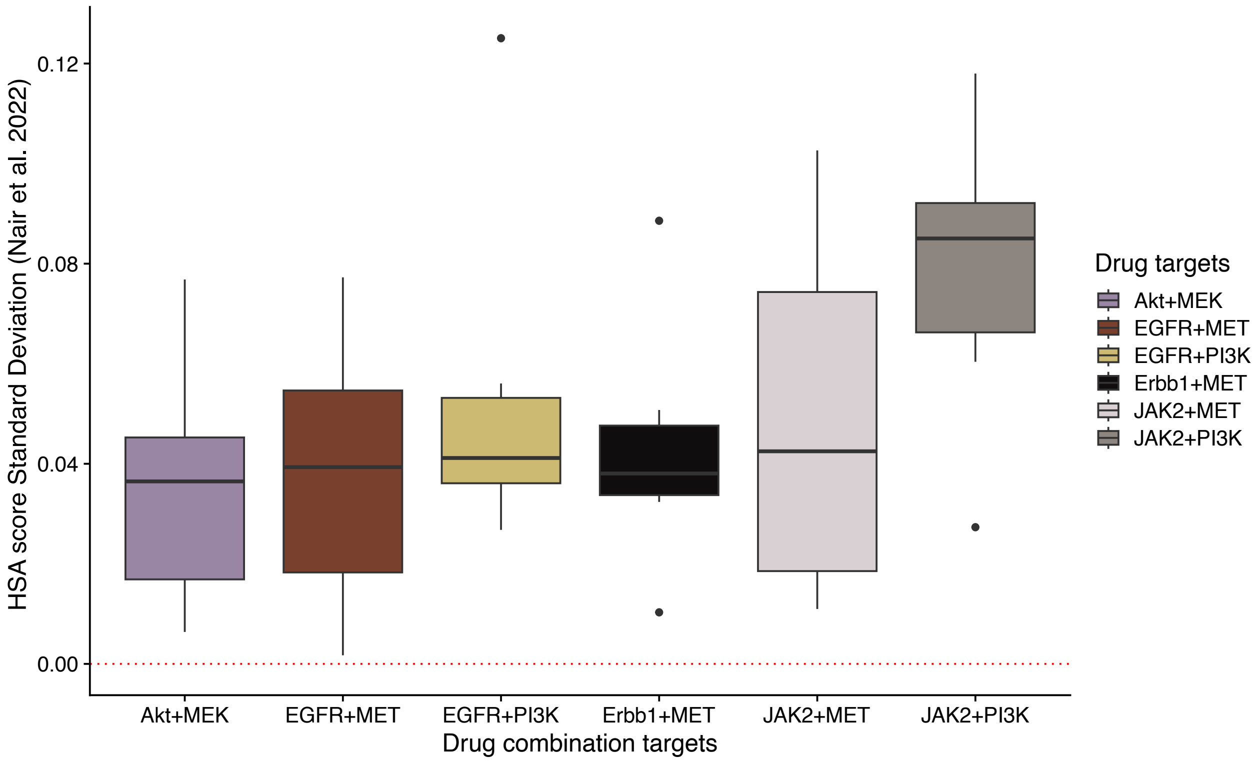


I

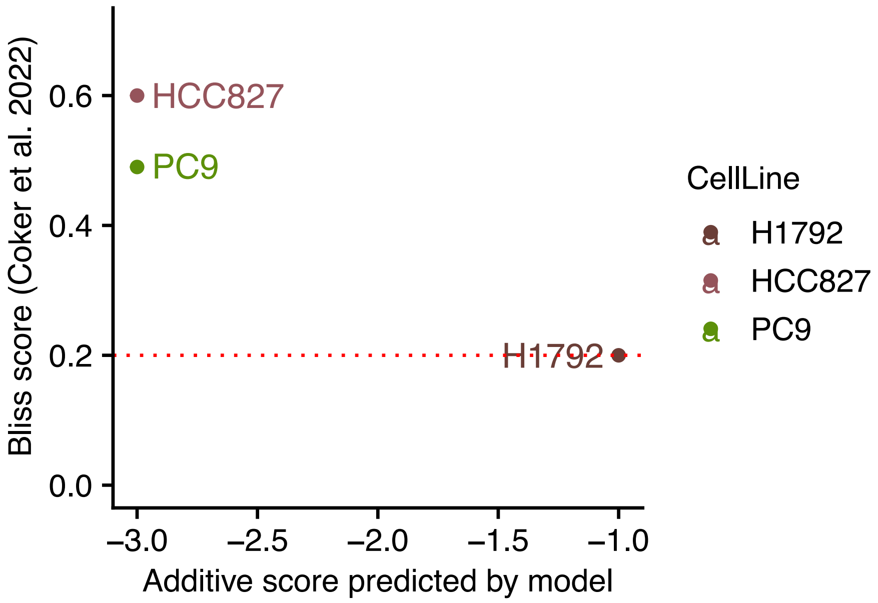


**Figure S8H-I. Concordance of simulated results with published results for individual drugs continued. (H)** Boxplot illustrating the standard deviation of additivity (y axis) of different drugs targeting specific proteins (x axis), taken from Nair *et al.* (2022). **(I)** Plot comparing the predicted additivity of the model for MAPK+Akt inhibition (x axis) with BLISS score (y axis), taken from Coker et al. The lower the BLISS score, the more synergistic the drug treatment is.

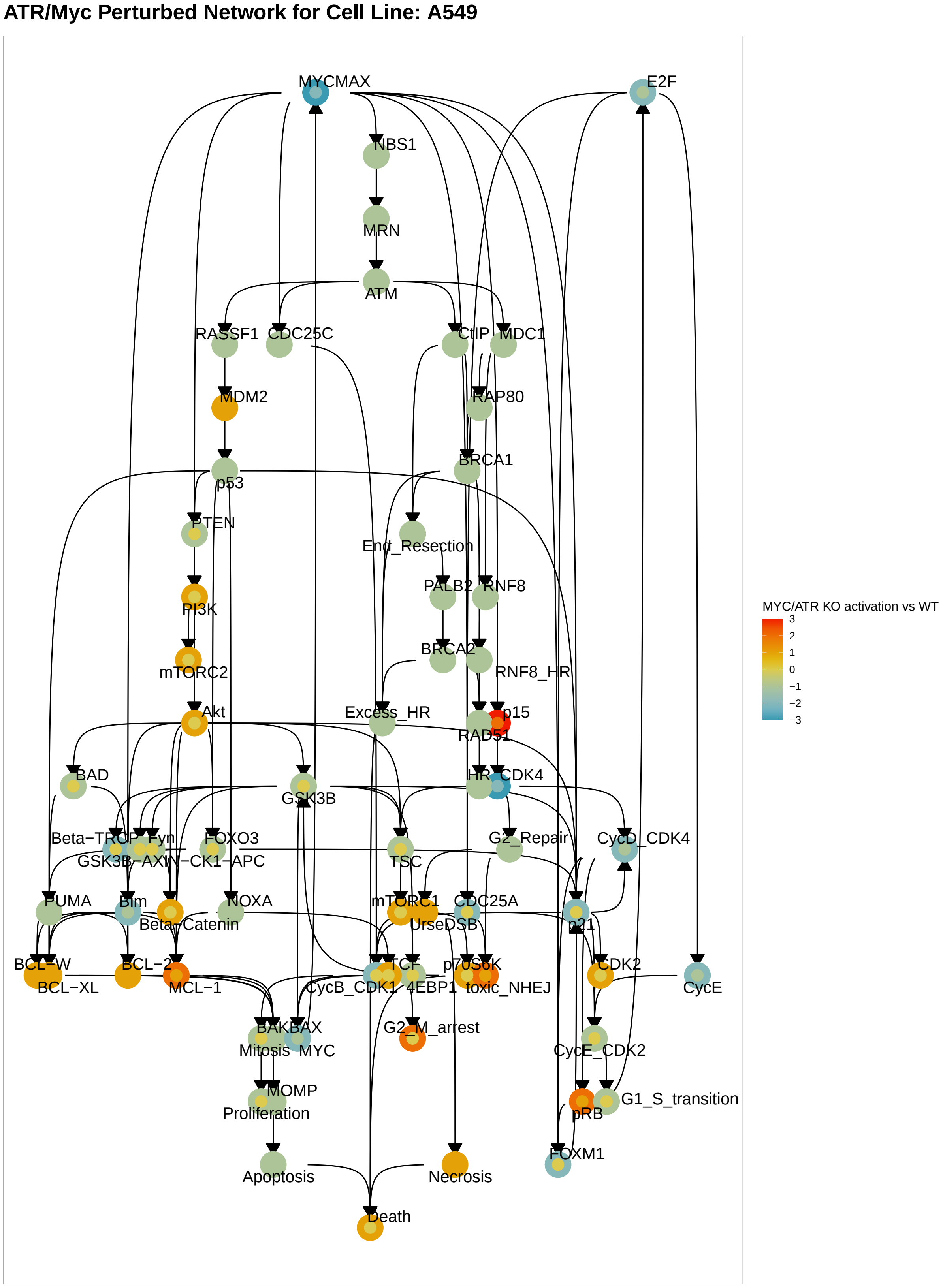


**Figure S9A. Mechanistic analysis of combined ATR and Myc inhibition.** Subnetwork of the model, covering the proteins and variables that changed after treatment of Myc/ATR inhibitors in A549. Colour of the node illustrates the difference between ATR/Myc inhibition vs untreated (blue for a relative decrease in activation and red for an increase) in PC9 (outer circle) and the difference in healthy lung cells (inner circle).

**ATR/Myc Perturbed Network for Cell Line: H1993**

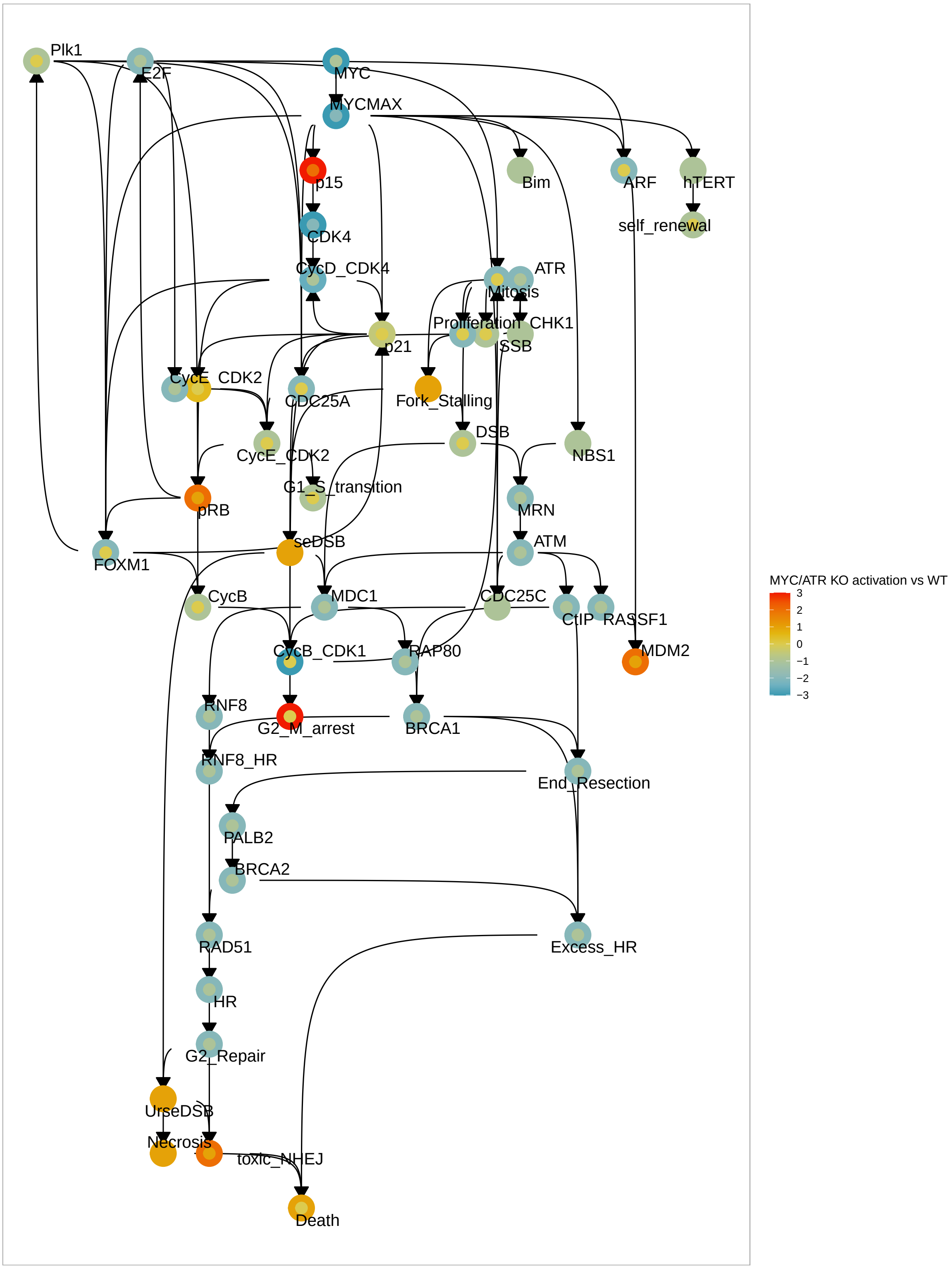


**Figure S9B. Mechanistic analysis of combined ATR and Myc inhibition.** Subnetwork of the model, covering the proteins and variables that changed after treatment of Myc/ATR inhibitors in H1993.

**ATR/Myc Perturbed Network for Cell Line: HCC827**

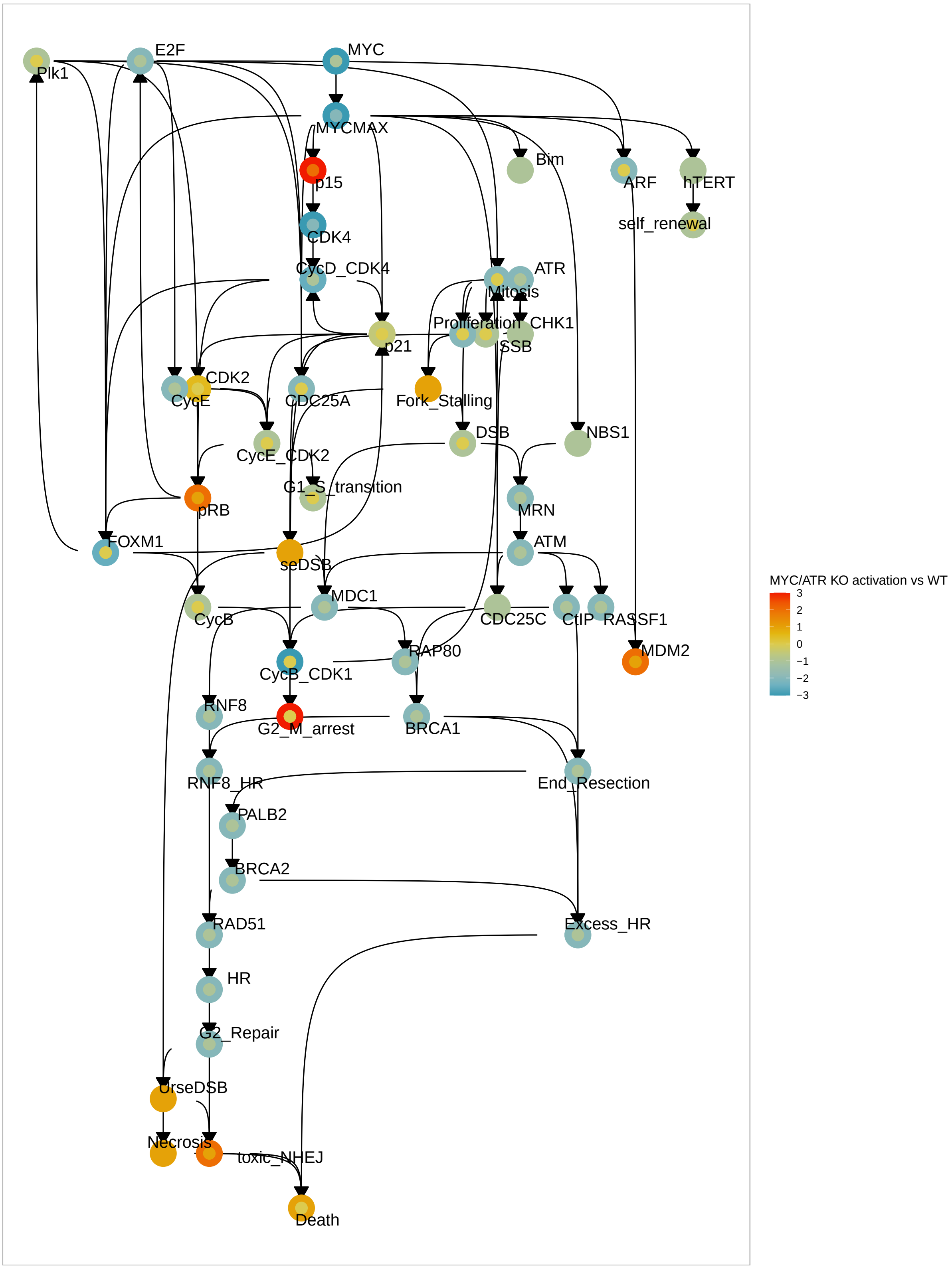


**Figure S9C. Mechanistic analysis of combined ATR and Myc inhibition.** Subnetwork of the model, covering the proteins and variables that changed after treatment of Myc/ATR inhibitors in HCC827.

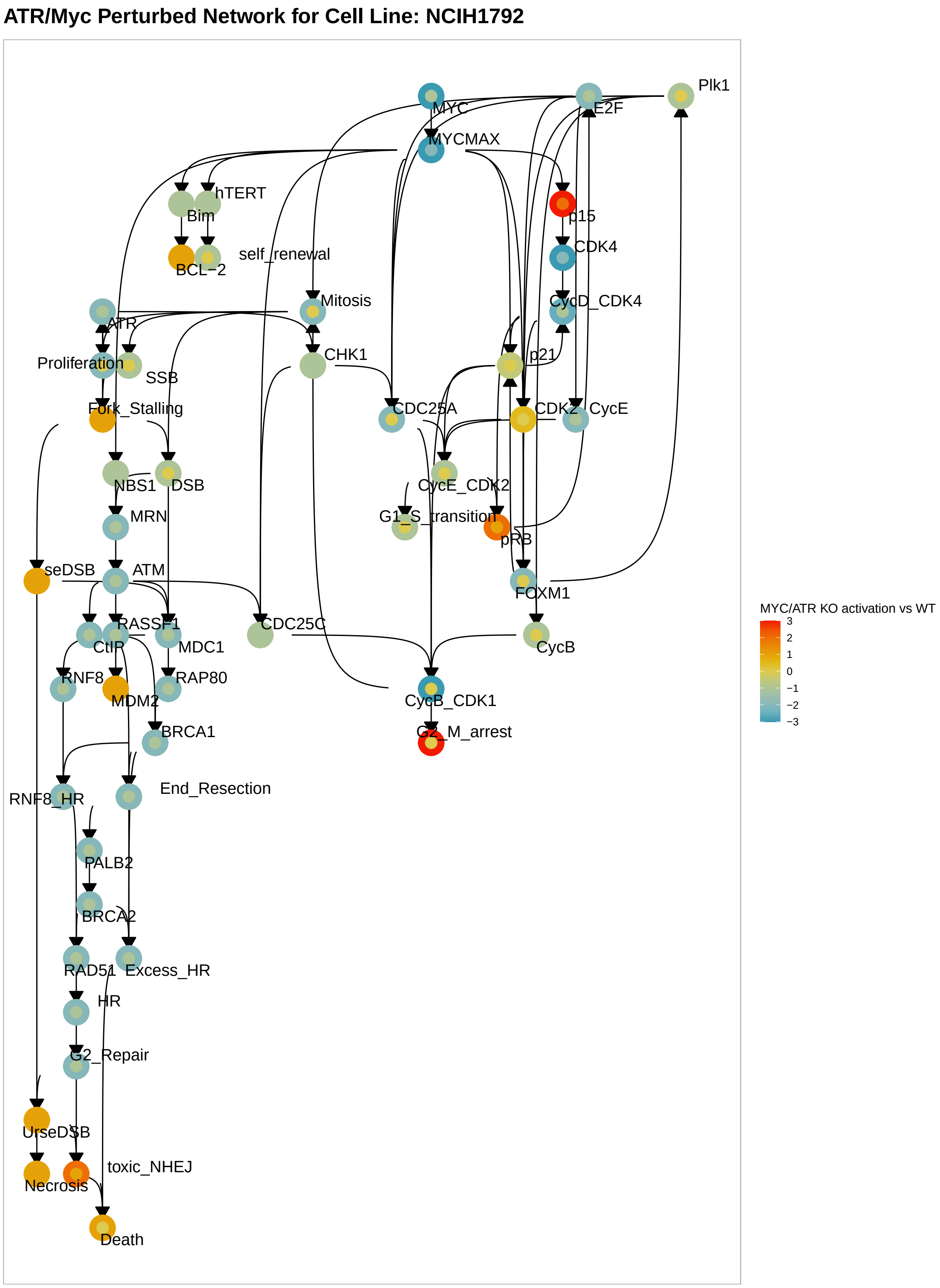


**Figure S9D. Mechanistic analysis of combined ATR and Myc inhibition.** Subnetwork of the model, covering the proteins and variables that changed after treatment of Myc/ATR inhibitors in NCIH1792.

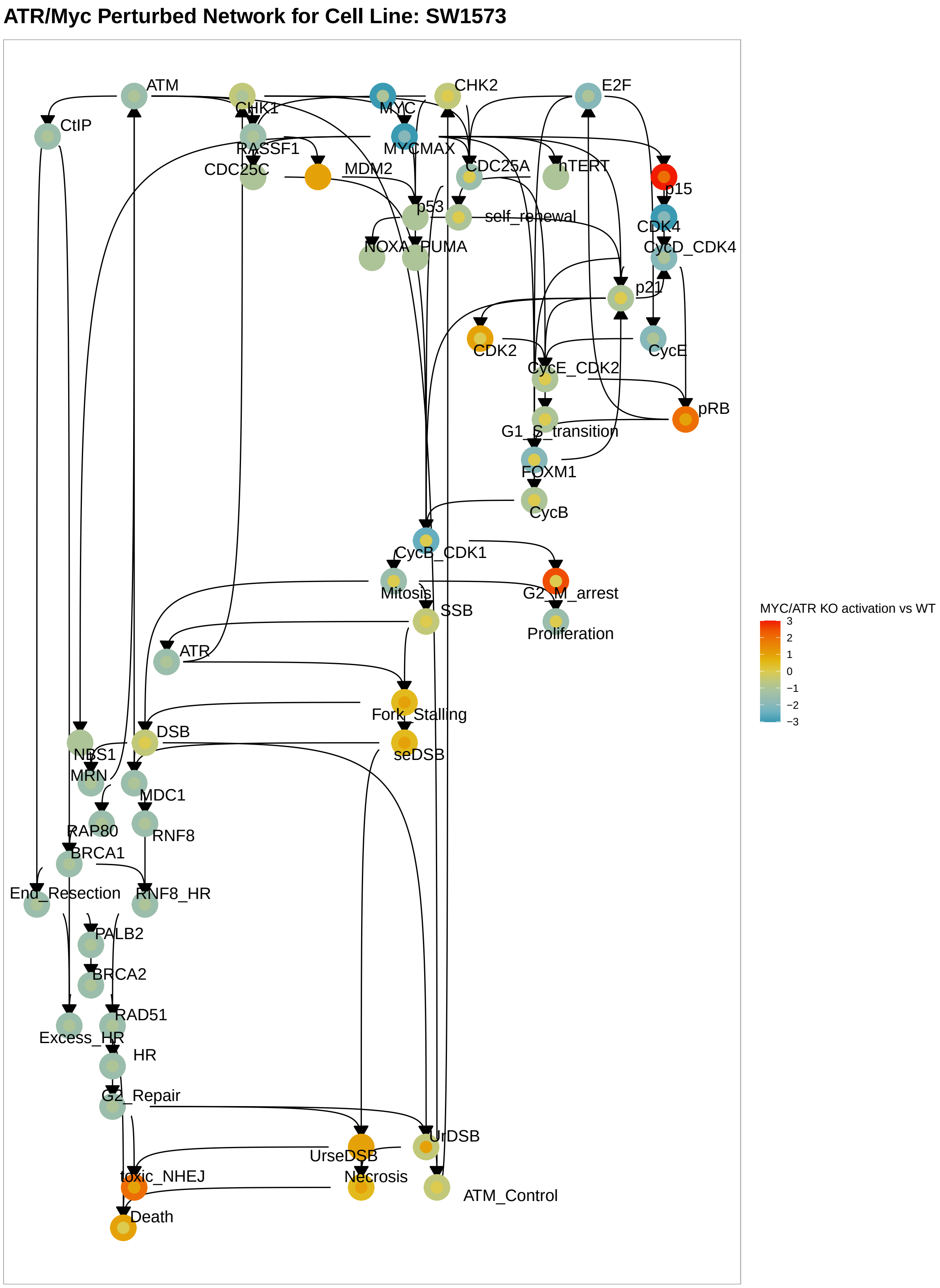


**Figure S9E. Mechanistic analysis of combined ATR and Myc inhibition.** Subnetwork of the model, covering the proteins and variables that changed after treatment of Myc/ATR inhibitors in SW1573.

### Supplementary Tables

**Table S1. Regulatory edges in the *in silico* model.**

| From | To | Type | Citation |
|---|---|---|---|
| 14_3_3 | CDC25C | Inhibitor | Graves et al. 2001 |
| 53BP1 | End_Resection | Inhibitor | Zimmermann et al. 2013 |
| 53BP1 | Ku | Activator | Adams and Carpenter. 2006 |
| 53BP1 | MRN | Activator | Lee et al. 2010 |
| 53BP1 | RNF8_HR | Inhibitor | Nakada et al. 2012 |
| Akt | AMPK | Inhibitor | Hahn-Windgassen et al. 2005 |
| Akt | Bim | Inhibitor | Qi et al. 2006 |
| Akt | c-Jun | Activator | Go et al. 2001 |
| Akt | Caspase9 | Inhibitor | Cardone et al. 1998 |
| Akt | hTERT | Activator | Daniel et al. 2012 |
| Akt | MCL-1 | Activator | Fennel and Swanton. 2012 |
| Akt | NFkB | Activator | Kucuksayan et al. 2016 |
| Akt | NRF2 | Activator | Wang et al. 2008 |
| Akt | TSC | Inhibitor | Vicary and Roman. 2016 |
| APC | GSK3B-AXIN-CK1-APC | Activator | Shang et al. 2017 |
| Apoptosis | Death | Activator | Roos and Kaina. 2006 |
| ATM | ATM_Control | Activator | Buscemi et al. 2004 |
| ATM | CDC25C | Inhibitor | Thanasoula et al. 2012 |
| ATM | CtIP | Activator | Han et al. 2021 |
| ATM | DNA-PKcs | Activator | Chen et al. 2007 |
| ATM | MDC1 | Activator | Lou et al. 2006 |
| ATM_Control | CHK2 | Activator | Buscemi et al. 2004 |
| ATR | Fork_Stalling | Inhibitor | Paulsen and Cimprich. 2007 |
| Beta-Catenin | hTERT | Activator | Pestana et al. 2017 |
| Beta-Catenin | TCF | Activator | Uematsu et al. 2003 |
| Beta-TRCP | Beta-Catenin | Inhibitor | Fuchs et al. 2004 |
| BRCA1 | 53BP1 | Inhibitor | Feng et al. 2015 |
| BRCA1 | End_Resection | Activator | Cruz-Garcia et al. 2014 |
| BRCA1 | RNF8_HR | Inhibitor | Nakada et al. 2012 |
| BRCA2 | RAD51 | Activator | Luo et al. 2016 |
| c-Jun | BCL-2 | Inhibitor | Park et al. 1997 |
| c-Jun | MYC | Activator | Noguchi et al. 1999 |
| Caspase3 | Apoptosis | Activator | Tait and Green. 2010 |
| CDC25C | CycB_CDK1 | Activator | Liu et al. 2020 |
| CDK1 | CycB_CDK1 | Activator | Maddika et al. 2007 |
| CDK2 | CycE_CDK2 | Activator | Gudas et al. 1999 |
| CDK4 | CycD_CDK4 | Activator | Maddika et al. 2007 |
| CDK4 | TSC | Inhibitor | Romero-Pozuelo et al. 2020 |
| CDKN2A | ARF | Activator | Sherr. 2001 |
| CDKN2A | p16 | Activator | Chan et al. 2021 |
| CHK1 | CDC25C | Inhibitor | Uto et al. 2004 |
| CHK2 | CDC25C | Inhibitor | Uto et al. 2004 |
| CtIP | BRCA1 | Activator | Yun and Hiom. 2009 |
| CtIP | End_Resection | Activator | Cruz-Garcia et al. 2014 |
| CUL3 | KEAP1-CUL3-RBX1 | Activator | Thu et al. 2011 |
| CUL3 | LZTR1 | Activator | Bigenzahn et al. 2018 |
| CycB | CycB_CDK1 | Activator | Maddika et al. 2007 |
| CycD | CycD_CDK4 | Activator | Satyanarayana and Kaldis. 2009 |
| CycD_CDK4 | p21 | Inhibitor | Abukhdeir and Park. 2008 |
| CycD_CDK4 | p27 | Inhibitor | James et al. 2008 |
| CycE | CycE_CDK2 | Activator | Gudas et al. 1999 |
| CycE_CDK2 | G1_S_transition | Activator | Bertoli et al. 2013 |
| DNA-PKcs | NHEJ Ligation Complex | Activator | Davis et al. 2014 |

| From | To | Type | Citation |
|---|---|---|---|
| DSB | Ku | Activator | Trenner and Sartori. 2019 |
| DSB | Ku | Activator | Stucki et al. 2005 |
| DSB | MRN | Activator | Lamarche et al. 2010 |
| DSB | UrDSB | Activator | Waterman et al. 2020 |
| E2F | CycD | Activator | Guo et al. 2011 |
| E2F | CycE | Activator | Hwang and Clurman. 2005 |
| EGF | Erbb1 | Activator | Wieduwilt and Moasser. 2008 |
| EGF | Erbb1 | Activator | Yarden and Pines. 2012 |
| EGF | Erbb1 | Activator | Low-Nam et al. 2011 |
| EGF | Erbb1 | Activator | Purba et al. 2017 |
| EGF | Erbb1 | Activator | Jones et al. 1999 |
| EGFR | Beta-Catenin | Activator | Nakayama et al. 2014 |
| EGFR | GRB2 | Activator | Lowenstein et al. 1992 |
| EGFR | IL-6 | Activator | Ray et al. 2018 |
| EGFR | JAK2 | Activator | Olayioye et al. 1999 |
| EGFR | PI3K | Activator | Fumarola et al. 2014 |
| EGFR | PKA | Activator | Caldwell et al. 2012 |
| ELF3 | MEK | Activator | Wang et al. 2018 |
| ELF3 | PI3K | Activator | Wang et al. 2018 |
| End_Resection | PALB2 | Activator | Nepomuceno et al. 2017 |
| Erbb1 | EGFR | Activator | Wieduwilt and Moasser. 2008 |
| Erbb1 | Erbb3 | Activator | Sithanandam and Anderson. 2008 |
| Erbb3 | PI3K | Activator | Cook et al. 2011 |
| ERK | AMPK | Inhibitor | Hwang et al. 2013 |
| ERK | c-Jun | Activator | Leppa et al. 1998 |
| ERK | Caspase9 | Inhibitor | Allan et al. 2003 |
| ERK | CycD | Activator | Chambard et al. 2007 |
| ERK | Erbb3 | Activator | Lee et al. 2019 |
| ERK | MCL-1 | Activator | Song et al. 2005 |
| ERK | MYC | Activator | Lavoie et al. 2020 |
| ERK | NRF2 | Activator | DeNicola et al. 2001 |
| Excess_HR | Death | Activator | Appanah et al. 2020 |
| Fork_Stalling | DSB | Activator | Cortez. 2015 |
| FOXM1 | p21 | Activator | Tan et al. 2010 |
| FOXO3 | Bim | Activator | Meng et al. 2010 |
| Fyn | NRF2 | Inhibitor | Niture et al. 2011 |
| G2_Repair | UrDSB | Inhibitor | Arnoult et al. 2017; Paulsen et al. 2021 |
| GRB2 | MUC1 | Activator | Pandey et al. 1995 |
| GSK3B | 4EBP1 | Inhibitor | Ito et al. 2016 |
| GSK3B | Fyn | Activator | Jain and Jaiswal. 2007 |
| GSK3B | GSK3B-AXIN-CK1-APC | Activator | Pongracz and Stockley. 2006 |
| GSK3B | MCL-1 | Inhibitor | Fennel and Swanton. 2012 |
| GSK3B | p21 | Inhibitor | Robertson et al. 2017 |
| GSK3B | TSC | Activator | Evangelisti et al. 2020 |
| HGF | MET | Activator | Organ and Tsao. 2011 |
| HR | G2_Repair | Activator | Beucher et al. 2009 |
| hTERT | self_renewal | Activator | Leño et al. 2018 |
| IL-6 | IL-R6 | Activator | Uciechowski and Dempke. 2020 |
| IL-R6 | JAK2 | Activator | Ni et al. 2004 |
| IR | DSB | Activator | Vignard et al. 2013 |
| IR | SSB | Activator | Mahaney et al. 2009 |
| JAK2 | GRB2 | Activator | Bousoik and Montazeri Aliabadi. 2018 |
| JAK2 | NFkB | Activator | Funakoshi-Tago et al. 2011 |
| JAK2 | PI3K | Activator | Bousoik and Montazeri Aliabadi. 2018 |
| KEAP1 | KEAP1-CUL3-RBX1 | Activator | Thu et al. 2011 |
| KEAP1-CUL3-RBX1 | NFkB | Inhibitor | Thu et al. 2011 |
| KEAP1-CUL3-RBX1 | NRF2 | Inhibitor | Iso et al. 2016 |
| Ku | DNA-PKcs | Activator | Dobbs et al. 2010 |

| From | To | Type | Citation |
|---|---|---|---|
| KU70 | Ku | Activator | Rivera-Calzada et al. 2006 |
| KU80 | Ku | Activator | Rivera-Calzada et al. 2006 |
| LIG4 | NHEJ Ligation Complex | Activator | Ahnesorg et al. 2006 |
| LZTR1 | RAS | Inhibitor | Bigenzahn et al. 2018 |
| MAX | MAXMGA | Activator | Baudino and Cleveland. 2001 |
| MAX | MYCMAX | Activator | Baudino and Cleveland. 2001 |
| MAXMGA | MYCMAX | Inhibitor | Baudino and Cleveland. 2001 |
| MDC1 | RNF8 | Activator | Kolas et al. 2007 |
| MEK | CHK1 | Activator | Lee et al. 2017 |
| MET | Erbb3 | Activator | Engelman et al. 2007 |
| MET | GRB2 | Activator | Sacco and Clague. 2015 |
| MET | MUC1 | Activator | Singh et al. 2008 |
| MET | PI3K | Activator | Hervieu and Kermorgant. 2018 |
| MGA | MAXMGA | Activator | Baudino and Cleveland. 2001 |
| Mitosis | DSB | Activator | Haber. 2000 |
| Mitosis | Proliferation | Activator | Levine and Holland. 2018 |
| MOMP | Apoptosis | Activator | Tait and Green. 2010 |
| MRE11 | MRN | Activator | Lamarche et al. 2010 |
| MRN | ATM | Activator | Uziel et al. 2003 |
| MUC1 | EGFR | Activator | Piyush et al. 2017 |
| MUC1 | GRB2 | Activator | Singh and Bandyopadhyay. 2007 |
| MUC1 | GSK3B | Inhibitor | Huang et al. 2005 |
| MUC1 | MET | Inhibitor | Singh et al. 2008 |
| MUC1 | NFkB | Activator | Chaturvedi et al. 2011 |
| MUC1 | p53 | Activator | Singh et al. 2008 |
| MYC | MYCMAX | Activator | Baudino and Cleveland. 2001 |
| MYCMAX | ARF | Activator | Zindy et al. 1998 |
| MYCMAX | Bim | Activator | Muthalgu et al. 2014 |
| MYCMAX | CycD | Activator | García-Gutiérrez et al. 2019 |
| MYCMAX | hTERT | Activator | Khattar and Tergaonkar. 2017 |
| MYCMAX | NBS1 | Activator | Chiang et al. 2003 |
| MYCMAX | p15 | Inhibitor | Staller et al. 2001 |
| MYCMAX | PTEN | Activator | Kaur and Cole. 2013 |
| NBS1 | MRN | Activator | Lamarche et al. 2010 |
| Necrosis | Death | Activator | Golstein and Kroemer. 2007 |
| NFkB | c-Jun | Inhibitor | Eliseev et al. 2005 |
| NFkB | CycD | Activator | Hinz et al. 1999 |
| NHEJ Ligation Complex | UrDSB | Inhibitor | Dong et al. 2018 |
| NOXA | MCL-1 | Inhibitor | Zhang et al. 2011 |
| NRF2 | 53BP1 | Activator | Williams et al. 2017 |
| NRF2 | BCL-2 | Activator | Niture and Jaiswal. 2012 |
| NRF2 | BCL-XL | Activator | Niture and Jaiswal. 2013 |
| NRF2 | CycB_CDK1 | Activator | Murakami and Motohashi. 2015 |
| NRF2 | CycD_CDK4 | Activator | Shah et al. 2013 |
| NRF2 | CycD_CDK4 | Activator | Xie et al. 2021 |
| NRF2 | CycE_CDK2 | Activator | Kong et al. 2021 |
| p15 | CDK4 | Inhibitor | Sandhu et al. 1997 |
| p16 | CDK4 | Inhibitor | Kim and Sharpless. 2006 |
| p21 | STAT3 | Inhibitor | Coqueret and Gascan. 2000 |
| p53 | ARF | Inhibitor | Zeng et al. 2011 |
| p53 | NFkB | Inhibitor | Yang et al. 2015 |
| p53 | p21 | Activator | Shamloo and Usluer. 2019 |
| p53 | PTEN | Activator | Stambolic et al. 2001 |
| p90RSK | NFkB | Activator | Lin et al. 2019 |
| p90RSK | TSC | Inhibitor | Saxton. 2017 |
| PALB2 | BRCA2 | Activator | Zhang et al. 2009 |
| PKA | NF1 | Inhibitor | Feng et al. 2003 |

| From | To | Type | Citation |
|---|---|---|---|
| PTPRK | STAT3 | Inhibitor | Xu et al. 2019 |
| PTPRK | ZNRF3 | Activator | Chang et al. 2020 |
| PTPRT | STAT3 | Inhibitor | Zhang et al. 2007 |
| RAD50 | MRN | Activator | Lamarche et al. 2010 |
| RAD51 | HR | Activator | Godin et al. 2016 |
| RAF | MEK | Activator | Roberts and Der. 2007 |
| RAP80 | BRCA1 | Activator | Lu and Matunis. 2013 |
| RAS | NRF2 | Activator | DeNicola et al. 2011 |
| RAS | SKP2 | Activator | Lee et al. 2015 |
| RASSF1 | MDM2 | Inhibitor | Dubois et al. 2019 |
| RBX1 | KEAP1-CUL3-RBX1 | Activator | Thu et al. 2011 |
| RNF168 | RNF8 | Activator | Lu and Matunis. 2013 |
| RNF8 | 53BP1 | Activator | Lu and Matunis. 2013 |
| RNF8 | RNF8_HR | Activator | Nakada et al. 2012 |
| RNF8_HR | RAD51 | Activator | Nakada et al. 2012 |
| SFRP4 | WNT | Inhibitor | Ghoshal and Ghosh. 2016 |
| SKP2 | STK11 | Activator | Lee et al. 2015 |
| SSB | ATR | Activator | Maréchal and Zou. 2013 |
| SSB | Fork_Stalling | Activator | Alexander and Orr-Weaver. 2016 |
| STAT3 | BCL-2 | Activator | Weerasinghe et al. 2007 |
| STAT3 | BCL-XL | Activator | Weerasinghe et al. 2007 |
| STAT3 | CycB | Activator | Zhang et al. 2014 |
| STAT3 | CycD | Activator | Kang et al. 2019 |
| STAT3 | FOXM1 | Activator | Liao et al. 2018 |
| STAT3 | MCL-1 | Activator | Weerasinghe et al. 2007 |
| STAT3 | MUC1 | Activator | Gao et al. 2009 |
| STIL | Plk1 | Activator | Erez et al. 2007 |
| TCF | CycD | Activator | Klein and Assoian. 2008 |
| TCF | MET | Activator | Boon et al. 2002 |
| TCF | MYC | Activator | Dang. 2012 |
| TSC | mTORC1 | Inhibitor | Tian et al. 2019 |
| Twist | ARF | Inhibitor | Kwok et al. 2007 |
| Twist | p53 | Inhibitor | Piccinin et al. 2012 |
| UrDSB | ATM_Control | Activator | Ismail et al. 2005 |
| UrDSB | Necrosis | Activator | Higuchi. 2003 |
| WNT | Twist | Activator | Howe et al. 2003 |
| XRCC4 | NHEJ Ligation Complex | Activator | Ahnesorg et al. 2006 |
| ZNRF3 | Frizzled | Inhibitor | Chang et al. 2020 |

**Table S1. Regulatory edges in the *in silico* model.** Each row describes an edge, originating from the node in the “from” column and directed to the node in the “to” column.

**Table S2. Target functions used in the *in silico* model.**

| Node | Target Function | Citation | Reason |
|---|---|---|---|
| 14 3 3 | avg(pos)-avg(neg) | Qi et al., 2003 | Sequesters Cdc25c in cytoplasm and inhibits its activity |
| 4EBP1 | 3-mTORC1 | Wu et al., 2019 | Inhibted by mTORC1 |
| 53BP1 | RNF8-BRCA1 + NRF2 -1 | Hu et al., 2014 Chapman et al., 2012 Williams et al., 2017 Yang et al., 2021 | Activated by RNF8 and NRF2 inhibited by BRCA1 |
| Akt | ceil(avg(PI3K , mTORC2)) - max(0 , AMPK-1) | Guo et al., 2013 Xia et al., 2021 | Activated by PI3K and mTORC2 inhibited by AMPK |
| AMPK | STK11 - max(ERK , Akt) | Thoreen and Sabatini 2005 Hahn Windgassen et al., 2005 | Activated by STK11 dowregulated by ERK and or Akt |
| APC | 1 | Kimelman and Xu 2006 Xing et al., 2003 | Contributes to GSK3B AXIN CK1 APC complex |
| Apoptosis | ceil(avg(MOMP , Caspase3)) | Tait et al., 2010 Brentnall et al., 2013 | Activated by MOMP and Caspase 3 |
| ARF | (max(0 , MYCMAX-2) + max(0 , E2F-2)) * min(1 , CDKN2A) - (Twist-2) | Komori et al., 2005 Zindy et al., 1998 Kwok et al., 2007 | Encoded by CDKN2A Activated by overexpression of MYCMAX and ARF inhibited by Twist |
| ATM | MRN*max(0 , min(1 , 1-ATM_deficient))+min(1 , ATM_deficient*MRN) | | |
| ATM Control | ATM-(2-(UrDSB)) | Buscemi et al., 2004 Ismail et al., 2005 | ATM is activated by DNA double strand breaks |
| ATM deficient | avg(pos)-avg(neg) | | |
| ATR | avg(pos)-avg(neg) | Mar chal and Zou 2013 | |
| BAD | 3-ceil(avg(Akt , p90RSK)) | Datta et al., 1997 Hurbin et al., 2005 | Inhibited by Akt and p90RSK |
| BAKBAX | 3 - ceil(avg(BCL-2 , BCL-W , BCL-XL , MCL-1)) | Fennel and Swanton 2012 Kale et al., 2018 | Inhibited by BCL 2 BCL W BCL XL and MCL 1 |

| Node | Target Function | Citation | Reason |
|---|---|---|---|
| BCL 2 | (2 - ceil(avg(BAD , Bim , PUMA)) + min(NFkB , min(NRF2 , STAT3)) - max(0 , 1-c-Jun)) * max(1 , (c-Jun-1)) | Chen et al., 2005 Liu et al., 2015 Chen et al., 2017 Kang et al., 2019 Park et al., 1997 | Activated by BIM BAD PUMA inhibited by NFkB NRF2 STAT3 and c Jun |
| BCL W | 3-floor(avg(Bim , PUMA)) | Chen et al., 2005 Zhang et al., 2013 | Inhibited by BIM and PUMA |
| BCL XL | 2 - ceil(avg(BAD , Bim , PUMA)) + floor(avg(NFkB , NRF2 , Beta-Catenin , STAT3)) | Chen et al., 2005 Chen et al., 2017 Niture and Jaiswal 2013 Zhang et al., 2013 | Activated by NFkB NRF2 Beta Catenin and STAT3 inhibited by BAD Bim and PUMA |
| Beta Catenin | ((2-GSK3B-AXIN-CK1-APC) + Akt + max(0 , EGFR-2)) * (3-Beta-TRCP) | MacDonals and He 2012 Zhang and Wang 2020 Nakayama et al., 2014 Fuchs et al., 2004 | Activated by GSK3B AXIN CK1 APC inhibited by Akt EGFR and Beta TRCP |
| Beta TRCP | 2 * min(max(GSK3B , 0) , 1) | Paul and Dey 2007 | Inhibits Beta Catenin GSK 3 which can phosphorylate TrCP increasing NRF2 ubiquitination |
| Bim | 3 - ceil(avg(Akt , ERK+1)) + floor(avg(MYCMAX , FOXO3)) | Muthalagu et al., 2014 Meng et al., 2010 Takezawa et al., 2011 Qi et al., 2006 | Activated by MYCMAX and FOXO3 inhibited by Akt and ERK |
| BRCA1 | max(CtIP , RAP80) * min(1 , CtIP) | Nakamura et al., 2010 Hu et al., 2011 | Activated by CtIP and RAP80 CtIP function essential for activity |
| BRCA2 | avg(pos)-avg(neg) | Zhang et al., 2009 | Activated by PALB2 |
| c Jun | floor(avg(Akt , ERK)) * max (1 , (2-NFkB)) | Nguyen et al., 2004 Zhao et al., 2015 Eliseev et al., 2005 | Activated by Akt ERK and NFkB |
| Caspase3 | avg(pos)-avg(neg) | | |
| Caspase9 | ceil(MOMP - floor(avg(Akt , ERK , CycB_CDK1))) | Kalkavan and Green 2018 Allan and Clarke 2007 Cardone et al., 1998 Allan et al., 2003 | Activated by MOMP inhibited by Akt ERK and Cyclin B CDK1 |
| CBF | 3-NCID | | |

| Node | Target Function | Citation | Reason |
|---|---|---|---|
| CDC25A | ceil(avg(E2F , Plk1 , MYCMAX)) - ceil(avg(CHK1 , CHK2)) | Karlsson Rosenthal and Millar 2006 Vigo et al., 1999 Z rnig and Evan 1996 Mailand et al., 2002 | Activated by E2F Plk1 and MYCMAX inhibited by CHK1 and CHK2 |
| CDC25C | 1+(MYCMAX)-floor(avg(14_3_3 , ATM , CHK1 , CHK2)) | | |
| CDK1 | 3-Wee1 | Ghelli Luserna di Ror et al., 2020 | Inhibited by Wee1 |
| CDK2 | 3-avg(p21 , Wee1) | Al Bitar and Gali Muhtasib 2019 Otto and Sicinski 2017 | Inhibited by p21 and Wee1 |
| CDK4 | 3-(max(p15 , p16)) | Bockstaele et al., 2006 | Inhibited by p15 and p16 |
| CDKN2A | 2 | Chan et al., 2021 | Protein encoding gene |
| CHK1 | max(ATR , floor(avg(ATR , MEK-1))) | Lee et al., 2017 | Activated by ATR Activity potentiated by high MEK activity |
| CHK2 | avg(pos)-avg(neg) | | |
| CtIP | avg(pos)-avg(neg) | Han et al., 2021 | Activated by ATM |
| CUL3 | 2 | Mikac et al., 2021 | Constitutively active unless mutated |
| CycB | ceil(avg(FOXM1 , STAT3)) - FOXO3 | Xu et al., 2012 | Activated by FOXM1 and STAT3 inhibited by FOXO3 |
| CycB CDK1 | ceil(avg(CycB , CDK1 , CDC25A , NRF2) - max(0 , p21-2)) * min(1 , CDC25C) * min(1 , CycB) * min(1 , CDK1) * min(1 , CDC25A) | Petri et al., 2007 Timofeev et al., 2010 Murakami and Motohashi 2015 Kreis et al., 2015 | Activated by NRF2 Cyclin B and CDK1 inhibited by p21 |
| CycD | ceil(avg(YB1 , NFkB , TCF , ERK , STAT3)) - floor(avg(GSK3B , 4EBP1)) + MYCMAX | Harada et al., 2014 Hinz et al., 1999 Klein and Assoian 2008 Lim et al., 2006 Garc a Guti rrez et al., 2019 Chambard et al., 2007 Kang et al., 2019 Barnhart et al., 2008 Yu et al., 2012 | Activated by YB1 NFkB TCF MYCMAX ERK and STAT3 inhibited by GSK3B 4EBP1 |

| Node | Target Function | Citation | Reason |
|---|---|---|---|
| CycD CDK4 | floor(avg(CycD , CDK4 , NRF2) - avg(p27 , p21)) * min(1 , min(CycD , CDK4)) | Datar et al., 2006 Xie et al., 2021 Shah et al., 2013 Naruse et al., 2000 He et al., 2005 | Activated by NRF2 Cyclin D and CDK4 inhibited by p27 and p21 |
| CycE | avg(pos)-avg(neg) | | |
| CycE CDK2 | ceil(avg(min(CycE , CDK2) * min(1 , CDC25A) , NRF2)) - (avg(max(0 , p27-1) , max(0 , p21-1))) | Shen and Huan 2012 Gudas et al., 1999 He et al., 2005 Sheaff et al., 1997 | Activated by NRF2 Cyclin E CDK2 CDC25A inhibited by p27 and p21 |
| Death | (max(max(Apoptosis , Necrosis) , toxic_NHEJ)) + max(0 , min(1 , Excess_HR-1)) | Golstein and Kroemer 2007 Roos and Kaina 2006 Appanah et al., 2020 | Cell death may be apoptotic or necrotic Excess HR is toxic to the cell |
| Dishevelled | avg(Frizzled , LRP) *min(1 , LRP)*min(1 , Frizzled) | Cong et al., 2004 | Activated by LRP and Frizzled whic are both essential for activity |
| DNA PKcs | max(ceil(avg((ATM-1) , Ku)) , Ku) * min(1 , Ku) | Dobbs et al., 2010 Chen et al., 2007 | Requires presence of Ku for activity degree of activity depends on ATM and Ku activities |
| DSB | max(max(avg(Mitosis , IR) , avg(Fork_Stalling , IR)) , IR) | Barnard et al., 2013 Crasta et al., 2012 Cortez 2015 | IR induces high levels of DSBs and routinely occur during mitosis Fork stalling leads to DNA DSBs |
| E2F | 3-pRB | Giacinti and Giordano 2006 | Inhibited by pRB |
| EGF | 2 | Wells 1999 | Activated by EGFR |
| EGFR | Erbb1 * (min(1 , MUC1) * max(1 , (MUC1-1))) | Purba et al., 2017 Kharbanda et al., 2014 | Activated by Erbb1 and MUC1 MUC1 function essential for activity |
| ELF3 | 1 | Wang et al., 2018 | Activated by RAS and PI3K |
| End Resection | max(BRCA1 , avg(CtIP , BRCA1)-53BP1) | Cruz Garc a et al., 2014 Zimmermann et al., 2013 | Activated by CtIP and BRCA1 inhibited by 53BP1 |
| Erbb1 | avg(pos)-avg(neg) | | |

| Node | Target Function | Citation | Reason |
|---|---|---|---|
| Erbb3 | ((max(0 , (1-ERK))*max(0 , min(1 , Erbb1-2)) + max(0 , (1-Erbb1))*max(0 , min(1 , MET-2))))*3 | Engelman et al., 2007 Sithanandam and Anderson 2008 Lee et al., 2019 | Activated by ERK MET and EGFR |
| ERK | MEK | Kalkavan and Green 2018 Yao et al., 2011 Marshall 1994 Lee et al., 2017 | MEK is essential for ERK activity ERK is further activated by MUC1 |
| Excess HR | min(End_Resection , 3-max(0 , min(1 , max(BRCA1 , BRCA2)))) | | |
| Fork Stalling | SSB-ATR | Alexander and Orr Weaver 2016 Paulsen and Cimprich 2007 | Replication fork stalling is caused by DNA single strand breaks ATR prevents fork collapse and promotesrestart of replication fork preogression |
| FOXM1 | floor(avg(CycD_CDK4 , E2F , MYCMAX , STAT3)) - pRB | Liao et al., 2018 VanArsdale et al., 2015 | Activated by CyclinD CDK4 E2F MYCMAX and STAT3 inhibited by pRB |
| FOXO3 | 3 - ceil(avg(Akt , ERK)) + min(1 , min(AMPK , p53)) | Renault et al., 2011 Wang et al., 2017 Chiacchiera and Simone 2010 Liu et al., 2018 | Activated by AMPK and p53 inhibited by Akt and ERK |
| Frizzled | WNT-max(0 , ZNRF3-1) | Chang et al., 2020 | Activated by Wnt inhibited by ZNRF3 |
| Fyn | avg(pos)-avg(neg) | | |
| G1 S transition | avg(pos)-avg(neg) | Bertoli et al., 2013 | Activated by Cyclin E CDK2 |
| G1arrest | avg(pos)-avg(neg) | Park et al., 2001 | Inhibited by p27 |
| G2 M arrest | 3 - CycB_CDK1 | Castedo et al., 2002 | Inhibited by CyclinB CDK1 |
| G2 Repair | avg(pos)-avg(neg) | Truong et al., 2013 | HR and MMEJ mediate DNA double strand break repair in S and G2 phases of the cell cycle |

| Node | Target Function | Citation | Reason |
|---|---|---|---|
| GRB2 | ceil(avg(EGFR , JAK2 , MET , MUC1)) | Sacco and Clague 2015 Bousoik and Montazeri Aliabadi 2018 Bidkhori et al., 2012 | Activated by EGFR JAK2 MET and MUC1 |
| GSK3B | 3 - ceil(avg(Akt , ERK , MUC1 , p70S6K-1 , p90RSK)) | | Inhibited by Akt ERK MUC1 p70S6K and p90RSK |
| GSK3B AXIN CK1 APC | (3-Dishevelled) * min(GSK3B , APC) | Paul and Dey 2008 Huang et al., 2005 | Inhibited by Dishevelled APC and GSK3B are key members of this protein complex |
| HGF | 2 | Organ and Tsao 2011 | Activated by MET |
| HR | avg(pos)-avg(neg) | Budke et al., 2012 | Mediates homologous recombination |
| hTERT | ceil(avg(Akt , Beta-Catenin , MYCMAX)) | Daniel et al., 2012 Pestana et al., 2017 Khattar and Tergaonkar 2017 | Activated by Akt Beta Catenin and MYCMAX |
| Hypoxia | 2 | | |
| IL 6 | 2 + max(0 , EGFR-2) | Ray et al., 2018 | Constitutive expression High EGFR activity elevates expression |
| IL R6 | avg(pos)-avg(neg) | | |
| IR | 3-IR_FLIP | Barnard et al., 2013 | DSBs are induced by IR |
| IR FLIP | 3 | | |
| JAK2 | floor(avg(EGFR , IL-R6)) + max(0 , IL-R6-2) + max(0 , EGFR-2) | Johnson et al., 2018 Wee and Wang 2017 | Independent activation by EGFR and IL R6 |
| KEAP1 | 2 | Nguyen et al., 2009 | Constituively activated unless mutated |
| KEAP1 CUL3 RBX1 | min(RBX1 , min(CUL3 , KEAP1)) | Thu et al., 2011 | Ubiquitin ligase complex composed by KEAP1 CUL3 and RBX1 |
| Ku | avg(max(DSB , seDSB) , 53BP1) * min(1 , KU70)*min(1 , KU80) | | Components of Ku complex are Ku70 and KU80 Activated by DSBs and 53BP1 |
| KU70 | 1 | Mari et al., 2006 | Component of the Ku heterodimer |

| Node | Target Function | Citation | Reason |
|---|---|---|---|
| KU80 | 1 | Mari et al., 2006 | Component of the Ku heterodimer |
| LIG4 | 1 | Gu et al., 2007 | Component of the NHEJ Ligation Complex |
| LRP | avg(pos)-avg(neg) | | |
| LZTR1 | CUL3-1 | Bigenzahn et al., 2018 | Activated by CUL3 |
| MAX | 3 | Kato et al., 1992 | Constitutively active unless mutated |
| MAXMGA | min (MGA , MAX) | Diolaiti et al., 2015 | Activated by MAX and MGA requires function of both these proteins for activity |
| MCL 1 | 2 - ceil(avg(Bim , NOXA , PUMA , GSK3B)) + min(min(Akt , ERK) , STAT3) | Fennel and Swanton 2012 Zhang et al., 2011 Song et al., 2005 Kang et al., 2017 Weerasinghe et al., 2007 | Activated by Akt ERK and STAT3 inhibited by Bim NOXA PUMA and GSK3B |
| MDC1 | max(DSB , seDSB) * min(1 , ATM+max(0 , min(1 , STAT3-2)))*min(1 , ATM) | Eliezer et al., 2014 | ATM mediated phosphorylation of H2AX at double strand breaks is essential for MDC1 activation |
| MDM2 | 3 - ceil(avg(ARF , RASSF1)) | Nag et al., 2016 Song et al., 2008 | Inhibited by ARF and RASSF1 |
| MEK | RAF-max(0 , min(1 , 1-ELF3)) | Roberts and Der 2007 Wang et al., 2018 | Activated by RAF inhibited by ELF3 |
| MET | HGF +(min(1 , TCF)-1) | Raghav et al., 2012 Singh et al., 2008 Boon et al., 2002 | Activated by HGF and TCF inhibited by MUC1 |
| MGA | 1 | Baudino and Cleveland 2001 | Inhibits MYCMAX through the MAXMGA heterodimer |
| Mitosis | ceil(avg(CycB_CDK1 , Plk1)) | Castedo et al., 2002 Gheghiani et al., 2017 | Activated by CyclinB CDK1 and Plk1 |
| MOMP | avg(pos)-avg(neg) | | |
| MRE11 | 1 | Langerak et al., 2011 | Component of the MRN complex |

| Node | Target Function | Citation | Reason |
|---|---|---|---|
| MRN | min((MRE11) , min (NBS1 , RAD50)) * max(DSB , floor(avg(DSB , 53BP1))) | Uziel et al., 2003 Lee et al., 2010 | Composed of MRE11 NBS1 and RAD50 primarily activated by DSBs and also by 53BP1 |
| mTORC1 | 3-TSC | Tian et al., 2019 | Inhibited by TSC |
| mTORC2 | avg(pos)-avg(neg) | | |
| MUC1 | floor(avg(STAT3 , GRB2 , MET , 1)) | Gao et al., 2009 Li et al., 2001 Singh et al., 2008 | Activated by STAT3 GRB2 and MET |
| MYC | ceil(avg(ERK , TCF , c-Jun)) - ceil(avg(GSK3B , 4EBP1))*min(2 , max(1 , MycAmp)) | Duffy et al., 2021 Robertson et al., 2018 Lavoie et al., 2020 Barnhart et al., 2008 Yochum et al., 2009 Noguchi et al., 1999 | Activated by ERK STAT3 and B catenin inhibited by FOXO3 p53 GSK3B and 4EBP1 |
| MycAmp | avg(pos)-avg(neg) | | |
| MYCMAX | min(MYC , MAX) * min(2 , 3-(MAXMGA)) | Baudino et al., 2001 | Requires both MYC and MAX activity antagonised by MAXMGA |
| NBS1 | 1 *min(1 , MYCMAX) | Kobayashi et al., 2004 Chiang et al., 2003 | Component of the MRN complex MYCMAX function essential for activity |
| NCID | avg(pos)-avg(neg) | | The ultimate event in Notch multiple proteolytic processing leads to the release in the nucleus of the activated form of Notch intracellular Notch or NotchIC |
| Necrosis | max(UrDSB , min(1 , UrseDSB)) | | Unrepaired DNA Double Strand Breaks result in Necrotic death |
| NF1 | 3-PKA | Bergoug et al., 2020 | PKA downregulates NF1 expression |

| Node | Target Function | Citation | Reason |
|---|---|---|---|
| NFkB | ceil(avg(Akt , p90RSK , JAK2 , MUC1))-floor(avg(p53 , KEAP1-CUL3-RBX1)) | Lin et al., 2019 Yang et al., 2015 Chaturvedi et al., 2011 Kucuksayan et al., 2016 | Activated by Akt p90RSK JAK2 and MUC1 inhibited by p53 and KEAP1 |
| NHEJ Ligation Complex | DNA-PKcs * min(min(1 , LIG4) , min(XRCC4 , XLF)) | Davis et al., 2014 Ahnesorg et al., 2006 | Protein complex consists of XRCC4 XLF4 and LIG4 Activated by DNA PKcs |
| NOTCH1 | avg(pos)-avg(neg) | | NOTCH1 only active in NSCLC in hypoxic conditions |
| NOXA | avg(pos)-avg(neg) | | |
| NRF2 | 3 - floor(avg(Fyn , KEAP1-CUL3-RBX1*2 , Beta-TRCP)) + floor(avg(ERK , Akt)) | Barrera Rodr guez 2018 Sparaneo et al., 2016 | Activated by ERK and Akt inhibited by Fyn and KEAP1 Another regulator of NRF2 is glycogen synthase kinase 3 beta GSK 3 This protein can phosphorylate TrCP increasing the ability of TrCP to ubiquitinate NRF2 |
| p15 | 3-MYCMAX | Garc a Guti rrez et al., 2019 | Inhibited by MYCMAX |
| p16 | CDKN2A - YB1 | Chan et al., 2021 | Inhibtied by YB1 Encoded by CDKN2A |
| p21 | ceil(avg(NRF2 , FOXO3 , FOXM1) + p53) - floor(avg(max(MYCMAX-1 , 0) , CycD_CDK4 , Akt , NRF2 , GSK3B)) | Lee et al., 2008 Li et al., 2018 Garc a Guti rrez et al., 2019 Robertson et al., 2017 Tan et al., 2010 | Activated by p53 FOXO3 and FOXM1 inhibited by MYCMAX CycD CDK4 Akt NRF2 and GSK3B |

| Node | Target Function | Citation | Reason |
|---|---|---|---|
| p27 | (1 + FOXO3 - floor(avg(MYCMAX , Akt , p90RSK , CycE_CDK2 , NRF2))) | Chikkegowda et al., 2021 Garc a Guti rrez et al., 2019 Nho and Hergert 2014 Sherr and Roberts 1999 Rao et al., 2017 Poomakkoth et al., 2016 | Activated by FOXO3 inhibited by MYCMAX Akt p90RSK Cyclin E CDK2 and NF2 |
| p27 amp | 1 | | |
| p53 | 3 - ceil(avg(MDM2 , Twist , NCID)) + min(1 , (min(min(CHK1 , CHK2) , min(AMPK , MUC1)))) | Thoreen and Sabatini 2005 Ch ne 2003 Yang et al., 2005 Taylor and Stark 2001 Piccinin et al., 2012 | Activated by AMPK CHK1 CHK2 and MUC1 inhibited by MDM2 and Twist It is inhibited by NOTCH1 through the modulation of its stability at the posttransla-tional level |
| p70S6K | avg(pos)-avg(neg) | | |
| p90RSK | avg(pos)-avg(neg) | | |
| PALB2 | avg(pos)-avg(neg) | Zhang et al., 2009 | Activated by BRCA1 |
| PI3K | floor(avg(EGFR , EGFR , MET , JAK2 , RAS , MUC1) - PTEN + Erbb3) * min(1 , PIK3CA) + max(ELF3-1 , 0) | Bidkhori et al., 2012 Martinez Mart and Felip 2012 Hervieu and Kermorgant 2018 Bousoik and Montazeri Aliabadi 2018 Shaw and Cantley 2006 Engelman et al., 2007 Raina et al., 2011 Hao et al., 2021 | Activated by EGFR MET JAK2 MUC1 IRS4 Erbb3 and RAS and inhibited by PTEN PIK3CA encodes the catalytic subunit of PI3K and its function is essential to the activity of PI3K |
| PIK3CA | 1 | Martinez Mart and Felip 2012 | Encodes catalytic subunit of PI3K |
| PKA | avg(pos)-avg(neg) | Caldwell et al., 2012 | Activated by Erbb1 |
| Plk1 | 2 -ceil(avg(p53 , FOXO3))+ floor(avg(STIL , FOXM1)) | McKenzie et al., 2010 Costa 2005 Bucur et al., 2014 Erez et al., 2007 Castiel et al., 2011 | Activated by FOXM1 and STIL inhibited by p53 FOXO3 p21 |
| pRB | 2 - min(CycE_CDK2 , CycD_CDK4) | VanArsdale et al., 2015 | Inibitied by CycE CDK2 and CycD CDK4 |

| Node | Target Function | Citation | Reason |
|---|---|---|---|
| Proliferation | avg(pos)-avg(neg) | | |
| PTEN | floor(avg(p53 , MYCMAX)) - floor(avg(CBF , c-Jun)) | Stambolic et al., 2001 Hettinger et al., 2007 | Activated by p53 and MYCMAX inhibited by c Jun Notch signalling downregulates the expression of PTEN via the induction of hairy enhancer of split 1 |
| PTPRK | 2 | Xu et al., 2019 Chang et al., 2020 | Activates ZNRF3 inhibits STAT3 |
| PTPRT | 1 | Zhang et al., 2007 | Activated by ho-modimerisation activates STAT3 |
| PUMA | max(FOXO3 , p53) *min(1 , p53) | Nakano and Vousden 2001 You et al., 2006 Wang et al., 2007 | Activated by FOXO3 and p53 p53 function essential for activity |
| RAD50 | 1 | Dupr et al., 2006 | Component of the MRN complex |
| RAD51 | floor(min(BRCA2 , RNF8_HR)) | Esashi et al., 2007 | Activated by BRCA2 and RNF8 |
| RAD51 | floor(min(BRCA2 , RNF8_HR)) | Esashi et al., 2007 Nakada et al., 2012 | Activated by BRCA2 and RNF8 |
| RAF | avg(pos)-avg(neg) | Hao et al., 2021 | Ras essential for activity Activity further potentiated by IRS4 |
| RAP80 | avg(pos)-avg(neg) | Strauss and Goldberg 2011 | Recruited by MDC1 to DNA double strand breaks |
| RAS | SOS - max(0 , (avg(NF1 , LZTR1)-1)) | Boriack Sjodin et al., 1998 Johnson et al., 1994 | Activated by SOS inhibited by NF1 |
| RASSF1 | avg(pos)-avg(neg) | Palakurthy et al., 2009 | Inhibited by DNMT3B |
| RBX1 | 2 | Wei and Sun 2010 | Constitutively active unless mutated |
| RNF168 | 1 | Chen et al., 2021 | Component of the RNF8 RNF168 ubiquitin ligase complex |

| Node | Target Function | Citation | Reason |
|---|---|---|---|
| RNF8 | MDC1*min(1 , RNF168) | Huen et al., 2007 Bartocci and Denchi 2013 | Activated by MDC1 acts in concurrence with RNF168 to promote accumulation of 53BP1 |
| RNF8 HR | RNF8 - max(0 , 2*(1- min(1 , BRCA1) - min(1 , 53BP1))) | Nakada et al., 2012 | Both BRCA1 and 53BP1 must be absent for RNF8 to promote homologous recombination via RAD51 |
| seDSB | avg(pos)-avg(neg) | | |
| self renewal | avg(pos)-avg(neg) | | |
| Senescence | avg(p21 , p16) *min(1 , p21)*min(1 , p16) | | |
| SFRP4 | 1 | Lee et al., 2008 | Inhibits Wnt |
| SKP2 | avg(pos)-avg(neg) | Lee et al., 2015 | Activated by Ras |
| SOS | avg(pos)-avg(neg) | | |
| SSB | max(ceil(avg(Mitosis , IR)) , IR) | Mahaney et al., 2009 Crasta et al., 2012 | Induced by ionising radiation and routinely occurs during mitosis |
| STAT3 | ((floor(avg(JAK2 , MET , EGFR)) + max(0 , JAK2-2))) + max(0 , 1-PTPRT) + max(0 , 1-PTPRK) | Xu et al., 2019 | Activated by JAK2 and MET inhibited by PTPRK The STAT3 transcription factor binds to activated MET through phosphorylated tyrosine residue Y1356 of MET |
| STIL | 1 | Erez et al., 2007 Castiel et al., 2011 | Activates Plk1 |
| STK11 | max(2 , SKP2) | Choi et al., 2015 Lee et al., 2015 | Constituively activated unless mutated Maximally activated by high SKP2 activity |
| TCF | avg(pos)-avg(neg) | | |
| toxic NHEJ | NHEJ_Ligation_Complex*min(1 , UrseDSB)-G2_Repair | | |

| Node | Target Function | Citation | Reason |
|---|---|---|---|
| TSC | max(GSK3B , AMPK) - floor(avg(ERK , Akt , Akt , Akt , p90RSK)) + 1 | Corradetti et al., 2004 Vicary and Roman 2016 Tian et al., 2019 Saxton 2017 Evangelisti et al., 2020 Ma et al., 2005 | Activated by GSK3B and AMPK inhibited by ERK Akt and p90RSK |
| Twist | avg(pos)-avg(neg) | Howe et al., 2003 | Activated by Wnt inhibited by ZNRF3 |
| UrDSB | DSB-floor(avg(G2_Repair , NHEJ_Ligation_Complex)) | Mao et al., 2008 Seol et al., 2018 | Unrepaired DNA double strand breaks are repaird by HR NHEJ and MMEJ |
| UrseDSB | avg(pos)-avg(neg) | | |
| Wee1 | max(3-Plk1 , floor(avg(CHK1 , CHK2))) | Watanabe et al., 2004 | Activated by Plk1 CHK1 and CHK2 |
| WNT | 3-SFRP4 | Lee et al., 2008 | Inhibited by SFRP4 |
| XLF | 1 | | |
| XRCC4 | 1 | Koch et al., 2004 | Component of the NHEJ Ligation Complex |
| YB1 | avg(pos)-avg(neg) | | |
| ZNRF3 | avg(pos)-avg(neg) | Chang et al., 2020 | Inhibits Frizzled |

**Table S2. Target functions used in the *in silico* model.** The target function for each node is specified, along with a brief description of the mechanism and a reference where required. Notes on the syntax of the target functions are available from https://biomodelanalyzer.org/. A number in the Target Function column represents nodes that are set to a constant value.

**Table S3. Summary of the nodes included in the model**

| Model node name | Gene symbol | Full protein name |
|---|---|---|
| 14-3-3 | YWHAZ | 14-3-3 protein |
| 4EBP1 | EIF4EBP1 | 4E-Binding Protein 1 |
| 53BP1 | TP53BP1 | p53 Binding Protein 1 |
| Akt | AKT1 | Protein Kinase B |
| AMPK | PRKAA1/PRKAA2 | AMP-activated protein kinase |
| APC | APC | Adenomatous Polyposis Coli |
| ARF | CDKN2A | ADP Ribosylation Factor |
| ATM | ATM | Ataxia Telangiectasia Mutated |
| ATM-Control | ATM | Ataxia Telangiectasia Mutated (Control) |
| ATM-deficient | ATM | Ataxia Telangiectasia Mutated (Deficient) |
| ATR | ATR | Ataxia Telangiectasia and Rad3 related |
| BAD | BAD | Bcl-2-associated death promoter |
| BAKBAX | BAK1/BAX | Bcl-2 homologous antagonist killer/Bcl-2-associated X protein |
| baseline | | Baseline |
| BCL-2 | BCL2 | B-cell lymphoma 2 |
| BCL-W | BCL2L2 | Bcl-2-like protein 2 |
| BCL-XL | BCL2L1 | B-cell lymphoma-extra large |
| Beta-Catenin | CTNNB1 | Beta-Catenin |
| Beta-TRCP | BTRC | Beta-transducin repeat-containing protein |
| Bim | BCL2L11 | Bcl-2-like protein 11 |
| BRCA1 | BRCA1 | Breast cancer type 1 susceptibility protein |
| BRCA2 | BRCA2 | Breast cancer type 2 susceptibility protein |
| c-Jun | JUN | Jun proto-oncogene |
| CBF | RUNX1 | Core-binding factor subunit alpha-1 |
| CDC25A | CDC25A | Cell division cycle 25A |
| CDC25C | CDC25C | Cell division cycle 25C |
| CDK1 | CDK1 | Cyclin-dependent kinase 1 |
| CDK2 | CDK2 | Cyclin-dependent kinase 2 |
| CDK4 | CDK4 | Cyclin-dependent kinase 4 |
| CDKN2A | CDKN2A | Cyclin-dependent kinase inhibitor 2A |
| CHK1 | CHEK1 | Checkpoint kinase 1 |
| CHK2 | CHEK2 | Checkpoint kinase 2 |
| CtIP | RBBP8 | RBBP8 (CtIP) |
| CUL3 | CUL3 | Cullin 3 |
| CycB | CCNB1 | Cyclin B1 |
| CycD | CCND1 | Cyclin D1 |
| CycE | CCNE1 | Cyclin E1 |
| Dishevelled | DVL1/DVL2/DVL3 | Dishevelled segment polarity protein |
| DNA-PKcs | PRKDC | DNA-dependent protein kinase catalytic subunit |
| E2F | E2F1 | E2F transcription factor 1 |
| EGF | EGF | Epidermal growth factor |
| EGFR | EGFR | Epidermal growth factor receptor |
| ELF3 | ELF3 | E74-like factor 3 |
| Erbb1 | ERBB1 | Erb-b2 receptor tyrosine kinase 1 |
| Erbb3 | ERBB3 | Erb-b2 receptor tyrosine kinase 3 |
| ERK | MAPK1/MAPK3 | Extracellular signal-regulated kinase |
| FOXM1 | FOXM1 | Forkhead box protein M1 |
| FOXO3 | FOXO3 | Forkhead box protein O3 |
| Frizzled | FZD1-FZD10 | Frizzled class receptor |
| Fyn | FYN | Fyn proto-oncogene |
| GRB2 | GRB2 | Growth factor receptor-bound protein 2 |

| Model node name | Gene symbol | Full protein name |
|---|---|---|
| GSK3B-AXIN-CK1-APC | GSK3B/AXIN1/CSNK1A1/APC | Glycogen synthase kinase 3 beta/Axin 1/Casein kinase 1 alpha 1/Adenomatous polyposis coli |
| GSK3B | GSK3B | Glycogen synthase kinase 3 beta |
| HGF | HGF | Hepatocyte growth factor |
| hTERT | TERT | Telomerase reverse transcriptase |
| IL-6 | IL6 | Interleukin 6 |
| IL-R6 | IL6R | Interleukin 6 receptor |
| IR | INSR | Insulin receptor |
| IR-FLIP | | Irradiation flipping node |
| JAK2 | JAK2 | Janus kinase 2 |
| KEAP1-CUL3-RBX1 | KEAP1/CUL3/RBX1 | Kelch-like ECH-associated protein 1/Cullin 3/Ring-box 1 |
| KEAP1 | KEAP1 | Kelch-like ECH-associated protein 1 |
| KU70 | XRCC6 | X-ray repair cross-complementing protein 6 |
| KU80 | XRCC5 | X-ray repair cross-complementing protein 5 |
| LIG4 | LIG4 | DNA ligase 4 |
| LRP | LRP1 | LDL receptor-related protein 1 |
| LZTR1 | LZTR1 | Leucine-zipper-like transcription regulator 1 |
| MAX | MAX | Myc-associated factor X |
| MCL-1 | MCL1 | Induced myeloid leukemia cell differentiation protein |
| MDC1 | MDC1 | Mediator of DNA damage checkpoint 1 |
| MDM2 | MDM2 | Mouse double minute 2 homolog |
| MEK | MAP2K1/MAP2K2 | Mitogen-activated protein kinase kinase 1/2 |
| MET | MET | MET proto-oncogene |
| MGA | MGA | MAX gene-associated |
| MOMP | BAX/BAK1 | Mitochondrial outer membrane permeabilization (Bcl-2 homologous antagonist killer/Bcl-2-associated X protein) |
| MRE11 | MRE11 | MRE11 homolog |
| MRN | MRE11/RAD50/NBN | MRE11-RAD50-NBS1 complex |
| mTORC1 | MTOR | Mechanistic target of rapamycin complex 1 |
| mTORC2 | MTOR | Mechanistic target of rapamycin complex 2 |
| MUC1 | MUC1 | Mucin 1 |
| MYC | MYC | Myc proto-oncogene |
| MycAmp | MYC | Myc proto-oncogene amplification |
| NBS1 | NBN | Nijmegen breakage syndrome 1 |
| NCID | NOTCH1 | Notch intracellular domain |
| NF1 | NF1 | Neurofibromin 1 |
| NFkB | NFKB1/NFKB2/RELA | Nuclear factor kappa-light-chain-enhancer |
| NOTCH1 | NOTCH1 | Neurogenic locus notch homolog protein 1 |
| NOXA | PMAIP1 | Phorbol-12-myristate-13-acetate-induced protein 1 |
| NRF2 | NFE2L2 | Nuclear factor erythroid 2-related factor 2 |
| p15 | CDKN2B | Cyclin-dependent kinase inhibitor 2B |
| p16 | CDKN2A | Cyclin-dependent kinase inhibitor 2A |
| p21 | CDKN1A | Cyclin-dependent kinase inhibitor 1A |
| p27 | CDKN1B | Cyclin-dependent kinase inhibitor 1B |
| p27-amp | CDKN1B | Cyclin-dependent kinase inhibitor 1B amplification |
| p53 | TP53 | Tumor protein p53 |
| p70S6K | RPS6KB1/RPS6KB2 | Ribosomal protein S6 kinase beta-1/2 |
| p90RSK | RPS6KA1/RPS6KA3 | Ribosomal protein S6 kinase alpha-1/3 |
| PALB2 | PALB2 | Partner and localizer of BRCA2 |
| PI3K | PIK3CA/PIK3CB/PIK3CD/PIK3C | Phosphoinositide 3-kinase catalytic subunits |
| PIK3CA | PIK3CA | Phosphoinositide 3-kinase catalytic subunit alpha |
| PKA | PRKACA/PRKACB/PRKACG | Protein kinase A catalytic subunits |
| Plk1 | PLK1 | Polo-like kinase 1 |
| pRB | RB1 | Retinoblastoma protein |
| PTEN | PTEN | Phosphatase and tensin homolog |

| Model node name | Gene symbol | Full protein name |
|---|---|---|
| PTPRK | PTPRK | Protein tyrosine phosphatase receptor type K |
| PTPRT | PTPRT | Protein tyrosine phosphatase receptor type T |
| PUMA | BBC3 | Bcl-2-binding component 3 |
| RAD50 | RAD50 | RAD50 homolog |
| RAD51 | RAD51 | RAD51 recombinase |
| RAF | RAF1 | Raf-1 proto-oncogene |
| RAP80 | UIMC1 | Ubiquitin interaction motif containing 1 |
| RAS | HRAS/KRAS/NRAS | Harvey/Kirsten/Neuroblastoma RAS viral oncogene homolog |
| RASSF1 | RASSF1 | Ras association domain family member 1 |
| RBX1 | RBX1 | Ring-box 1 |
| RNF168 | RNF168 | Ring finger protein 168 |
| RNF8 | RNF8 | Ring finger protein 8 |
| seDSB | | Single ended double stanrd break |
| SFRP4 | SFRP4 | Secreted frizzled-related protein 4 |
| SKP2 | SKP2 | S-phase kinase-associated protein 2 |
| SOS | SOS1/SOS2 | Son of sevenless homolog 1/2 |
| SSB | SSBP1 | Single-stranded DNA-binding protein 1 |
| STAT3 | STAT3 | Signal transducer and activator of transcription 3 |
| STIL | STIL | SCL/TAL1 interrupting locus |
| STK11 | STK11 | Serine/threonine kinase 11 |
| TCF | TCF7/TCF7L1/TCF7L2 | T-cell factor 7/7-like 1/7-like 2 |
| toxic-NHEJ | | Toxic non-homologous end joining |
| TSC | TSC1/TSC2 | Tuberous sclerosis complex 1/2 |
| Twist | TWIST1 | Twist family bHLH transcription factor 1 |
| UrseDSB | | Unresolved double-stranded break |
| Wee1 | WEE1 | WEE1 G2 checkpoint kinase |
| WNT | WNT1-WNT10B | Wingless-type MMTV integration site family |
| XLF | NHEJ1 | Non-homologous end joining factor 1 |
| XRCC4 | XRCC4 | X-ray repair cross-complementing protein 4 |
| YB1 | YBX1 | Y-box binding protein 1 |
| ZNRF3 | ZNRF3 | Zinc and ring finger 3 |

**Table S3. Summary of the nodes included in the model.** Summary of all the nodes, with corresponding gene symbols of the proteins involved where applicable.

**Table S4. List of mutations used to model cell lines.**

| Cell line | Mutated Gene | Model Value | Mutation | Mutation Function |
|---|---|---|---|---|
| HEALTHY | Erbb1 | 2 | | |
| HCC44 | STIL | 3 | p.G487fs*7 | Ambiguous function |
| HCC44 | LZTR1 | 0 | CNA loss | Loss of Function |
| HCC44 | p53 | 0 | p.R175L | Gain of Function |
| HCC44 | RAS | 3 | p.G12C | Gain of Function |
| NCIH1373 | ATM_deficient | 1 | p.E2943* | Loss of Function |
| NCIH1373 | CDKN2A | 0 | p.C72* | Loss of Function |
| NCIH1373 | CUL3 | 0 | p.E375* | Loss of Function |
| NCIH1373 | RAS | 3 | p.G12C | Gain of Function |
| NCIH1373 | SFRP4 | 0 | CNA loss | Loss of Function |
| NCIH23 | PTPRT | 0 | p.G495V | Loss of Function |
| NCIH23 | LRP | 0 | p.R4084L | Loss of Function |
| NCIH23 | STK11 | 0 | p.W332* | Loss of Function |
| NCIH23 | p53 | 0 | p.M246I | Ambiguous function |
| NCIH23 | RAS | 3 | p.G12C | Gain of Function |
| SW1573 | PI3K | 3 | p.K111E | Gain of Function |
| SW1573 | RAS | 3 | p.G12C | Gain of Function |
| SW1573 | CDKN2A | 0 | CNA loss | Loss of Function |
| SW1573 | Beta-Catenin | 3 | p.S33F | Gain of Function |
| NCIH358 | ELF3 | 0 | p.D239fs*62 | Loss of Function |
| NCIH358 | PTPRK | 0 | CNA loss | Loss of Function |
| NCIH358 | RAS | 3 | p.G12C | Gain of Function |
| NCIH358 | p53 | 0 | Deletion | Loss of Function |
| NCIH358 | LRP | 0 | p.S878* | Loss of Function |
| NCIH1792 | CDKN2A | 0 | CNA loss | Loss of Function |
| NCIH1792 | RAS | 3 | p.G12C | Gain of Function |
| NCIH1792 | p53 | 0 | CAN loss | Loss of Function |
| NCIH1792 | Erbb3 | 0 | p.V104 R106delVVR | Loss of Function |
| A549 | RAS | 3 | p.G12S | Gain of Function |
| A549 | STK11 | 0 | CNA loss | Loss of Function |
| A549 | KEAP1 | 0 | CNA loss | Loss of Function |
| A549 | CDKN2A | 0 | Deletion | Loss of Function |
| A549 | ATR | 0 | Splice Variant | Loss of Function |
| A549 | ARF | 0 | Deletion | Loss of Function |
| PC9 | Erbb1 | 3 | CNA gain | Gain of Function |
| PC9 | p53 | 0 | p.R248Q | Loss of Function |
| PC9 | CDKN2A | 0 | G67V | Loss of Function |
| HCC827 | Erbb1 | 3 | p.E746K | Gain of Function |
| HCC827 | p53 | 0 | p.V218L | Loss of Function |
| H1993 | p53 | 0 | p.C242W | Loss of Function |
| H1993 | STK11 | 0 | p.E199* | Loss of Function |
| H1993 | MET | 3 | Amplification | Gain of Function |

**Table S4. List of mutations used to model cell lines.** For each simulated cell line, the constrained nodes and their value are given, representing oncogenic mutations known to present in the cell line. A value of 0 indicates a loss-of-function mutation conferring no activity and for 1 basal activity, 2 elevated activity and 3 strong activation for gain-of-function mutations. Exact mutations and their functions were extracted from Cell Model Passport (Van der Meer, 2019).

**Table S5. Experiments from the literature used to validate the *in silico* model.**

| Source | Experiment | Cell line | Mutations | Constraints | Expected results | Model results |
|---|---|---|---|---|---|---|
| IR cell line sensitivity from Hannon lab | IR | NCIH1373 | CDKN2A OFF, PTPRK OFF, SFRP4 OFF, CUL3 OFF, p53 OFF, RAS Very high, ATM_deficient Low | IR_FLIP OFF | Death Low | Death Low |
| | No IR | NCIH1373 | CDKN2A OFF, PTPRK OFF, SFRP4 OFF, CUL3 OFF, p53 OFF, RAS Very high, ATM_deficient Low | | IR_FLIP Very high, Death Low | IR_FLIP Very high, Death Low |
| | IR | NCIH1792 | CDKN2A OFF, RAS Very high, p53 OFF, Erbb3 OFF | IR_FLIP OFF | Death Low | Death Low |
| | No IR | NCIH1792 | CDKN2A OFF, RAS Very high, p53 OFF, Erbb3 OFF | | IR_FLIP Very high, Death Low | IR_FLIP Very high, Death Low |
| | IR | NCIH358 | ELF3 OFF, LRP OFF, p53 OFF, PTPRK OFF, RAS Very high | IR_FLIP OFF | Death Low | Death Low |
| | No IR | NCIH358 | ELF3 OFF, LRP OFF, p53 OFF, PTPRK OFF, RAS Very high | | IR_FLIP Very high, Death Low | IR_FLIP Very high, Death Low |
| | IR | HCC44 | LZTR1 OFF, p53 OFF, RAS Very high, STIL Very high | IR_FLIP OFF | Death Low | Death Low |
| | No IR | HCC44 | LZTR1 OFF, p53 OFF, RAS Very high, STIL Very high | | IR_FLIP Very high, Death Low | IR_FLIP Very high, Death Low |
| | IR | NCIH2291 | MGA OFF, PTPRK OFF, p53 OFF, RAS Very high | IR_FLIP OFF | Death Low | Death Low |
| | No IR | NCIH2291 | MGA OFF, PTPRK OFF, p53 OFF, RAS Very high | | IR_FLIP Very high, Death Low | IR_FLIP Very high, Death Low |
| | IR | SW1573 | PI3K Very high, RAS Very high, Beta-Catenin Very high, CDKN2A OFF | IR_FLIP OFF | Death Low | Death Low |
| | No IR | SW1573 | PI3K Very high, RAS Very high, Beta-Catenin Very high, CDKN2A OFF | | IR_FLIP Very high, Death Low | IR_FLIP Very high, Death Low |
| | No IR | NCIH23 | PTPRT OFF, ATM_deficient Low, LRP OFF, STK11 OFF, p53 OFF, RAS Very high | | IR_FLIP Very high, Death Low | IR_FLIP Very high, Death Low |
| | IR | NCIH23 | PTPRT OFF, LRP OFF, STK11 OFF, p53 OFF, RAS Very high, ATM_deficient Low | IR_FLIP OFF | Death Very high | Death Low |
| | IR | NCIH2122 | STK11 OFF, RAS Very high, p53 OFF, KEAP1 OFF | IR_FLIP OFF | Death Very high | Death Low |
| | No IR | NCIH2122 | STK11 OFF, RAS Very high, p53 OFF, KEAP1 OFF | | IR_FLIP Very high, Death Low | IR_FLIP Very high, Death Low |
| Minjgee et al. 2011 | Control, no KRAS(V12) | FaDu | CDKN2A OFF, p53 OFF | | RAS High, Akt Low | RAS High, Akt Low |
| | Test, KRAS(V12) | FaDu | CDKN2A OFF, p53 OFF | RAS Very high | Akt High | Akt High |
| Zhang et al. 2007 | Control, scrambled siRNA | MCF-7 | NBS1 OFF, PIK3CA Very high | | PTPRT Low, STAT3 Low | PTPRT Low, STAT3 High |
| | Test, PTPRT siRNA | MCF-7 | NBS1 OFF, PIK3CA Very high | PTPRT OFF | STAT3 Very high | STAT3 Very high |
| Wael et al. 2014 | RNAi negative control | NCI-H2170 | RAS High, KEAP1 OFF, CDKN2A OFF, p53 OFF | | Proliferation High, Apoptosis Low, Akt Low | Proliferation High, Apoptosis OFF, Akt Low |
| | Test, NOTCH1 shRNA | NCI-H2170 | RAS High, KEAP1 OFF, CDKN2A OFF, p53 OFF | NOTCH1 OFF | Proliferation High, Apoptosis Low, Akt Low | Proliferation High, Apoptosis OFF, Akt Low |
| | RNAi negative control | A549 | RAS Very high, STK11 OFF, KEAP1 OFF, CDKN2A OFF, ATR OFF, ARF OFF | | Proliferation High, Apoptosis Low, Akt Low | Proliferation High, Apoptosis Low, Akt Low |
| | Test, NOTCH1 shRNA | A549 | RAS Very high, STK11 OFF, KEAP1 OFF, CDKN2A OFF, ATR OFF, ARF OFF | NOTCH1 OFF | Proliferation Very high, Apoptosis OFF, Akt High | Proliferation Very high, Apoptosis OFF, Akt High |

| | | | | | | |
|---|---|---|---|---|---|---|
| Cortez et al. 2015 | Control: no miR-34a, no ionising radiation | A549 | RAS Very high, STK11 OFF, KEAP1 OFF, CDKN2A OFF, ATR OFF, ARF OFF | | RAD51 Low, IR_FLIP Very high, Death OFF | RAD51 Low, IR_FLIP Very high, Death Low |
| | Ionising radiation, no miR-34a | A549 | RAS Very high, STK11 OFF, KEAP1 OFF, CDKN2A OFF, ATR OFF, ARF OFF | IR_FLIP OFF | RAD51 Very high, Death Low | RAD51 Very high, Death High |
| | miR-34a, 4Gy ionising radiation | A549 | RAS Very high, STK11 OFF, KEAP1 OFF, CDKN2A OFF, ATR OFF, ARF OFF | RAD51 OFF, IR_FLIP OFF | Death High | Death Very high |
| | miR-34a, no ionising radiation | A549 | RAS Very high, STK11 OFF, KEAP1 OFF, CDKN2A OFF, ATR OFF, ARF OFF | RAD51 OFF | IR_FLIP Very high, Death Low | IR_FLIP Very high, Death High |
| Llabata et al. 2020 | Control, empty ZsGreen | A549 | RAS Very high, STK11 OFF, KEAP1 OFF, CDKN2A OFF, ATR OFF, ARF OFF | | MGA Low, Proliferation High, MYCMAX High | MGA Low, Proliferation High, MYCMAX Very high |
| | Treat, MGA-ZsGreen overexpression vector | A549 | RAS Very high, STK11 OFF, KEAP1 OFF, CDKN2A OFF, ATR OFF, ARF OFF | MGA Very high | MYCMAX OFF, Proliferation OFF | MYCMAX OFF, Proliferation Low |
| Xu et al. 2017 | Control, control-siRNA | NCI-H1650 | APC OFF | | Akt High, ERK High, Apoptosis Low, Proliferation Low | Akt Low, ERK High, Apoptosis Low, Proliferation Low |
| | Treat, MUC1-siRNA | NCI-H1650 | APC OFF | MUC1 OFF | Akt Low, ERK Low, Apoptosis High, Proliferation OFF | Akt OFF, ERK OFF, Apoptosis High, Proliferation OFF |
| | Control, no JAK2 RNAi | A549 | RAS Very high, STK11 OFF, KEAP1 OFF, CDKN2A OFF, ATR OFF, ARF OFF | | Proliferation High, JAK2 High | Proliferation High, JAK2 High |
| | Treat, JAK2 RNAi | A549 | RAS Very high, STK11 OFF, KEAP1 OFF, CDKN2A OFF, ATR OFF, ARF OFF | JAK2 OFF | Proliferation Low | Proliferation Low |
| | Treat, flag-JAK2 | A549 | RAS Very high, STK11 OFF, KEAP1 OFF, CDKN2A OFF, ATR OFF, ARF OFF | JAK2 Very high | Proliferation Very high | Proliferation Very high |
| Yang et al. 2018 | Control, no DNA-PKcs inhibitor, no IR | A549 | RAS Very high, STK11 OFF, KEAP1 OFF, CDKN2A OFF, ATR OFF, ARF OFF | | IR_FLIP Very high, DSB Low, Death OFF | IR_FLIP Very high, DSB Low, Death Low |
| | DNA-PKcs inhibition, IR | A549 | RAS Very high, STK11 OFF, KEAP1 OFF, CDKN2A OFF, ATR OFF, ARF OFF | IR_FLIP OFF | DSB Very high, Death Very high | DSB Very high, Death Very high |
| | DNA-PKcs inhibition, no IR | A549 | RAS Very high, STK11 OFF, KEAP1 OFF, CDKN2A OFF, ATR OFF, ARF OFF | | IR_FLIP Very high, DSB Low, Death OFF | IR_FLIP Very high, DSB Low, Death Low |
| | IR, no DNA-PKcs inhibition | A549 | RAS Very high, STK11 OFF, KEAP1 OFF, CDKN2A OFF, ATR OFF, ARF OFF | IR_FLIP OFF | DSB Very high, Death High | DSB Very high, Death High |
| Wang et al. 2018 | Control, no shELF3 | A549 | RAS Very high, STK11 OFF, KEAP1 OFF, CDKN2A OFF, ATR OFF, ARF OFF | | ELF3 Low, PI3K Low, MEK Very high | ELF3 Low, PI3K Low, MEK Very high |
| | Test, shELF3 | A549 | RAS Very high, STK11 OFF, KEAP1 OFF, CDKN2A OFF, ATR OFF, ARF OFF | ELF3 OFF | PI3K Low, MEK Low | PI3K Low, MEK High |
| Edelmann et al. 2005 | Cells irradiated with 10 Gy after 4 days | BEAS-2B | EGF High | IR_FLIP OFF | Death High | Death High |
| | Control | BEAS-2B | EGF High | IR_FLIP Very high | Death Low | Death Low |
| Sato et al. 2020 | Control | HBEC3 | EGF High | | Proliferation Low, p21 High | Proliferation Low, p21 Low |
| | KRASV12 transfected | HBEC3 | EGF High | RAS Very high | Proliferation High, p21 High | Proliferation High, p21 High |
| | p53RNAi | HBEC3 | EGF High | p53 OFF | Proliferation High, p21 OFF | Proliferation Low, p21 OFF |
| | p53RNAi and KRASV12 transfected | HBEC3 | EGF High | RAS Very high, p53 OFF | Proliferation Very high, p21 OFF | Proliferation Very high, p21 OFF |

| | | | | | | |
|---|---|---|---|---|---|---|
| Fok et al. 2019 | Control | A549 | RAS Very high, STK11 OFF, KEAP1 OFF, CDKN2A OFF, ATR OFF, ARF OFF | | IR_FLIP Very high, Death OFF | IR_FLIP Very high, Death Low |
| | Test, AZD7648 | A549 | RAS Very high, STK11 OFF, KEAP1 OFF, CDKN2A OFF, ATR OFF, ARF OFF | | IR_FLIP Very high, Death Low | IR_FLIP Very high, Death Low |
| | Test, AZD7648 plus IR 4Gy | A549 | RAS Very high, STK11 OFF, KEAP1 OFF, CDKN2A OFF, ATR OFF, ARF OFF | IR_FLIP OFF | Death Very high | Death Very high |
| | Test, IR 4Gy | A549 | RAS Very high, STK11 OFF, KEAP1 OFF, CDKN2A OFF, ATR OFF, ARF OFF | IR_FLIP OFF | Death Low | Death High |
| Zhao et al. 2016 | Control, no Gefitinib | A549 | RAS Very high, STK11 OFF, KEAP1 OFF, CDKN2A OFF, ATR OFF, ARF OFF | | Erbb1 High, Apoptosis OFF, Proliferation High, PI3K Low, Akt Low, mTORC1 High, mTORC2 Low | Erbb1 High, Apoptosis Low, Proliferation High, PI3K Low, Akt Low, mTORC1 High, mTORC2 Low |
| | Treat, Gefitinib | A549 | RAS Very high, STK11 OFF, KEAP1 OFF, CDKN2A OFF, ATR OFF, ARF OFF | Erbb1 OFF | Apoptosis High, Proliferation OFF, PI3K OFF, Akt OFF, mTORC1 Low, mTORC2 Low | Apoptosis Low, Proliferation Low, PI3K OFF, Akt OFF, mTORC1 High, mTORC2 OFF |
| | Treat, Gefitinib and LY294002 | A549 | RAS Very high, STK11 OFF, KEAP1 OFF, CDKN2A OFF, ATR OFF, ARF OFF | Erbb1 OFF, PI3K OFF | Apoptosis High | Apoptosis High |
| | Treat, LY294002 | A549 | RAS Very high, STK11 OFF, KEAP1 OFF, CDKN2A OFF, ATR OFF, ARF OFF | Erbb1 OFF, PI3K OFF | Apoptosis High | Apoptosis High |
| Gao et al. 2017 | Control, no COMPOUND7809 | A549 | RAS Very high, STK11 OFF, KEAP1 OFF, CDKN2A OFF, ATR OFF, ARF OFF | | EGFR High, Proliferation High, Apoptosis OFF | EGFR High, Proliferation High, Apoptosis Low |
| | Treat, COMPOUND7809 | A549 | RAS Very high, STK11 OFF, KEAP1 OFF, CDKN2A OFF, ATR OFF, ARF OFF | EGFR OFF | Proliferation OFF, Apoptosis Low | Proliferation Low, Apoptosis Low |
| | Control, no COMPOUND7809 | SK-LU-1 | p53 OFF, RAS Very high, APC OFF | | EGFR High, Proliferation High, Apoptosis OFF | EGFR High, Proliferation Very high, Apoptosis OFF |
| | Treat, COMPOUND7809 | SK-LU-1 | p53 OFF, RAS Very high, APC OFF | EGFR OFF | Proliferation Low, Apoptosis Low | Proliferation High, Apoptosis OFF |
| | Control, no COMPOUND7809 | NCIH23 | p53 OFF, RAS Very high, STK11 OFF, LRP OFF, ATM_deficient Low | | EGFR High, Proliferation High, Apoptosis OFF | EGFR Low, Proliferation High, Apoptosis Low |
| | Treat, COMPOUND7809 | NCIH23 | p53 OFF, RAS Very high, STK11 OFF, LRP OFF, ATM_deficient Low | EGFR OFF | Proliferation Low, Apoptosis Low | Proliferation Low, Apoptosis Low |
| Bigenzahn et al. 2018 | Control, no LZTR1 sgRNA | K-562 | p53 OFF, Akt Very high | | LZTR1 Low, MEK High | LZTR1 Low, MEK High |
| | Test, LZTR1 sgRNA | K-562 | p53 OFF, Akt Very high | LZTR1 OFF | MEK Very high | MEK High |
| Wang et al. 2017 | 4Gy IR, no KRASmut transfection | NCI-H1703 | CDKN2A OFF, p53 OFF | IR_FLIP OFF | RAS High, Death High | RAS High, Death High |
| | Control, no IR or KRASmut transfection | NCI-H1703 | CDKN2A OFF, p53 OFF | | RAS High, IR_FLIP Very high, Death Low | RAS High, IR_FLIP Very high, Death Low |
| | KRASmut transfection, no IR | NCI-H1703 | CDKN2A OFF, p53 OFF | RAS Very high | IR_FLIP Very high, Death Low | IR_FLIP Very high, Death Low |
| | KRASmut transfection, 4Gy IR | NCI-H1703 | CDKN2A OFF, p53 OFF | RAS Very high, IR_FLIP OFF | Death Low | Death Low |
| Zhou et al. 2013 | Control | A549 | RAS Very high, STK11 OFF, KEAP1 OFF, CDKN2A OFF, ATR OFF, ARF OFF | | IR_FLIP Very high, RNF8 Low, Death OFF, RAD51 Low, G2_M_arrest Low | IR_FLIP Very high, RNF8 Low, Death Low, RAD51 Low, G2_M_arrest Low |
| | IR (2Gy) | A549 | RAS Very high, STK11 OFF, KEAP1 OFF, CDKN2A OFF, ATR OFF, ARF OFF | IR_FLIP OFF | RNF8 Very high, Death High, RAD51 Very high, G2_M_arrest High | RNF8 Very high, Death High, RAD51 Very high, G2_M_arrest Low |
| | IR (2Gy), Lenti-siRNF8 | A549 | RAS Very high, STK11 OFF, KEAP1 OFF, CDKN2A OFF, ATR OFF, ARF OFF | IR_FLIP OFF, RNF8 OFF | Death Very high, RAD51 Low, G2_M_arrest High | Death Very high, RAD51 OFF, G2_M_arrest Low |

| | | | | | | |
|---|---|---|---|---|---|---|
| | Lenti-siRNF8 | A549 | RAS Very high, STK11 OFF, KEAP1 OFF, CDKN2A OFF, ATR OFF, ARF OFF | RNF8 OFF | IR_FLIP Very high, Death OFF, RAD51 Low, G2_M_arrest High | IR_FLIP Very high, Death Low, RAD51 OFF, G2_M_arrest Low |
| Francesco et al. 2013 | 2Gy ionising radiation, no miR-27a | A549 | RAS Very high, STK11 OFF, KEAP1 OFF, CDKN2A OFF, ATR OFF, ARF OFF | IR_FLIP OFF | ATM Very high, G2_M_arrest Low, Proliferation High, Death Very high | ATM Very high, G2_M_arrest Low, Proliferation High, Death High |
| | Control, no miR-27a, no ionising radiation | A549 | RAS Very high, STK11 OFF, KEAP1 OFF, CDKN2A OFF, ATR OFF, ARF OFF | | ATM Low, IR_FLIP Very high, Proliferation High, Death OFF | ATM Low, IR_FLIP Very high, Proliferation High, Death Low |
| | miR27a, 2Gy ionising radiation | A549 | RAS Very high, STK11 OFF, KEAP1 OFF, CDKN2A OFF, ATR OFF, ARF OFF | ATM OFF, IR_FLIP OFF | G2_M_arrest OFF, Proliferation Very high, Death High | G2_M_arrest OFF, Proliferation Very high, Death Very high |
| | miR27a, no 2Gy ionising radiation | A549 | RAS Very high, STK11 OFF, KEAP1 OFF, CDKN2A OFF, ATR OFF, ARF OFF | ATM OFF | IR_FLIP Very high, Proliferation Very high, Death OFF | IR_FLIP Very high, Proliferation Very high, Death High |
| You et al. 2004 | Control, no anti-Wnt2 monoclonal antibody | A549 | RAS Very high, STK11 OFF, KEAP1 OFF, CDKN2A OFF, ATR OFF, ARF OFF | | WNT High, Caspase3 OFF, Apoptosis Low | WNT High, Caspase3 OFF, Apoptosis Low |
| | Test, treatment with anti-Wnt2 monoclonal antibody | A549 | RAS Very high, STK11 OFF, KEAP1 OFF, CDKN2A OFF, ATR OFF, ARF OFF | WNT OFF | Caspase3 High, Apoptosis High | Caspase3 OFF, Apoptosis Low |
| Xu et al. 2019 | Control, no PTPRK siRNA | A549 | RAS Very high, STK11 OFF, KEAP1 OFF, CDKN2A OFF, ATR OFF, ARF OFF | | PTPRK High, Proliferation High | PTPRK High, Proliferation High |
| | Test, PTPRK siRNA | A549 | RAS Very high, STK11 OFF, KEAP1 OFF, CDKN2A OFF, ATR OFF, ARF OFF | PTPRK OFF | Proliferation Very high | Proliferation Very high |
| Dorr et al. 2015 | Control, no CUL3 siRNA | A549 | RAS Very high, STK11 OFF, CDKN2A OFF, ATR OFF, ARF OFF | | CUL3 High, NRF2 High, Proliferation High | CUL3 High, NRF2 Very high, Proliferation High |
| | Test, CUL3 siRNA | A549 | RAS Very high, STK11 OFF, CDKN2A OFF, ATR OFF, ARF OFF | CUL3 OFF | NRF2 Very high, Proliferation Very high | NRF2 Very high, Proliferation High |
| Homma et al. 2009 | Control, no NRF2 siRNA | NCI-H292 | ARF OFF, Erbb1 Very high | | NRF2 High, Proliferation High, G2_M_arrest OFF, G1arrest OFF, pRB OFF, p21 OFF | NRF2 Very high, Proliferation High, G2_M_arrest OFF, G1arrest OFF, pRB OFF, p21 High |
| | Treat, NRF2 siRNA | NCI-H292 | ARF OFF, Erbb1 Very high | NRF2 OFF | Proliferation Low, G2_M_arrest Low, G1arrest Low, pRB Low, p21 Low | Proliferation Low, G2_M_arrest Low, G1arrest Low, pRB Low, p21 Low |
| | Control, no NRF2 siRNA | A549 | RAS Very high, STK11 OFF, KEAP1 OFF, CDKN2A OFF, ATR OFF, ARF OFF | | NRF2 High, Proliferation High, G2_M_arrest OFF, G1arrest OFF, pRB OFF, p21 Low | NRF2 Very high, Proliferation High, G2_M_arrest Low, G1arrest OFF, pRB OFF, p21 High |
| | Treat, NRF2 siRNA | A549 | RAS Very high, STK11 OFF, KEAP1 OFF, CDKN2A OFF, ATR OFF, ARF OFF | NRF2 OFF | Proliferation Low, G2_M_arrest High, G1arrest Low, pRB Low, p21 High | Proliferation Low, G2_M_arrest High, G1arrest Low, pRB Low, p21 Low |
| | Control, no trametinib | PC-9 | Erbb1 Very high, CDKN2A OFF, p53 OFF | | MEK High, Akt High, MET High, ERK High, Caspase3 OFF, Apoptosis OFF, Proliferation High | MEK Very high, Akt High, MET High, ERK Very high, Caspase3 OFF, Apoptosis OFF, Proliferation Very high |
| | Treat, LY294002 | PC-9 | Erbb1 Very high, CDKN2A OFF, p53 OFF | PI3K OFF | Caspase3 OFF, Proliferation High | Caspase3 OFF, Proliferation High |
| | Treat, trametinib | PC-9 | Erbb1 Very high, CDKN2A OFF, p53 OFF | MEK OFF | Akt Very high, MET Low, ERK OFF, Caspase3 OFF, Apoptosis OFF, Proliferation High | Akt Very high, MET High, ERK OFF, Caspase3 OFF, Apoptosis OFF, Proliferation Low |

| | | | | | | |
|---|---|---|---|---|---|---|
| Chiba et al. 2016 | Treat, trametinib, LY294002 | PC-9 | Erbb1 Very high, CDKN2A OFF, p53 OFF | MEK OFF, PI3K OFF | MET High, ERK OFF, Proliferation OFF, Caspase3 High | MET High, ERK OFF, Proliferation OFF, Caspase3 High |
| | Control, no trametinib | HCC827 | Erbb1 Very high, p53 OFF | | MEK High, Akt High, MET High, ERK High, Caspase3 OFF, Apoptosis OFF, Proliferation High | MEK Very high, Akt High, MET High, ERK Very high, Caspase3 OFF, Apoptosis OFF, Proliferation Very high |
| | Treat, LY294002 | HCC827 | Erbb1 Very high, p53 OFF | PI3K OFF | Caspase3 OFF, Proliferation High | Caspase3 OFF, Proliferation High |
| | Treat, trametinib | HCC827 | Erbb1 Very high, p53 OFF | MEK OFF | Akt Very high, MET High, ERK OFF, Caspase3 OFF, Apoptosis OFF, Proliferation High | Akt Very high, MET High, ERK OFF, Caspase3 OFF, Apoptosis OFF, Proliferation Low |
| | Treat, trametinib, LY294002 | HCC827 | Erbb1 Very high, p53 OFF | MEK OFF, PI3K OFF | MET High, ERK OFF, Caspase3 High, Proliferation OFF | MET High, ERK OFF, Caspase3 High, Proliferation OFF |
| | Control, no trametinib | HCC827CNXR | p53 OFF | MET Very high | MEK High, Akt High, ERK High, Caspase3 OFF, Apoptosis OFF, Proliferation High | MEK Very high, Akt High, ERK Very high, Caspase3 OFF, Apoptosis OFF, Proliferation Very high |
| | Treat, trametinib | HCC827CNXR | p53 OFF | MET Very high, MEK OFF | Akt High, ERK OFF, Caspase3 Low, Apoptosis Low, Proliferation OFF | Akt Low, ERK OFF, Caspase3 Low, Apoptosis Low, Proliferation OFF |
| | Control, no trametinib | EBC-1 | p53 OFF, NRF2 Very high | MET Very high | MEK High, Erbb1 High, Akt High, ERK High, Caspase3 OFF, Apoptosis OFF, Proliferation Very high | MEK Very high, Erbb1 High, Akt High, ERK Very high, Caspase3 OFF, Apoptosis OFF, Proliferation Very high |
| | Treat, crizotinib | EBC-1 | p53 OFF, NRF2 Very high | MET OFF | Caspase3 Low, Apoptosis Low, Proliferation Low | Caspase3 Low, Apoptosis Low, Proliferation Low |
| | Treat, trametinib | EBC-1 | p53 OFF, NRF2 Very high | MET Very high, MEK OFF | Erbb1 Low, Akt High, ERK OFF, Caspase3 High, Apoptosis Low, Proliferation Low | Erbb1 High, Akt Low, ERK OFF, Caspase3 Low, Apoptosis Low, Proliferation OFF |
| | Treat, trametinib, crizotinib | EBC-1 | p53 OFF, NRF2 Very high | MET OFF, MEK OFF | ERK OFF, Caspase3 High, Apoptosis High, Proliferation Low | ERK OFF, Caspase3 High, Apoptosis High, Proliferation OFF |
| | Control, no trametinib | H1993 | p53 OFF, STK11 OFF | MET Very high | MEK High, Erbb1 High, Akt High, ERK High, Caspase3 OFF, Apoptosis OFF, Proliferation High | MEK Very high, Erbb1 High, Akt High, ERK Very high, Caspase3 OFF, Apoptosis OFF, Proliferation Very high |
| | Treat, crizotinib | H1993 | p53 OFF, STK11 OFF | MET OFF | ERK Low, Caspase3 Low, Apoptosis Low, Proliferation Low | ERK Low, Caspase3 Low, Apoptosis Low, Proliferation Low |
| | Treat, trametinib | H1993 | p53 OFF, STK11 OFF | MET Very high, MEK OFF | Erbb1 Low, Akt High, ERK OFF, Caspase3 High, Apoptosis High, Proliferation OFF | Erbb1 High, Akt High, ERK OFF, Caspase3 Low, Apoptosis Low, Proliferation Low |
| | Treat, trametinib, crizotinib | H1993 | p53 OFF, STK11 OFF | MEK OFF, MET OFF | ERK OFF, Caspase3 High, Apoptosis High, Proliferation OFF | ERK OFF, Caspase3 High, Apoptosis High, Proliferation OFF |

| | | | | | | |
|---|---|---|---|---|---|---|
| Li et al. 2015 | Treat, WB-308 | NCI-H1975 | Erbb1 OFF, APC OFF, ATR OFF, p53 OFF, TSC OFF | | Akt OFF, ERK OFF, p21 Low, BCL-2 OFF, BAKBAX High, CycD OFF, Apoptosis High, Proliferation OFF | Akt OFF, ERK OFF, p21 High, BCL-2 OFF, BAKBAX High, CycD OFF, Apoptosis High, Proliferation OFF |
| | Control, no WB-308 | NCI-H1975 | Erbb1 Very high, APC OFF, ATR OFF, p53 OFF, TSC OFF | | Akt High, ERK Very high, p21 High, BCL-2 High, BAKBAX Low, CycD High, Apoptosis OFF, Proliferation Very high | Akt High, ERK Very high, p21 OFF, BCL-2 Very high, BAKBAX OFF, CycD Very high, Apoptosis OFF, Proliferation Very high |
| | Control, no WB-308 | PC-9 | Erbb1 Very high, p53 OFF, CDKN2A OFF | | Akt High, ERK Very high, p21 High, BCL-2 Very high, BAKBAX Low, CycD High, Caspase3 OFF, G2_M_arrest Low, Apoptosis OFF, Proliferation Very high | Akt High, ERK Very high, p21 OFF, BCL-2 Very high, BAKBAX OFF, CycD Very high, Caspase3 OFF, G2_M_arrest OFF, Apoptosis OFF, Proliferation Very high |
| | Control, no WB-308 | A549 | RAS Very high, STK11 OFF, KEAP1 OFF, CDKN2A OFF, ATR OFF, ARF OFF | | EGFR High, Akt High, ERK Very high, p21 High, BCL-2 High, BAKBAX OFF, CycD High, Apoptosis OFF, Proliferation High | EGFR High, Akt Low, ERK Very high, p21 High, BCL-2 High, BAKBAX Low, CycD Very high, Apoptosis Low, Proliferation High |
| | Treat, WB-308 | A549 | RAS Very high, STK11 OFF, KEAP1 OFF, CDKN2A OFF, ATR OFF, ARF OFF | EGFR OFF | Akt Low, ERK High, p21 Low, BCL-2 Low, BAKBAX Low, CycD Low, Apoptosis Low, Proliferation OFF | Akt OFF, ERK Very high, p21 High, BCL-2 Low, BAKBAX Low, CycD Very high, Apoptosis Low, Proliferation Low |
| | Treat, WB-308 | PC-9 | p53 OFF, CDKN2A OFF | Erbb1 OFF | Akt OFF, ERK OFF, p21 Low, BCL-2 OFF, BAKBAX High, CycD OFF, G2_M_arrest High, Caspase3 High, Apoptosis High, Proliferation OFF | Akt OFF, ERK OFF, p21 Low, BCL-2 OFF, BAKBAX High, CycD OFF, G2_M_arrest Very high, Caspase3 High, Apoptosis High, Proliferation OFF |
| Wang et al. 2008 | Control | A549 | RAS Very high, STK11 OFF, KEAP1 OFF, CDKN2A OFF, ATR OFF, ARF OFF | | Death Low | Death Low |
| | Nrf2 siRNA + doxorubicin | A549 | RAS Very high, STK11 OFF, KEAP1 OFF, CDKN2A OFF, ATR OFF, ARF OFF | IR_FLIP OFF, NRF2 OFF | Death Very high | Death Very high |
| | Transient knockdown of Nrf2 | A549 | RAS Very high, STK11 OFF, KEAP1 OFF, CDKN2A OFF, ATR OFF, ARF OFF | NRF2 OFF | Death High | Death Low |

**Table S5. Experiments from the literature used to validate the *in silico* model.** For each experiment, the table shows the cell line used, the constraints applied to the model and the expected outcome, based on the experimental result. The final column shows the result of the model simulations.

**Table S6. List of druggable nodes in the model and the drugs that target them.**

| Node | GeneSymbol | UNIPROT ID | FDA approval | chEMBL ID |
|---|---|---|---|---|
| 4EBP1 | EIF4EBP1 | Q13541 | FALSE | CHEMBL3351214, CHEMBL4296088 |
| Akt | AKT1 | P31749 | FALSE | CHEMBL4523748, CHEMBL3038463, CHEMBL3885629, CHEMBL4106175, CHEMBL5169081, CHEMBL5169068, CHEMBL2111353, CHEMBL4282 |
| AMPK | PRKAA2 | P54646 | TRUE | CHEMBL3038455, CHEMBL3885504, CHEMBL3038456, CHEMBL4106159, CHEMBL4106158, CHEMBL3038457, CHEMBL3038454, CHEMBL2116, CHEMBL2096907 |
| APC | APC | D6RFL6 | FALSE | |
| ARF | CDKN2A | P42771 | FALSE | CHEMBL4680027 |
| ATR | ATR | H0Y8Y6 | FALSE | |
| BAD | BAD | Q92934 | FALSE | CHEMBL5169266, CHEMBL3817 |
| BAKBAX | BAK1 | Q16611 | FALSE | CHEMBL3885523, CHEMBL3885516, CHEMBL3137283, CHEMBL5609, CHEMBL5169269 |
| BAKBAX | BAX | Q07812 | FALSE | |
| Beta-Catenin | CTNNB1 | C9IZ65 | FALSE | |
| BCL-2 | BCL2 | P10415 | TRUE | CHEMBL4523685, CHEMBL4748224, CHEMBL5169264, CHEMBL5169266, CHEMBL3885516, CHEMBL3885513, CHEMBL4860, CHEMBL5169265 |
| BCL-W | BCL2L2 | G3V3B7 | FALSE | |
| BCL-XL | BCL2L1 | Q07817 | FALSE | CHEMBL3883285, CHEMBL3883286, CHEMBL3137283, CHEMBL4523705, CHEMBL3883287, CHEMBL4625, CHEMBL3883288, CHEMBL4748229 |
| Bim | BCL2L11 | C9J417 | FALSE | |
| CDC25A | CDC25A | P30304 | FALSE | CHEMBL3775 |
| CDK2 | CDK2 | P24941 | FALSE | CHEMBL4523633, CHEMBL2094128, CHEMBL3038470, CHEMBL3885552, CHEMBL2111326, CHEMBL4888454, CHEMBL4106153, CHEMBL4888444, CHEMBL4106152, CHEMBL3038517, CHEMBL4523689, CHEMBL301, CHEMBL1907605, CHEMBL2094126, CHEMBL4523634, CHEMBL3038469, CHEMBL3559691 |
| CDKN2A | CDKN2A | P42771 | FALSE | CHEMBL4680027 |
| CHK1 | CHEK1 | E9PRU7 | FALSE | |
| CHK2 | CHEK1 | E9PRU7 | FALSE | |
| c-Jun | JUN | P05412 | TRUE | CHEMBL4977, CHEMBL2111421 |
| CycE | CCNE1 | P24864 | FALSE | CHEMBL2111444, CHEMBL4523715, CHEMBL3885554, CHEMBL3038471, CHEMBL1907605, CHEMBL2094126, CHEMBL3617, CHEMBL3038468 |
| Dishevelled | DVL1 | O14640 | FALSE | CHEMBL6027 |
| E2F | E2F1 | Q01094 | FALSE | CHEMBL4382, CHEMBL4630726 |
| EGFR | EGFR | Q504U8 | FALSE | |
| FOXM1 | FOXM1 | Q08050 | FALSE | |
| FOXO3 | FOXO3 | O43524 | FALSE | CHEMBL5778 |
| GRB2 | GRB2 | J3QLF6 | FALSE | |
| GSK3B | GSK3B | P49841 | FALSE | CHEMBL3883309, CHEMBL262, CHEMBL2095188 |

| Node | GeneSymbol | UNIPROT ID | FDA approval | chEMBL ID |
|---|---|---|---|---|
| JAK2 | JAK2 | O60674 | FALSE | |
| KEAP1 | KEAP1 | Q14145 | TRUE | CHEMBL4296122, CHEMBL4106129, CHEMBL4296123, CHEMBL2069156, CHEMBL3038498 |
| LRP | LRP1 | Q7Z7K9 | FALSE | |
| MCL-1 | MCL1 | Q07820 | FALSE | CHEMBL4523712, CHEMBL3430887, CHEMBL3885523, CHEMBL3430886, CHEMBL3885515, CHEMBL4361 |
| MDM2 | MDM2 | Q00987 | FALSE | CHEMBL4523719, CHEMBL4879538, CHEMBL4523702, CHEMBL3883306, CHEMBL3038493, CHEMBL4523720, CHEMBL4523718, CHEMBL5023, CHEMBL4888446, CHEMBL4106123, CHEMBL1907611 |
| MEK | MAP2K1 | Q02750 | TRUE | CHEMBL4523703, CHEMBL3885566, CHEMBL4523755, CHEMBL4523740, CHEMBL3587, CHEMBL2111351, CHEMBL2111289 |
| MET | MET | C9JKM5 | FALSE | |
| mTORC1 | MTOR | B1AKP8 | FALSE | |
| mTORC2 | MTOR | B1AKP8 | FALSE | |
| NFkB | NFKB1 | P19838 | TRUE | CHEMBL2094258, CHEMBL3251 |
| NRF2 | NFE2L2 | H7C498 | FALSE | |
| p21 | CDKN1A | P38936 | FALSE | |
| p27 | CDKN1B | P46527 | FALSE | |
| p53 | TP53 | E7EQX7 | FALSE | |
| p70S6K | RPS6KB1 | K7ER06 | FALSE | |
| p90RSK | RPS6KA1 | E9PPC1 | FALSE | |
| Plk1 | PLK1 | P53350 | FALSE | |
| pRB | RB1 | P06400 | FALSE | |
| PTEN | PTEN | P60484 | FALSE | CHEMBL4523606, CHEMBL2052032 |
| PUMA | BBC3 | Q9BXH1 | FALSE | |
| RAF | RAF1 | P04049 | TRUE | CHEMBL4106189, CHEMBL3885566, CHEMBL2111351, CHEMBL3883317, CHEMBL3559685, CHEMBL1906 |
| RAF | BRAF | H7C560 | FALSE | |
| RAS | KRAS | P01116 | TRUE | CHEMBL5169273, CHEMBL4523623, CHEMBL2189121, CHEMBL4524006 |
| STAT3 | STAT3 | P40763 | FALSE | CHEMBL4296101, CHEMBL4026, CHEMBL4523691 |
| SOS | SOS1 | Q07889 | FALSE | CHEMBL2079846, CHEMBL5169070 |
| STK11 | STK11 | A0A087WT72 | FALSE | |
| Wee1 | WEE1 | P30291 | FALSE | CHEMBL5169067, CHEMBL4630732, CHEMBL5491 |
| YB1 | YBX1 | P67809 | FALSE | CHEMBL4296006 |
| Caspase3 | CASP3 | A8MVM1 | FALSE | |
| CDK1 | CDK1 | E5RIU6 | FALSE | |
| CycB | CCNB1 | P14635 | FALSE | CHEMBL2412, CHEMBL4523630, CHEMBL4680053, CHEMBL1907602, CHEMBL2094127 |
| CycD | CCND1 | P24385 | TRUE | CHEMBL2095942, CHEMBL3885552, CHEMBL1907601, CHEMBL3610, CHEMBL4523688, CHEMBL3885551, CHEMBL2111455 |

| Node | GeneSymbol | UNIPROT ID | FDA approval | chEMBL ID |
|---|---|---|---|---|
| CDK4 | CDK4 | P11802 | TRUE | CHEMBL3038472, CHEMBL2095942, CHEMBL3301385, CHEMBL2111326, CHEMBL4888444, CHEMBL4523963, CHEMBL4106184, CHEMBL4630748, CHEMBL3038517, CHEMBL4523715, CHEMBL3885554, CHEMBL4523686, CHEMBL4523732, CHEMBL1907601, CHEMBL3885553, CHEMBL3885548, CHEMBL3559691, CHEMBL331 |
| hTERT | TERT | O14746 | FALSE | |
| p16 | CDKN2A | P42771 | FALSE | CHEMBL4680027 |
| Erbb1 | EGFR | Q504U8 | FALSE | |
| Erbb3 | ERBB3 | F8VRL0 | FALSE | |
| CDC25C | CDC25C | P30307 | FALSE | CHEMBL2378 |
| ATM | ATM | E9PIN0 | FALSE | |
| Fyn | FYN | P06241 | FALSE | CHEMBL2363074, CHEMBL1841 |
| IL-6 | IL6 | P05231 | TRUE | CHEMBL1795129 |
| PKA | PRKACA | P17612 | FALSE | CHEMBL4101, CHEMBL2094138 |
| CUL3 | CUL3 | Q13618 | FALSE | CHEMBL4296124, CHEMBL4296123 |
| RBX1 | RBX1 | P62877 | FALSE | |
| MYC | MYC | A0A0B4J1R1 | FALSE | |
| MAX | MAX | P61244 | FALSE | CHEMBL1250363, CHEMBL4106127, CHEMBL3301395 |
| Ku | XRCC6 | P12956 | FALSE | CHEMBL4106136 |
| Ku | XRCC5 | P13010 | FALSE | CHEMBL4106136 |
| DNA-PKcs | PRKDC | P78527 | FALSE | CHEMBL4106136, CHEMBL3142 |
| MRE11 | MRE11 | P49959 | FALSE | |
| NBS1 | NBN | O60934 | FALSE | CHEMBL5169101 |
| 14_3_3 | YWHAZ | P63104 | FALSE | CHEMBL4105899 |
| RAD51 | RAD51 | E9PNT5 | FALSE | |
| 53BP1 | TP53BP1 | Q12888 | FALSE | CHEMBL2424509 |
| BRCA1 | BRCA1 | P38398 | FALSE | CHEMBL5990 |
| XRCC4 | ERCC4 | Q92889 | FALSE | CHEMBL3883316 |
| LIG4 | LIG4 | P49917 | FALSE | CHEMBL4296097, CHEMBL4523595 |
| KU70 | XRCC6 | P12956 | FALSE | CHEMBL4106136 |
| KU80 | XRCC5 | P13010 | FALSE | CHEMBL4106136 |
| RNF168 | RNF168 | Q8IYW5 | FALSE | CHEMBL5169175 |
| ELF3 | ELF3 | P78545 | FALSE | CHEMBL2111426 |
| SKP2 | SKP2 | Q13309 | FALSE | CHEMBL3885557, CHEMBL3885558, CHEMBL3885634 |
| Caspase9 | CASP9 | F8VVS7 | FALSE | |
| Frizzled | FZD1 | Q9UP38 | FALSE | CHEMBL2346493 |
| CBF | RBPJ | D6RAT2 | FALSE | |
| NOTCH1 | NOTCH1 | P46531 | FALSE | CHEMBL4524007, CHEMBL2146346 |

**Table S6. List of druggable nodes in the *in silico* model and the drugs that target them.** Each row is a node from the model, with the gene symbol, Uniprot ID and whether that protein as an FDA approved drug targeting it. Additionally, those drugs with chemical probes are shown using the chEMBL database.

**Table S7. List of drugs used in Nair et al (2022). and their corresponding model targets.**

| DrugID | Drug Name | Primary target |
|---|---|---|
| 1 | Erlotinib | EGFR |
| 14 | Neratinib | Erbb1 |
| 29 | AZ628 | RAF |
| 37 | Crizotinib | MET |
| 60 | BI-2536 | Plk1 |
| 75 | RAF265 | RAF |
| 94 | TGX 221 | PI3K |
| 96 | Pimozide | STAT3 |
| 97 | Selumetinib | MEK |
| 99 | Vandetanib | EGFR |
| 113 | Alpelisib | PI3K |
| 119 | Lapatinib | Erbb1 |
| 125 | Dactolisib | PI3K |
| 154 | CHIR-99021 | GSK3B |
| 156 | AZD6482 | PI3K |
| 182 | Obatoclax | BCL-2 |
| 210 | Vemurafenib | RAF |
| 224 | AS605240 | PI3K |
| 238 | Idelalisib | PI3K |
| 246 | Nutlin | MDM2 |
| 249 | Cabozantinib | MET |
| 283 | Omipalisib | PI3K |
| 306 | Fedratinib | JAK2 |
| 326 | GSK690693 | Akt |
| 357 | MK-2206 | Akt |
| 368 | Afatinib | Erbb1 |
| 373 | Pictilisib | PI3K |
| 385 | Navitoclax | BCL-2 |
| 398 | VE-821 | ATR |
| 412 | Venetoclax | BCL-2 |
| 421 | KU-55933 | ATM |
| 429 | Dabrafenib | RAF |
| 430 | Trametinib | MEK |
| 446 | MIM1 | MCL-1 |
| 461 | NSC319726 | p53 |
| 519 | Mirin | MRE11 |
| 522 | RO-3306 | CDK1 |
| 527 | AT9283 | JAK2 |
| 530 | Serdemetan | MDM2 |
| 537 | SCH900776 | CHK1 |
| 538 | KU-60019 | ATM |
| 564 | MK-8033 | MET |
| 589 | Tivantinib | MET |

**Table S7. List of drugs used in Nair et al. (2022) and their corresponding model targets.** Drug name is shown along with its target and ID used in Nair et al. (2022).

[2] Victor Llombart and Marc R. Mansour. Therapeutic targeting of “undruggable” MYC. *eBioMedicine*, 75, January 2022. Publisher: Elsevier.

[3] Marie-Eve Beaulieu, Toni Jauset, Daniel Massó-Vallés, Sandra Martínez-Martín, Peter Rahl, Loïka Maltais, Mariano F. Zacarias-Fluck, Sílvia Casacuberta-Serra, Erika Serrano del Pozo, Christopher Fiore, Laia Foradada, Virginia Castillo Cano, Meritxell Sánchez-Hervás, Matthew Guenther, Eduardo Romero Sanz, Marta Oteo, Cynthia Tremblay, Génesis Martín, Danny Letourneau, Martin Montagne, Miguel Ángel Morcillo Alonso, Jonathan R. Whitfield, Pierre Lavigne, and Laura Soucek. Intrinsic cell-penetrating activity propels Omomyc from proof of concept to viable anti-MYC therapy. *Science Translational Medicine*, 11(484):eaar5012, March 2019. Publisher: American Association for the Advancement of Science.

[4] Nishanth Ulhas Nair, Patricia Greninger, Xiaohu Zhang, Adam A. Friedman, Arnaud Amzallag, Eliane Cortez, Avinash Das Sahu, Joo Sang Lee, Anahita Dastur, Regina K. Egan, Ellen Murchie, Michele Ceribelli, Giovanna S. Crowther, Erin Beck, Joseph McClanaghan, Carleen Klump-Thomas, Jessica L. Boisvert, Leah J. Damon, Kelli M. Wilson, Jeffrey Ho, Angela Tam, Crystal McKnight, Sam Michael, Zina Itkin, Mathew J. Garnett, Jeffrey A. Engelman, Daniel A. Haber, Craig J. Thomas, Eytan Ruppin, and Cyril H. Benes. A landscape of response to drug combinations in non-small cell lung cancer. *Nature Communications*, 14(1):3830, June 2023. Number: 1 Publisher: Nature Publishing Group.

[5] Oliver Hantschel. Unexpected Off-Targets and Paradoxical Pathway Activation by Kinase Inhibitors. *ACS Chemical Biology*, 10(1):234–245, January 2015. Publisher: American Chemical Society.

[6] Elizabeth A. Coker, Adam Stewart, Bugra Ozer, Anna Minchom, Lisa Pickard, Ruth Ruddle, Suzanne Carreira, Sanjay Popat, Mary O’Brien, Florence Raynaud, Johann de Bono, Bissan Al-Lazikani, and Udai Banerji. Individualized Prediction of Drug Response and Rational Combination Therapy in NSCLC Using Artificial Intelligence–Enabled Studies of Acute Phosphoproteomic Changes. *Molecular Cancer Therapeutics*, 21(6):1020–1029, June 2022.

[7] Uri Ben-David, Benjamin Siranosian, Gavin Ha, Helen Tang, Yaara Oren, Kunihiko Hinohara, Craig A. Strathdee, Joshua Dempster, Nicholas J. Lyons, Robert Burns, Anwesha Nag, Guillaume Kugener, Beth Cimini, Peter Tsvetkov, Yosef E. Maruvka, Ryan O’Rourke, Anthony Garrity, Andrew A. Tubelli, Pratiti Bandopadhayay, Aviad Tsherniak, Francisca Vazquez, Bang Wong, Chet Birger, Mahmoud Ghandi, Aaron R. Thorner, Joshua A. Bittker, Matthew Meyerson, Gad Getz, Rameen Beroukhim, and Todd R. Golub. Genetic and transcriptional evolution alters cancer cell line drug response. *Nature*, 560(7718):325–330, August 2018.
